# Genetic architectures of postmating isolation and morphology of two highly diverged rockfishes (genus *Sebastes*)

**DOI:** 10.1101/2022.05.27.493803

**Authors:** Nozomu Muto, Takuma Kawasaki, Ryo Kakioka, Atsushi J. Nagano, Yuta Shimizu, Shu Inose, Yohei Shimizu, Hiroshi Takahashi

## Abstract

Postmating isolation is thought to be an important driver of the late stages of speciation. However, relatively little is empirically known about the process compared to other isolating mechanisms that drive the early stages of speciation, especially in non-model organisms. We characterized the genetic architecture of postmating isolation between two rockfishes, *Sebastes schlegelii* and *S. trivittatus*, whose reproductive isolation is complete. We examined transmission ratio distortion (TRD) patterns of genetic markers in two reciprocal backcross populations. Markers showing either of the two types of TRD was widespread across the genome, with some of the distorted markers forming extensive clusters around the recombination coldspots. These suggest that the postmating isolation effectively prevents gene flow across the genome and the recombination landscape contributes to the genetic architecture. Comparisons between two backcross families and two developmental stages showed little similarity in the distorted markers, suggesting asymmetry and stage-specificity of the isolation. This may be due to hybrid incompatibility involving maternal factors or extrinsic selection. The lack of sex-ratio distortion in the mapping families suggested that Haldane’s rule in terms of hybrid inviability does not hold. Additionally, QTL mapping detected significant QTLs for sex and the morphological traits relevant to speciation and convergence of rockfishes, including body coloration. Genes in the melanocortin system, including *agouti-signaling protein 1* (*asip1*) and *melanocortin 1 receptor* (*mc1r*), might underlie the horizontal and vertical color patterns on the body, respectively. These findings constitute an essential step towards a comprehensive understanding of speciation and morphological diversification of rockfishes.

## Introduction

Speciation is a continuous process whereby gene flow between diverging species diminishes with reproductive isolation. Existing species pairs can be ordered in a continuous sequence—a speciation continuum—according to the degree of divergence (Coyne & Orr, 1989a). Studying the species pairs at different points of the continuum enables the reconstruction of the order and the rate of evolution of different forms of reproductive barriers in a given system (Coyne & Orr, 1989a). A widely accepted view is that premating isolation due to local adaptation or sexual isolation tends to evolve early in speciation, whereas postmating isolation evolves later, thereby completing speciation or maintaining separate species (Coyne & Orr, 2004; Schemske, 2010). However, the latter process is relatively unclear, reflecting lesser attention being paid to highly diverged species pairs whose reproductive isolation is strong or complete (Matute & Cooper, 2021).

Postmating isolation may result from reduced fertilization success of heterospecific gametes (gametic isolation), intrinsic hybrid sterility and inviability (intrinsic postzygotic isolation), and environmentally-dependent hybrid sterility and inviability (extrinsic postzygotic isolation) (Coyne & Orr, 2004). Among them, intrinsic postzygotic isolation is the most intensively studied in terms of both its genetic model and contribution to speciation; little is known about the other two mechanisms (Rometsch et al., 2020). Intrinsic postzygotic isolation is caused by negative epistatic interactions involving two or more genes in hybrids, i.e., Bateson-Dobzhansky-Muller Incompatibility (BDMI) (Maheshwari & Barbash, 2011). Previous empirical studies have shown that the strength of intrinsic postzygotic isolation, as well as the number of loci involved in BDMI, often evolve exponentially with divergence (‘snowball effect’) (Matute et al., 2010; Stelkens et al., 2010), consistent with theoretical predictions (Orr & Turelli, 2001). Similarly, intrinsic postzygotic isolation characterizes the isolation of most ‘good species’ examined (Coyne & Orr, 2004). However, it is still largely unknown what factors account for the variations in tempo and mode of evolution of intrinsic postzygotic isolation across different systems and how its genetic architecture relates to its efficacy (Coughlan & Matute, 2020). Therefore, understanding how postmating isolation contributes to the later stages of speciation requires more studies on various groups of organisms with diverse life histories and degrees of divergence, especially the understudied non-model organisms, and on gametic and extrinsic postzygotic isolation.

Postmating isolation between a pair of species can be detected by analyzing the deviation of marker genotype frequencies from the Mendelian expectations (transmission ratio distortion, TRD) in hybrid mapping populations, such as backcross and F_2_ generations (Fishman & McIntosh, 2019). Gametic isolation and hybrid inviability between parental species may cause variations in the fertilization success of gametes and mortality of progeny in hybrid crosses, resulting in TRD at genetic markers linked to the underlying loci of these barriers. Importantly, hybrid inviability due to BDMI is largely masked in F_1_, but manifests in advanced generation hybrids (Orr, 1995; Orr & Turelli, 2001). Extrinsic hybrid inviability can also be inferred from the patterns of TRD relative to the rearing conditions. Moreover, mapping the distorted markers on a reference genome or a linkage map can reveal the genetic architecture of the isolating barriers (e.g., Brennan et al., 2014). Using TRD as a proxy of postmating isolation is particularly useful when direct fitness measurement is difficult. For example, gametic isolation and prenatal hybrid inviability in internally fertilizing animals could be inferred from TRD in the progeny immediately after parturition, which is otherwise, highly challenging.

Speciation and associated phenotypic diversification is sometimes accompanied by recurrent evolution of convergent traits (Schluter, 2000). Convergent traits between distant lineages often result from independent adaptations to similar ecological challenges, which in turn can lead to multiple independent speciation events underlain by a common premating isolating mechanism (Schluter, 2000). Whether and to what extent they share similar genetic mechanisms varies across systems (Rosenblum et al., 2014). The similarity in genetic mechanisms of convergence between distant lineages ranges from independent fixations or introgression of identical mutations to shared molecular pathways with different nucleotides and genes (Chen et al., 1997; Evans et al., 2012; Stern, 2013). The first critical step toward understanding the convergence at the genetic level is to characterize the genetic architecture of the convergent trait in each lineage.

Rockfishes of the genus *Sebastes* are an ideal system to study speciation and the associated phenotypic diversification in marine environments. *Sebastes* is a genus comprising more than 100 species of viviparous marine fishes, most of which diverged in the last 8 million years (Hyde & Vetter, 2007; Kolora et al., 2021). These species have colonized diverse habitats from the near-shore rocky reefs of less than 1 m depth to the deep-sea continental slopes of up to 1,500 m depth (Love et al., 2002; Nakabo & Kai, 2013). Numerous incipient species pairs hybridizing in nature are scattered across the phylogeny of the genus, enabling multiple independent studies on the earliest stage of speciation (Buonaccorsi et al., 2011; Muto et al., 2013; Saha et al., 2017; Schwenke et al., 2018). Convergence in some morphological traits among independent lineages has also been suggested either verbally or through phylogenetic comparative methods (Ingram & Kai, 2014; Kai et al., 2003), providing opportunities to study how those traits evolved along with the radiation in diverse environments. Furthermore, artificial crosses can produce fertile F_1_ hybrids even between species pairs with no record of natural hybrids (T. Kawasaki, unpublished data). This feature enables genetic dissections of postmating isolation and quantitative trait loci (QTL) analysis of morphological traits using advanced generation hybrids.

Nevertheless, relatively little is known about the speciation and morphological diversification of rockfishes. Incipient species often occupy habitats of different depths, implying the role of depth segregation in the early stages of speciation (Ingram, 2011; Longo et al., 2022; Muto et al., 2019; Shum et al., 2014; Venerus et al., 2013). Genomic regions presumably related to depth segregation have also been identified (Behrens et al., 2021; Heras & Aguilar, 2019). However, other possible isolating barriers have not been well studied. Most importantly, studies on isolating barriers between highly diverged species with strong reproductive isolation is lacking; thus, how reproductive barriers develop towards the completion of speciation in *Sebastes* remains elusive. Furthermore, some convergent morphological characteristics, such as body coloration, shape, and number of gill rakers, are potentially involved in premating isolation or niche partitioning within the genus (e.g., Ingram 2011), but its genetic bases have not yet been examined.

In this study, we primarily aimed to characterize genetic architecture of postmating isolation between the two rockfishes *S. trivittatus* and *S. schlegelii*, and additionally those of the morphological difference between the species and sex determination. They are estimated to have diverged 6–7 million years ago (mya) (Hyde & Vetter, 2007; Kolora et al., 2021), with no documented record of natural hybridization (Figure 1). They have partially overlapping geographical distributions in the western Pacific: *S. trivittatus* is distributed in higher latitudes, from the coasts of Sakhalin Island and the southern Kamchatka Peninsula to the Yellow Sea; whereas *S. schlegelii* is found from the coasts of Sakhalin and the southern Kuril Islands to the East China Sea (Nakabo & Kai, 2013; Parin et al., 2014) (Figure 1). Both species mainly inhabit near-shore rocky reefs, and it is unknown whether they are segregated by habitat depth or trophic niche. Their morphological difference involves several traits important in the contexts of speciation and convergence of rockfishes, such as body coloration. Using two reciprocal backcross families (single F_1_ male × females of *S. trivittatus* and *S. schlegelii*), we first constructed a hybrid linkage map. We then explored the TRD of genetic markers as an approximation of genetic architectures of postmating isolation. Comparisons of the patterns of TRD between two developmental stages and between the families revealed stage-specificity and asymmetry of the isolation. This study documents postmating isolation in *Sebastes* for the first time and signifies the first critical step for improved understanding of speciation and morphological diversification of the group. This will consequently provide a valuable contrast to those in well-studied model organisms.

**FIGURE 1.**
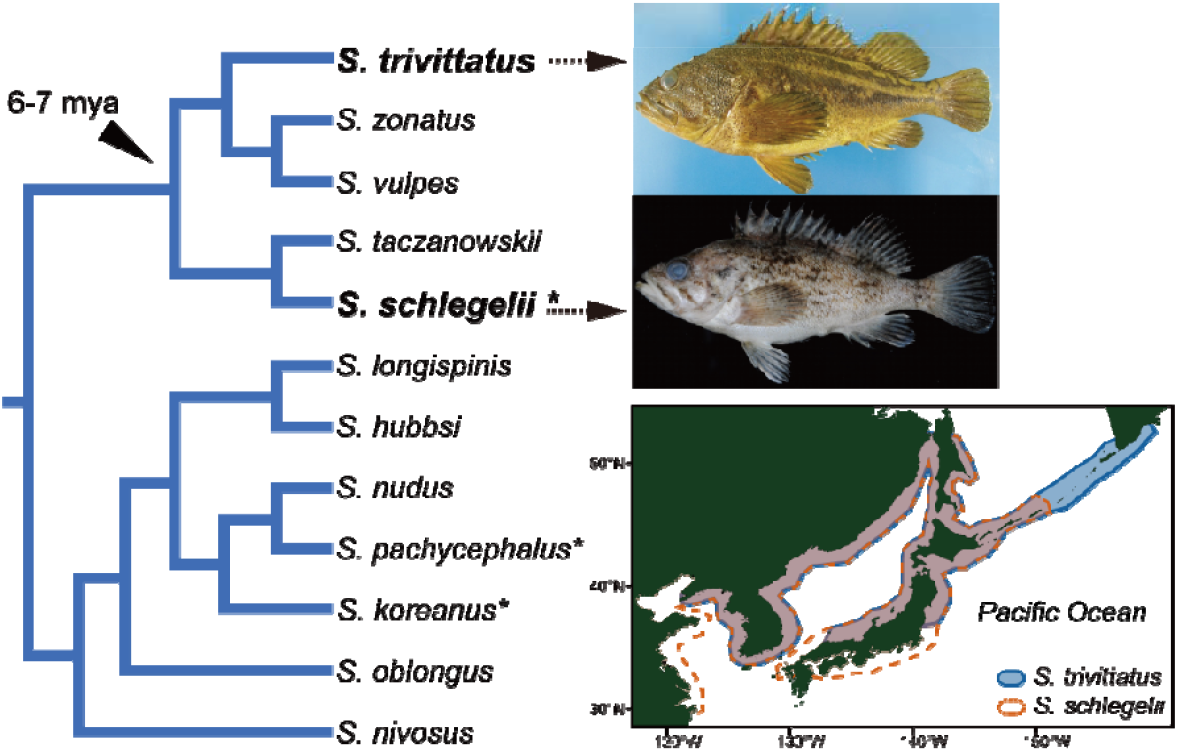
Phylogenetic relationship, overall appearance, and distribution ranges of *S. trivittatus* and *S. schlegelii*. Only the northwest Pacific subclade of the genus *Sebastes* is shown. Asterisks indicate the species for which sex determination by the *amhy* gene on chromosome 12 has been demonstrated. Images are of the specimens deposited in Kyoto University under the following registration numbers: *S. trivittatus*, FAKU 200925; *S. schlegelii*, FAKU 201558, kindly provided by Yoshiaki Kai. (120 mm width)

## Materials and methods

### Experimental crosses and sampling

Rockfishes are viviparous and have prolonged periods of sperm storage in female ovaries and gestation (each from one to several months) (Love et al., 2002). A female releases hundreds of thousands to millions of hatched larvae per spawning. Nutrients are supplied to the progeny both maternally (matrotrophy) and via yolk (Macfarlane & Bowers, 1995; Tengfei et al., 2021). The sperm storage and gestation periods of *S. schlegelii* are 4–6 and 1–2 months, respectively, whereas those of *S. trivittatus* are unknown (Sasaki, 2003). We artificially inseminated a wild female *S. trivittatus* in November 2012 using sperm obtained from a wild male *S. schlegelii*, both collected off Hokkaido, Japan. Parturition of the F_1_ yolk-sac larvae occurred ca. seven months from the insemination. Subsequently, a single male of the F_1_ progeny was crossed with a wild female *S. trivittatus* (different individual from the founder female) and a wild female *S. schlegelii* in December 2015. The parturitions occurred ca. six months later. The resulting backcross families are hereafter referred to as BC_Str and BC_Ssc, respectively. Artificial inseminations was performed following Kawasaki et al. (2017). Briefly, a sperm suspension was prepared by mincing testis and sperm reservoirs in urine obtained from male fish, which were euthanized with tricaine methanesulfonate (MS222). The suspension was then deposited into the external genital pore of the female fish. Rearing conditions for the fishes and a schematic diagram of the experimental design are provided in Appendix S1 and Figure S1. We sampled approximately 100 progeny of each family at each of the two stages: yolk-sac larvae (0 d after parturition) for genotyping and immature subadult (ca. 270 d: see Figure S1) for both phenotyping and genotyping. All the females used for the crosses had been separated from males at least for one year to prevent unintended mating. We could not use F_1_ females for the cross because female rockfishes naturally require a few years more than males for maturation (Love et al., 2002).

### Phenotyping

Independent lineages of *Sebastes* sometimes have strikingly convergent color patterns, including the presence of vertical bands on the dorsum, a horizontal stripe along the lateral line, and a horizontal stripe along the dorsal-fin base, but not necessarily similar colors. Both color and color patterns are often the most important trait to distinguish sister species of the genus (Ingram, 2011; Longo et al., 2022; Muto et al., 2019; Shum et al., 2014; Venerus et al., 2013), implying that visual mate recognition is one of key components of premating isolation in rockfishes. Convergence in meristic and morphometric traits involves gill-raker number and length, eye size, body depth and width, pectoral-fin length and width, upper- and lower-jaw lengths, and head spination (Ingram & Kai, 2014; Kai et al., 2003). Among these traits, eye size and body depth are associated with habitat depth, whereas number and length of gill rakers and body size are associated with trophic niche, implying that they are related to ecological isolation in rockfishes (Ingram, 2011). All of these traits that are potentially involved in convergence and/or premating isolation were phenotyped in subadults of the mapping families (Tables S1, S2). Additionally, we phenotyped other morphometric or meristic traits that often show relatively minor but significant differences between closely related species of rockfishes (e.g., Muto et al., 2011), thereby exploring a broader range of potentially relevant traits. Among the phenotyped traits, color-related traits, head spination, numbers of gill-rakers, lateral-line pores, and fin rays differ between *S. trivittatus* and *S. schlegelii*; however, the differences in other traits are unknown (Nakabo & Kai, 2013).

Therefore, for QTL mapping, we only used the traits for which significant differences were observed between the two backcross families, assuming that such differences represent the difference between the parental species. As for sex determination, the XY system controlled by the male master sex-determining gene *amhy* was demonstrated for *S. schlegelii, S. koreanus*, and *S. pachycephalus* (Song et al., 2021). The latter two species belong to the same subclade of *Sebastes* with *S. schlegelii* and *S. trivittatus* (Hyde & Vetter, 2007; Ingram & Kai, 2014)(Figure 1). The gene *amhy* is present in a ca. 5 kb male-specific region of the chromosome (CHR) 12 in the chromosome-level genome assembly of *S. schlegelii* (He et al., 2019), which we used as a reference genome in this study.

We measured standard length (SL) and body weight of each subadult fish and took a digital image of the left side of the body along with a KODAK Color Separation Guide using Pentax K200D immediately after euthanasia. We kept the distance between the camera and the fish constant across individuals. The sex of subadults was determined through visual inspections of the overall appearance of the fresh gonads: males have a pair of elongated, almost thread-like white gonads, whereas females have partly rounded orange gonads. The visual inspection was validated for a subset of individuals via histological observation. The fish was then fixed in 10% formalin and preserved in 70% ethanol.

The body color and the color patterns of each fish were characterized in three ways using the digital image (Figure S2). First, we coded the presence/absence of the vertical bands on the dorsum and the horizontal stripe along the lateral line and along the dorsal-fin base in each fish as binary traits based on visual inspection. The coding was done twice by the same person for consistency. Second, we quantified the overall intensities of black pigmentation (supposedly melanophore-based) and yellow pigmentation (xanthophore-based) on the body by counting the number of corresponding pixels. Each image was color-corrected by setting the grey patch of the color separation guide as the neutral grey using the eyedropper tool in the ‘Curves adjustment’ implemented in Adobe Photoshop CS6. We then cropped the image by a polygon defined by 11 landmarks on the body to restrict the analysis to an unambiguously definable area, excluding the fins, which were sometimes damaged. We counted the number of pixels on the cropped image that fell within the ranges of the hue/saturation/brightness (HSB) color space using the Color Threshold function of ImageJ 1.53c (Schneider et al., 2012); black: H = 0–255, S = 0–255, B = 0–100; yellow: H = 10–50, S = 0–255, B = 100–200. These ranges were determined by preliminary visual inspections of the color of the extracted pixels, of a subset of individuals, under variable combinations of HSB values. Third, we quantified the color patterns of the fish using the ‘patternize’ v0.0.2 package (Van Belleghem et al., 2018) on R v3.6.3 (R Core Team, 2020). We cropped the color-corrected digital image by a polygon defined by six landmarks on the body to restrict the analysis to the upper half of the flank, where the focal color patterns are expressed. We used the *patRegK* function to automatically align all images, assign pixel colors to a predefined number of clusters (K), and extract the color patterns from the image based on the distribution of each cluster. We analyzed with K = 4–6 (Figure S3). The variation in the extracted pattern was summarized by a Principal Component Analysis (PCA), performed for each color cluster of each K using the *patPCA* function. For QTL mapping, we used the PC scores derived from three color clusters from K = 5, which captured the color pattern variation most successfully (see Appendix S1 for more detail).

We quantified the overall body shape from the lateral view using geometric morphometrics. We used tpsDig2 (Rohlf, 2017) to digitize 16 landmarks on the digital image of each fish (Figure S2). Generalized Procrustes Analysis was performed on these landmarks using the *gpagen* function implemented in the R package ‘geomorph’ v4.0.0 (Adams et al., 2021). The variation in the obtained Procrustes shape coordinates among individuals was summarized by a PCA using the *gm*.*prcomp* function. Linear measurements and counts were made on 34 and 13 traits, respectively, on the left side of the body of the fixed samples. Methods for measuring and counting followed Muto et al. (2011, 2019) with the following additions: head spines, the number of spines on the dorsal aspect of the cranium; lacrimal spines, the number of spines on the lacrimal.

We log_10_ transformed SL, weight, all linear measurements, and intensities of pigmentations. We tested the dependence of each trait on size (SL) and sex in each family, and the difference between the two families, using either linear regression, the Wilcoxon’s rank-sum test, or the Fisher’s exact test, depending on the distribution of the trait (Tables S1, S2). The traits used for QTL mapping were tested for pairwise correlations using the *corr*.*test* function implemented in the R package ‘psych’ v2.1.6 (Revelle, 2021), with the false discover rate (FDR) correction. We used the residuals of linear regression of each trait on size or sex for the correlation test, if the trait was dependent on these variables.

### Genotyping

All parental individuals, 137 BC_Str individuals (48 larvae and 89 subadults) and 136 BC_Ssc individuals (48 larvae and 88 subadults) were subjected to genotyping by double-digest restriction-site-associated DNA sequencing (ddRAD-seq), according to Peterson et al. (2012) with minor modifications (see Appendix S1 for details). For subadults of BC_Str and their parents, a single library was prepared and sequenced with 50-bp single-end reads on an Illumina HiSeq 2500 system. A single library for the remaining individuals was prepared and sequenced with 150-bp paired-end reads on an illumina HiSeq X system. In either case, the sequencing was performed by Macrogen Japan Corporation (Kyoto, Japan). We used the two different sequencing platforms because the availability of related facilities changed during the study. We believe that the bias introduced by the combination of the two platforms is, if any, minimal because they both use the four-color chemistry. Only the forward sequences from the paired-end reads were used for consistency across the platforms. Parental individuals were sequenced twice or thrice for stringent genotyping.

We processed the ddRAD-seq reads using fastp v0.23.1 (Chen et al., 2018) to remove adapter sequences and trim the lengths to 50 bp. Reads with one or more bases with q-values < 20 were discarded. Two individuals of BC_Str subadults and an individual of BC_Ssc subadults were removed because of the few reads. The trimmed sequences were mapped to the chromosome-level reference genome assembly of a male *S. schlegelii* (He et al., 2019) (China National GeneBank Sequence Archive accession No. CNA0000824), using BWA-mem v0.7.17 (Li, 2013) with a mismatch penalty of 5. Unmapped reads and reads with a mapping quality < 5 were removed by Samtools v1.12 (Danecek et al., 2021). The resulting BAM files were used for identifying SNPs using ‘gstacks’ implemented in Stacks v2.5.4 (Catchen et al., 2013) with the default settings. The subsequent filtering steps were performed separately for each family to maximize the number of usable SNPs in the downstream analyses. We created a vcf file using ‘populations’ implemented in Stacks, retaining only the first SNP of a locus if two or more were detected. Genotypes with read depths < 10 or > 300 were removed, and SNPs were discarded if the resulting genotyping rate per SNP was < 0.6, using VCFtools v0.1.16 (Danecek et al., 2011). We then used SnpSift v4.3 (Cingolani et al., 2012) to retain only the SNPs for which the alleles observed in backcross progeny could be unambiguously determined to be inherited from either *S. trivittatus* or *S. schlegelii*. This was done by keeping only SNPs for which genotypes of grandparents and parents conformed to all the following criteria: 1) either homozygous in both grandparents or heterozygous in only one of the grandparents, 2) heterozygous in the F_1_ male parent, and 3) homozygous in the pure female parent. We then removed the SNPs genotyped in < 60 BC individuals.

### Linkage map and QTL

We first constructed a hybrid linkage map for each family using the R package ‘R/qtl’ (Arends et al., 2010) and then a consensus map of the two families using the R package ‘LPmerge’ v1.7 (Endelman & Plomion, 2014). The map for each family was constructed as follows. We removed markers showing deviation of genotype frequency from the expected even ratio of homozygotes and heterozygotes (hereafter referred to as single-locus TRD; chi-square test, *p* < 0.001 after FDR correction). Linkage groups (LGs) were initially formed with a minimum LOD score of 6 and a maximum recombination fraction (RF) of 0.35. The initial linkage grouping was highly consistent with the reference genome assembly of *S. schlegelii*, most markers on each LG being mapped to one of the 24 CHRs of the reference genome. However, deviations from this pattern were observed; a few markers on each LG mapped to an alternative (non-syntenic) CHR, and some markers formed an extra 25th LG. Such markers were either discarded, manually reassigned to another LG, or left unmodified based on examinations of individual LOD scores and RFs (Appendix S1; Figure S4; Table S3). Next, we ordered markers in each LG using the *orderMarkers* function. We checked the quality of the initial order by inspecting the plots of LOD score and RF between pairs of markers on each LG and corrected putative errors using either the *switch*.*order* or the *orderMarkers* function. We also sought better orders for LOD score and chromosome length using the *ripple* function. To identify problematic markers, we investigated the change in the length of each LG by eliminating one marker at a time using the *droponemarker* function. We removed the interior marker whose omission decreased the length by > 10 centimorgans (cM). We inferred genotyping errors using the *calc*.*errorlod* function and removed the genotypes with error LOD scores > 6. We also removed one outlier individual of each of BC_Str larvae and subadults concerning the numbers of crossovers. We then used the *ripple* function to search for marker orders with higher LOD scores. Finally, we compared the LOD score of the resulting marker order of each LG with that of the alternative order based on the physical position of markers on the corresponding CHR of the reference genome, and retained the order with a higher LOD score. If the order based on the physical position was chosen, we applied the *ripple* function again to search for orders with further higher LOD scores.

We performed QTL mapping separately for each family based on the consensus map using the R package ‘R/qtl2’ v0.24 (Broman et al., 2019). We used family-specific markers as well as markers shared by the two families. We first inserted pseudomarkers in the linkage map at intervals of 1 cM. We conducted a genome scan for each trait using the *scanone* function based on the Haley-Knott regression (Haley & Knott, 1992). We accounted for the effects of size (SL) or sex by using these variables as additive covariates if the focal trait was significantly associated with them. Genome scans for body weight were performed with or without size as a covariate to differentiate potential QTLs of growth and condition. We conducted 10,000 permutations to estimate the genome-wide significance threshold of LOD score at alpha = 0.05 using the *scan1perm* function. We estimated a 95% confidence interval (CI) for each QTL using the *bayes_int* function. We estimated the effect of a QTL by calculating the proportion of phenotypic variance explained (PVE): PVE = (sum of squares between genotypes) / (sum of squares total) (e.g., Takahashi et al., 2021). For the calculation of PVE and the visualization of phenotypic variation by genotype, we used the residuals of the regression of each trait on size or sex, if the trait was dependent on these variables; otherwise, we used raw phenotype values.

Genes possibly underlying the QTLs of color-related traits were searched for as follows. We first identified protein sequences of genes located in the regions of the reference genome of *S. schlegelii* that corresponded to the 95% CIs of the detected QTLs by processing the gene annotation of the reference genome using BEDOPS v2.4.41 (Neph et al., 2012). We then functionally annotated the identified genes using Pannzer2 (Törönen et al., 2018) and further extracted the genes with the terms “pigment” or “melan” in their gene ontology (GO) terms, following Gerwin et al. (2021). We performed blastp searches for the extracted genes to identify putative homologues in teleosts deposited in the NCBI RefSeq database. The QTLs of morphometric or meristic traits were not subjected to these procedures since filtering by GO terms is impractical for these traits.

### TRD

We tested two classes of TRD. First, we tested the single-locus TRD of each marker and assessed the distribution of the distorted markers on the reference genome. We used both the markers that were or were not filtered out during the construction of the linkage map for this analysis. Second, we tested non-random associations of genotypes between pairs of markers. In the present mapping families, the following two-marker genotypes are possible for a given pair of markers: StSt/StSt, StSt/StSc, StSc/StSt, and StSc/StSc for BC_Str; ScSc/ScSc, ScSc/StSc, StSc/ScSc, and StSc/StSc for BC_Ssc, with St and Sc representing alleles from *S. trivittatus* and *S. schlegelii*, respectively, and genotypes of two markers separated by a slash. Negative epistatic interaction (BDMI) between alleles at loci near the markers results in underrepresenting one or more of the two-marker genotypes, which is detectable as non-random associations of genotypes (Berdan et al., 2021; Pritchard et al., 2011). We calculated the expected two-marker genotype frequency from the observed genotype frequency of each marker. We then compared the observed and the expected two-marker genotype frequencies to detect non-random associations of genotypes using the chi-square test. For this analysis, we reduced the markers so that the minimum distance between adjacent markers became 1 cM using the *pickMarkerSubset* function in R/qtl. Consequently, a total of 47,694 tests among 316 markers and 59,580 tests among 353 markers were performed for BC_Str and BC_Ssc, respectively. Distributions of the pairs of markers showing significant non-random associations of genotypes were assessed on the linkage map rather than the reference genome since the analyzed markers had been filtered based on their positions on the linkage map. Both types of TRD were analyzed separately for larvae and subadults of each family and corrected for FDR using the *p*.*adjust* function in R.

## Results

### Linkage map

Genotype data after filtering consisted of 642 and 823 markers for BC_Str and BC_Ssc, respectively, with 545 markers common between them, and both families consisted of 135 individuals (87 subadults; 48 larvae). See Appendix S1 for more details on the sequencing and filtering. No marker was mapped to the unanchored sequences of the reference genome assembly. The hybrid linkage maps constructed separately for each family consisted of 24 LGs, consistent with karyotypes of both parental species as well as the reference genome (Figure 2) (Arai, 2011). The consensus map consisted of 898 markers, including 636 markers for BC_Str, 799 markers for BC_Ssc, and 537 shared markers, with a total length of 1413.5 cM and an average marker interval of 1.6 cM (Table S4). Each LG was syntenic to one of the 24 CHRs of the reference genome, showing high collinearity (Figure S5). Thus, the LGs were numbered according to the corresponding CHRs. Each LG had a ‘recombination coldspot’, where little or no recombination was observed across a wide range of the corresponding CHR of the reference genome (Figure S5). These regions were positioned distally on each LG except for that observed in the middle of LG 4, consistent with 23 acrocentric and one metacentric chromosome in both parental species (Ida et al., 1982).

**FIGURE 2.**
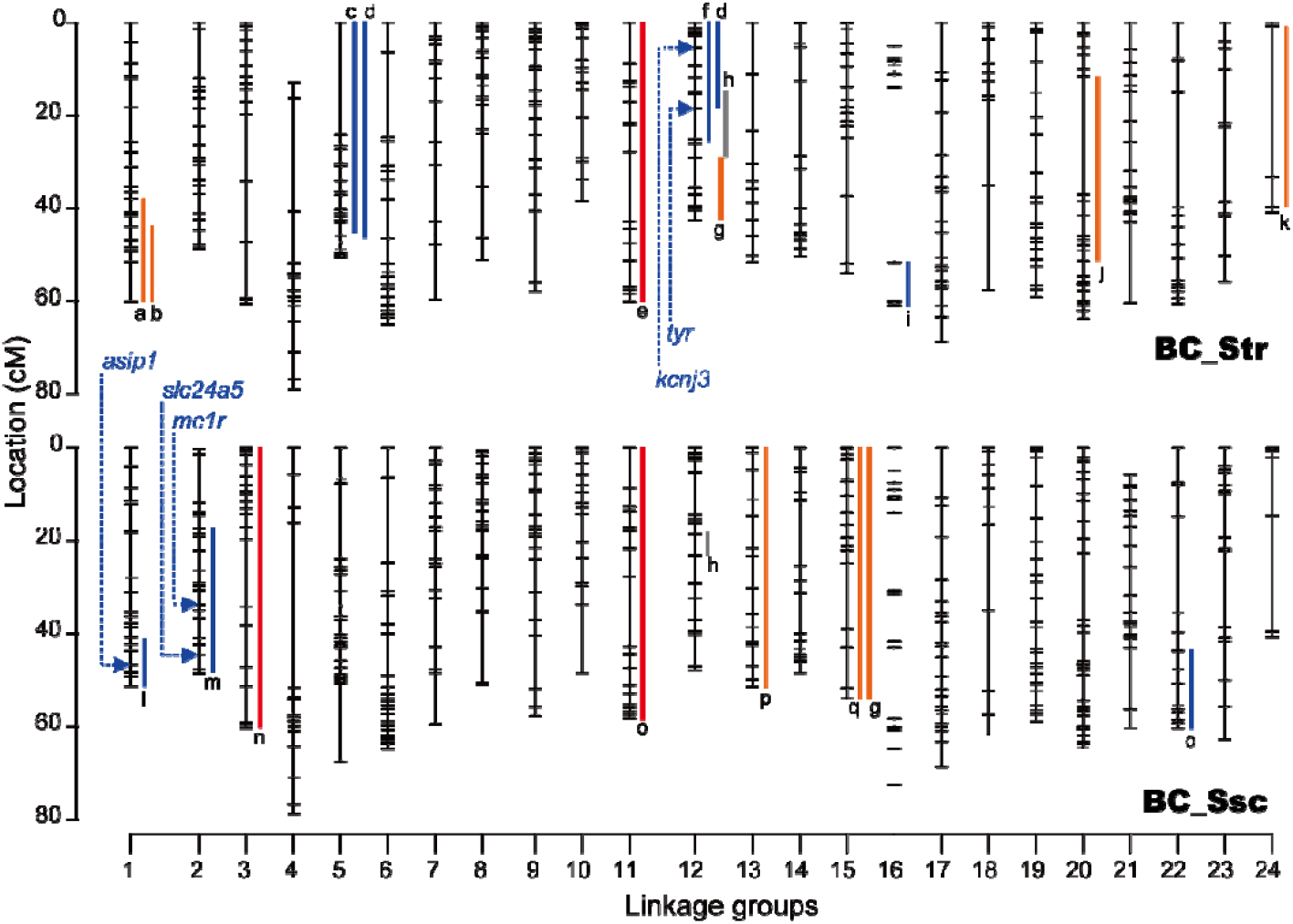
Consensus linkage maps and 95% confidence intervals of positions of QTLs for coloration (blue bars), shape (orange), counts (red), and sex (grey): a, PC2 scores based on geometric morphometrics; b, Anal-fin 2nd spine length; c, black pigmentation; d, yellow pigmentation; e, head spination; f, PC1 scores based on color cluster 1 derived from patternize analysis; g, lower peduncle length; h, sex; i, PC1 scores based on color cluster 3; j, predorsal length; k, anal-fin base length; l, stripe along lateral line; m, dark bands on dorsum; n, gill-rakers on lower arch; o, pectoral-fin rays; p, interorbital width; q, orbit diameter. Arrows indicate positions of the nearest markers of primary candidate genes of color-related traits discussed in the main text. The difference in the length of each linkage group between families is due to the presence of markers unique to either family (see also Table S4). (180 mm width)

### TRD

Distributions of genotype frequencies and single-locus TRD across the reference genome are shown in Figures 3, S6, and S7. Significant TRD at *p* < 0.05 after FDR correction was observed for a total of 131 markers, most of which showing TRD in only the subadults of either family (Figure 4; Table S5). In BC_Str, none of the distorted markers was common in both stages. In BC_Ssc, six distorted markers were shared between the stages, and one of them (‘scaffold6_37022538’) additionally showed TRD in the subadults of BC_Str. This marker, positioned terminally on CHR 6, showed an interesting pattern of distortion in which heterozygotes and homozygotes were overrepresented in BC_Str and BC_Ssc, respectively. Only 34.1% of the distorted markers of BC_Str subadults showed excesses of homozygotes. This largely reflects the presence of a few CHRs harboring extended regions with excesses of heterozygotes (see below). In contrast, 100%, 92.6%, and 83.3% of the distorted markers showed excesses of homozygotes in BC_Str larvae, BC_Ssc larvae, and BC_Ssc subadults, respectively.

**FIGURE 3.**
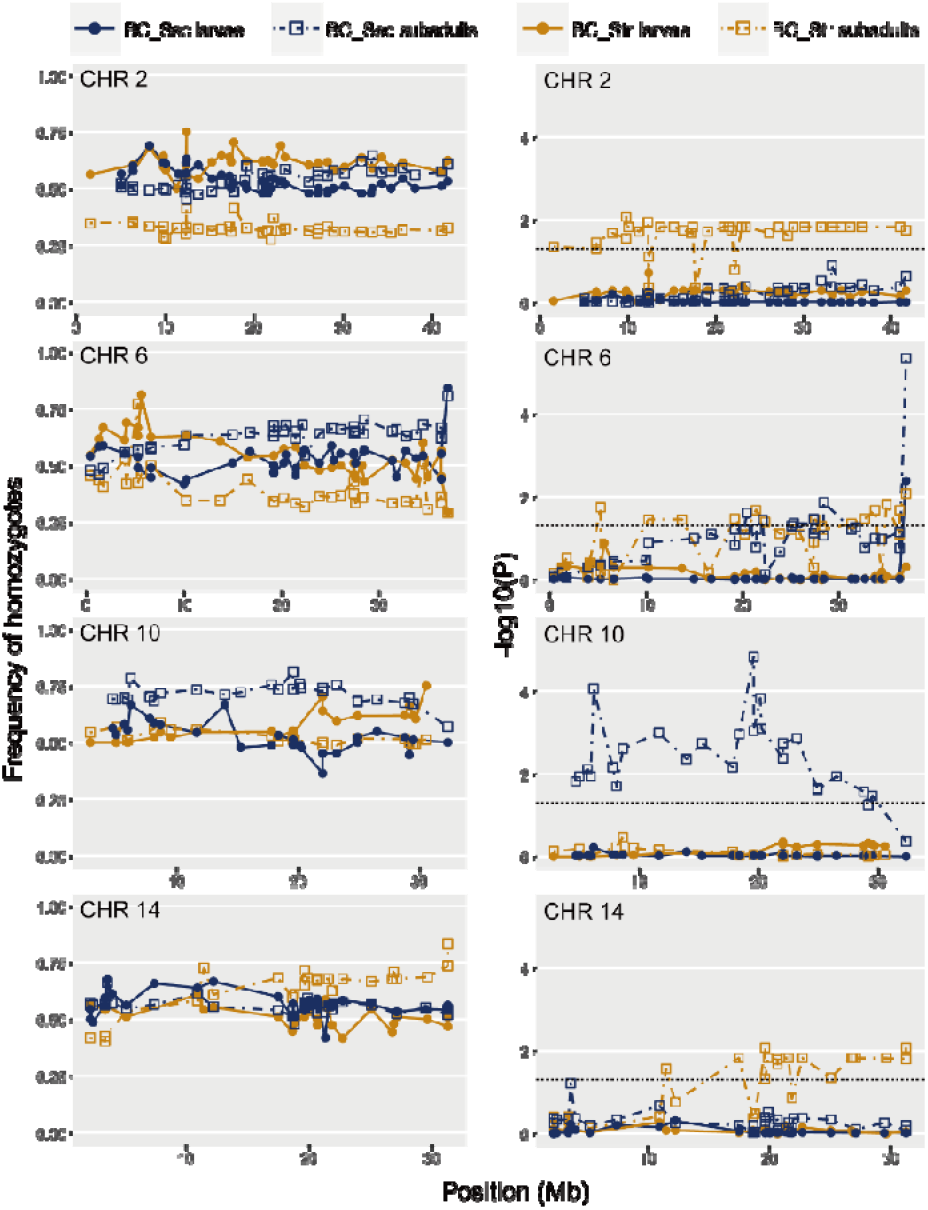
Genotype frequencies and transmission ratio distortion of markers on selected chromosomes, shown on the reference genome assembly of *S. schlegelii*. For each marker, frequencies of homozygotes (left column) and *p* values as −log_10_*p* from the chi-square tests for deviations of genotype frequencies from Mendelian expectations corrected for false discovery rates (FDR) (right column) are given. Dotted line indicates FDR-corrected *p* = 0.05. (135 mm width)

Several extended genomic regions, harboring more than ca. 20 markers with single-locus TRD in the same direction, were recorded (Figure 3). In BC_Str subadults, CHR 2 exhibited an excess of heterozygotes across its entire length. Approximately 27 Mb of CHR 6 showed an excess of heterozygotes in BC_Str and, in turn, an excess of homozygotes in BC_Ssc. CHRs 10 and 14 in BC_Ssc and BC_Str subadults, respectively, showed excesses of homozygotes across more than ca. 20 Mb. These regions on CHRs 6, 10, and 14 roughly corresponded to the recombination coldspots (Figure S5). Smaller clusters of 2–5 distorted markers spanning ca. 1.2–3.8 Mb were also observed in CHRs 7, 21, and 23 in BC_Str subadults and CHRs 4 and 19 in BC_Ssc subadults (Figures S6, S7). The remaining markers showing single-locus TRD were interspersed across the genome without forming a distinct cluster.

Tests for non-random associations of genotypes between pairs of markers on separate LGs yielded no significant result after FDR correction (all *p* > 0.42), partly because of numerous tests, making the tests highly conservative. We, therefore, focused on the marker pairs showing associations at nominal *p* < 0.01 and additional supports for the associations. The additional supports were the clustering of multiple marker pairs whereby two or more adjacent markers on a LG (hereafter referred to as a marker block) showed associations with another marker block on a different LG. Fifty five such associations (pairs of marker blocks) were found (Figure 5; Table S6); however, none of these was common across developmental stages or families, i.e., none of the associations observed in different stages/families involved marker blocks with overlapping ranges on both LGs. The numbers of the observed associations were comparable between the developmental stages in each family (Figure 4).

**FIGURE 4.**
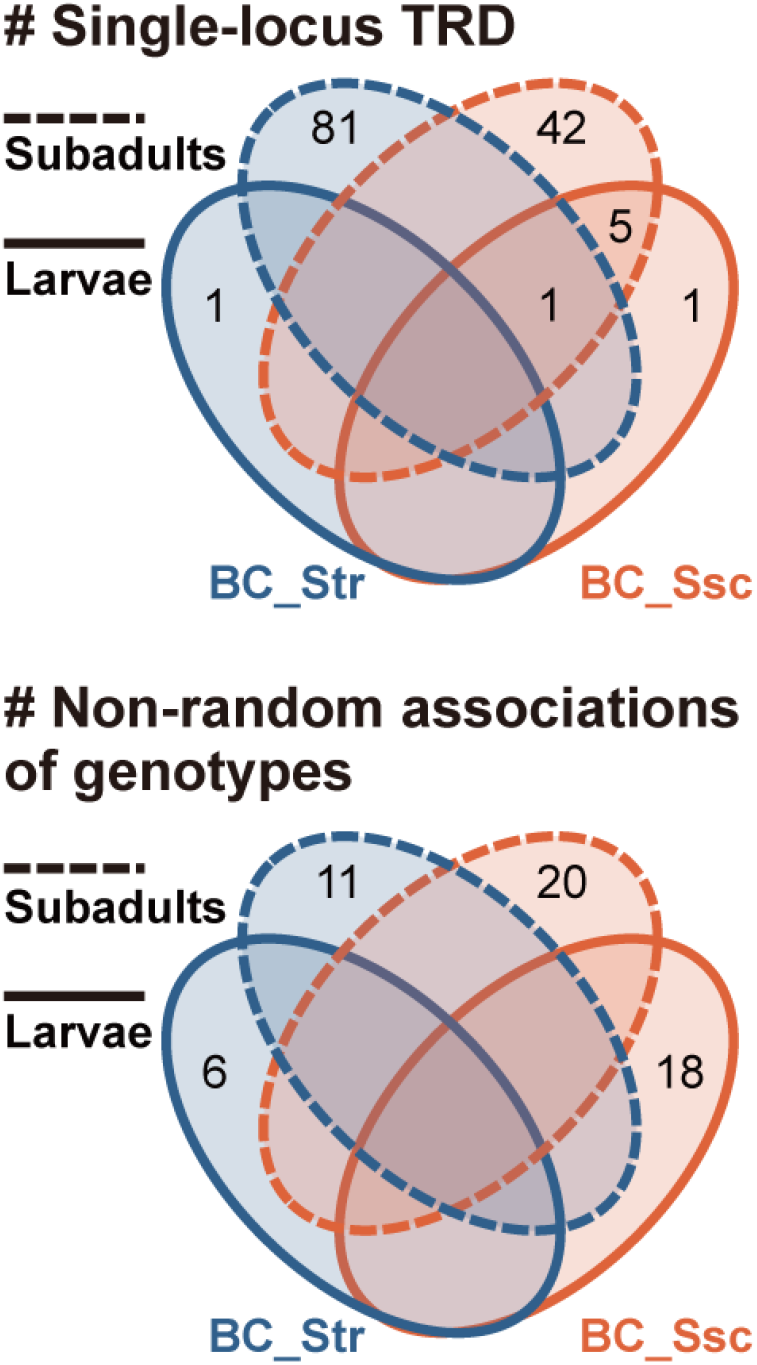
Venn diagrams for the number of markers showing significant single-locus transmission ratio distortion (TRD) and the number of pairs of marker blocks (groups of two or more adjacent markers) on separate linkage groups (LGs) showing non-random associations of genotypes. The non-random associations of genotypes between pairs of marker blocks were regarded as common across samples of different developmental stages/families if the associations in different samples involved marker blocks with overlapping ranges on both LGs (none observed in the present study). (80 mm width)

**FIGURE 5.**
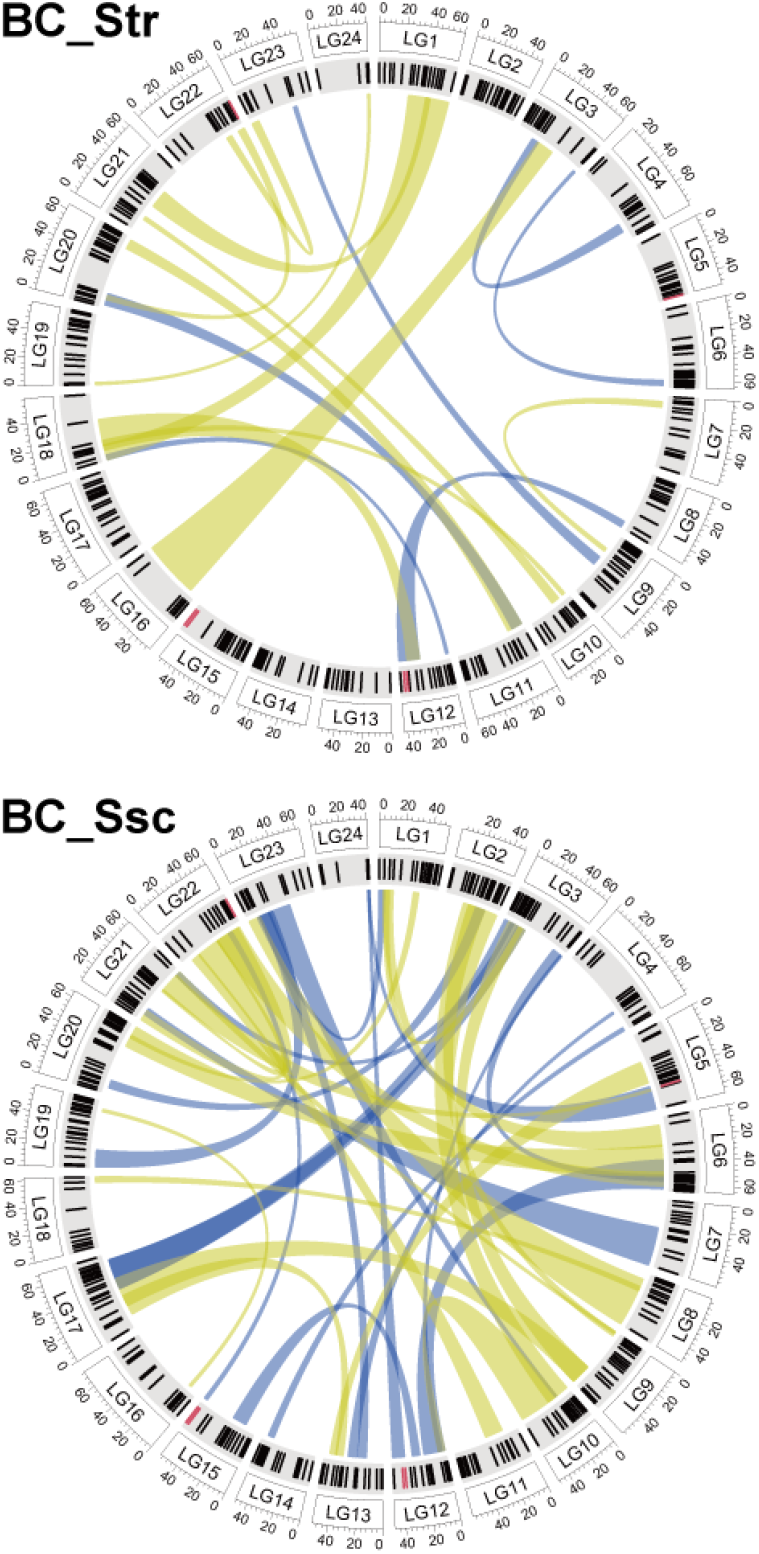
Possible epistatic interactions between pairs of markers on separate linkage groups (LGs). Each link represents a pair of marker blocks showing a non-random association of genotypes (chi-square test; nominal *p* < 0.01). Blue and yellow links correspond to the associations observed in larvae and subadults, respectively. The outer and inner circles represent each LG of the consensus linkage map and marker positions used for this analysis, respectively. Red ticks indicate the markers assigned to different (non-syntenic) LGs/chromosomes on the consensus linkage map and on the reference genome assembly of *S. schlegelii*. (80 mm width)

Sex ratio was slightly male-biased in subadults of BC_Str but did not differ significantly from parity (female:male = 0.40:0.60, n = 80; χ^2^ = 0.89, df = 1, *p* = 0.35), and was more even in BC_Ssc (female:male = 0.48:0.52, n = 95; χ^2^ = 0.09, df = 1, *p* = 0.75).

### QTL mapping

Thirty-nine morphological traits were subjected to QTL mapping, showing significant differences between the two families (Figure S8; Tables S1, S2). Significant QTLs with LOD scores greater than the genome-wide thresholds were detected for 11 and 9 traits in BC_Str and BC_Ssc, respectively (Figure 2; Table 1; Figures S9, S10). Single significant QTL for each trait was detected, except for yellow pigmentations of BC_Str, where two QTLs were detected on different LGs. The 1st and the 3rd color clusters of the patternize analysis represented the variations in a stripe along the lateral line and vertical bands on the dorsum, respectively (Figure S3). PC2 scores based on the geometric morphometrics represented variations in the relative positions of the head and body (Figure S8). Directions of the QTLs were consistent with the differences between the parental species except for the following traits: predorsal length and the 3rd color cluster of the patternize analysis in BC_Str; lower caudal-peduncle length and yellow pigmentations in BC_Ssc. The QTL for a horizontal stripe along the lateral line exhibited the highest PVE among all morphological traits. In addition, QTL for sex with a high LOD score was detected on LG 12 in each family, with the position roughly corresponding to that of the *amhy* gene: 26.4 Mb on the reference genome (Song et al., 2021), ca. 25 cM on the consensus map (Figure S5).

**TABLE 1.** Significant QTLs for body coloration, shape, counts, and sex. The ‘Direction’ column indicates whether *S. trivittatus* (Str) or *S. schlegelii* (Ssc) alleles cause higher trait values (for continuous traits) or the presence of the focal trait (for binary traits). Asterisks indicate allelic effects in the direction expected from the divergence between the parental species.

| Trait | Family | LG | Pos (cM) | Pos (Mb)<sup>a</sup> | LOD | CI lower | CI upper | PVE | Direction |
| --- | --- | --- | --- | --- | --- | --- | --- | --- | --- |
| Coloration |  |  |  |  |  |  |  |  |  |
| Color cluster 1: PC1 | BC_Str | 12 | 0.0 | 0.7 | 3.00 | 0.0 | 26.2 | 14.6 | Ssc* |
| Color cluster 3: PC1 | BC_Str | 16 | 60.8 | 25.8 | 3.36 | 51.6 | 60.8 | 16.6 | Str |
| Black pigmentation | BC_Str | 5 | 11.0 | 1.3 | 2.77 | 0.0 | 45.9 | 6.2 | Ssc* |
| Yellow pigmentation | BC_Str | 5 | 34.1 | 11.4 | 3.21 | 0.0 | 47.4 | 12.2 | Str* |
| Yellow pigmentation | BC_Str | 12 | 15.0 | 16.7 | 3.31 | 0.0 | 18.5 | 11.8 | Str* |
| Yellow pigmentation | BC_Ssc | 22 | 58.0 | 23.4 | 5.86 | 43.9 | 60.5 | 22.9 | Ssc |
| Stripe along lateral line | BC_Ssc | 1 | 51.0 | 29.6 | 9.27 | 40.8 | 51.5 | 46.9 | Str* |
| Dark bands on dorsum | BC_Ssc | 2 | 35.3 | 23.0 | 3.41 | 17.5 | 48.7 | 19.0 | Str* |
| Shape |  |  |  |  |  |  |  |  |  |
| GM: PC2 | BC_Str | 1 | 40.8 | 27.2 | 3.05 | 37.8 | 60.2 | 15.8 | Str* |
| Orbit diameter | BC_Ssc | 15 | 6.0 | 12.9 | 3.25 | 0.0 | 54.0 | 6.4 | Ssc* |
| Interorbital width | BC_Ssc | 13 | 2.0 | 2.7 | 3.75 | 0.0 | 51.6 | 4.2 | Ssc* |
| Lower peduncle length | BC_Str | 12 | 35.6 | 32.8 | 3.19 | 29.2 | 42.5 | 5.7 | Ssc* |
| Lower peduncle length | BC_Ssc | 15 | 24.9 | 23.9 | 3.04 | 0.0 | 54.0 | 5.0 | Str |
| Predorsal length | BC_Str | 20 | 37.0 | 6.8 | 2.70 | 11.5 | 51.3 | 2.2 | Ssc |
| Anal-fin base length | BC_Str | 24 | 24.0 | 12.1 | 2.51 | 0.8 | 39.6 | 17.4 | Str* |
| Anal-fin 2nd spine length | BC_Str | 1 | 51.0 | 29.6 | 4.84 | 43.9 | 60.2 | 10.5 | Str* |
| Counts |  |  |  |  |  |  |  |  |  |
| Gill rakers on lower arch | BC_Ssc | 3 | 23.0 | 32.3 | 2.78 | 0.0 | 60.5 | 15.5 | Str* |
| Pectoral-fin rays | BC_Ssc | 11 | 37.0 | 17.5 | 2.79 | 0.0 | 58.3 | 14.7 | Ssc* |
| Spines on head | BC_Str | 11 | 54.0 | 21.5 | 3.42 | 0.0 | 60.2 | 16.7 | Str* |
| Sex | BC_Str | 12 | 25.2 | 26.1 | 7.71 | 15.4 | 29.2 | 49.6 | NA |
| Sex | BC_Ssc | 12 | 21.0 | 27.6 | 20.80 | 18.5 | 23.2 | 94.9 | NA |

<sup>a</sup>Based on the marker position on the reference genome assembly of *S. schlegelii*. For the QTL on pseudomarker, the physical position of the nearest marker is provided.
Abbreviations: CI, confidence interval; cM: centimorgan; GM, geometric morphometrics; LG, linkage group; Mb, megabase; PVE, per cent variance explained.

We found 3–12 candidate genes for each of the QTLs of four color-related traits (Table S7). These included the following genes that have previously been associated with body coloration in other teleosts (see Discussion): agouti signaling protein 1 (*asip1*) for the horizontal stripe along the lateral line, solute carrier family 24 member 5 (*slc24a5*) and melanocortin 1 receptor (*mc1r*) for the vertical bands on the dorsum, tyrosinase (*tyr*) and potassium inwardly rectifying channel subfamily J member 13 (*kcnj13*) for PC1 scores of the patternize analysis, and *kcnj13* for the intensity of yellow pigmentations (Figure 2). Candidate genes were not searched for the other color-related QTLs since their 95% CIs were as wide as the lengths of the mapped LGs or their directions were inconsistent with the differences between the parental species.

## Discussion

### General patterns of TRD and postmating isolation

We observed numerous markers showing two classes of TRD: single-locus TRD and non-random associations of genotypes between pairs of markers. These markers were presumably linked to the loci that conferred differential fertilization success of gametes or selective mortality of hybrid progeny, and hence, the barrier loci underlying the postmating isolation between *S. trivittatus* and *S. schlegelii*. Importantly, these markers were widely distributed across the genome and some of them encompassed large proportions of particular chromosomes. Such a genetic architecture suggests genetic redundancy and the multifarious nature of the postmating isolation. These features would strongly reduce gene flow across the genome if the two species did mate, thereby enabling the postmating isolation to play a critical role in completing speciation or maintaining distinct gene pools (Coughlan & Matute, 2020; Nosil et al., 2009). Three of the four extensive genomic regions with single-locus TRD in subadults of one or both families coincided with the recombination coldspots. This suggests that each of these three regions represents the isolating effects of one or a few barrier loci affecting broad surrounding regions due to the tight linkage, rather than the effects of more numerous and widespread barrier loci. Overall, the combined effects of the recombination landscape and the effects of individual barrier loci created the observed complex genetic architecture.

TRD was observed in each of the four samples differing in the developmental stages and the directions of the crosses, with only a few distorted loci shared between them (Figure 4). TRD observed in larvae can be ascribed to postmating isolating barriers acting at the prenatal stage, from the artificial insemination until the parturition, including gametic isolation and hybrid inviability. TRD in subadults can be based on hybrid inviability at the postnatal stage from the parturition until sacrifice. In fact, the distinct genetic architectures observed for larvae and subadults suggest that largely different isolating mechanisms have been acting at the pre and postnatal stages. On the other hand, asymmetric isolation has been frequently reported in previous studies employing reciprocal interspecific crosses, and its primary cause is thought to be hybrid incompatibility involving uniparentally inherited genetic factors or maternal effects (Turelli & Moyle, 2007). More discussion on specific mechanisms of TRD in different samples is provided in section 4.1.2.

We found no significant bias in the sex ratio in either family, suggesting little or no difference in survival rates between sexes. We also observed no tendency for markers on the sex chromosome CHR 12 to be more frequently involved in either type of TRD. These results suggest that, regarding hybrid inviability, the present system does not obey the two ‘rules’ concerning reproductive isolation: disproportionate contribution of X or Z chromosomes to postmating isolation (large X effect) and more severe fitness reduction of heterogametic sex in hybrid crosses (Haldane’s rule) (Coyne & Orr, 1989b). This observation is consistent with homomorphic, recombining sex chromosomes in *Sebastes* (Arai, 2011; Song et al., 2021; this study), since empirical support for the two rules mostly came from organisms with heteromorphic sex chromosomes, around which the relevant theories have been built (Coyne, 2018). Teleosts generally do not obey Haldane’s rule regarding hybrid inviability (reviewed in Schilthuizen et al., 2011); the present system is considered one of such cases.

We found discordance between the current linkage map and the reference genome of *S. schlegelii* regarding the assignments of some markers to LGs/CHRs (Figure S5). Some of the markers also showed non-random associations of genotypes with markers on another LG (Figure 5). These results are reminiscent of reciprocal translocations between the parental species. However, reciprocal translocation in parental species generally has more extensive effects on a hybrid linkage map whereby LGs involved in the translocation are connected into a single LG via markers on the translocated region (Farré et al., 2011), which was not observed herein (Figure S4). We suspect that this discordance resulted from slight errors in our linkage map or the genome assembly. Further studies will be necessary to clarify whether the discordance represents some form of chromosomal rearrangements, an important mechanism of reproductive isolation and adaptive divergence between taxa (e.g., Matschiner et al., 2022).

A previous genomic study by Behrens et al. (2021) has identified several genomic regions showing elevated values of both *F*_ST_ and *D*_*XY*_ between each of the two pairs of incipient, depth-segregated species of *Sebastes* (*S. carnatus*–*S. chrysomelas* and *S. miniatus*–”*S. crocotulus*”), both pairs belonging to different subclades from the present parental species. These regions are supposedly involved in adaptive divergence and reproductive isolation in the presence of gene flow in each pair (“genomic islands of divergence”). The regions were located at 38.5–39.0 Mb on CHR 3, 18.5–20.0 Mb on CHR 4, 0.5–1.5 Mb on CHR 7, 10.5–11.0 Mb on CHR 9, and 30.0–32.5 Mb on CHR 13 on the same reference genome as that used in the present study. None of these regions overlaps with the extended TRD possibly associated with extrinsic selection in the present system (see below). Instead, these regions except that on CHR 3 coincided with the recombination coldspots (Figure S5), implicating conserved recombination landscapes across the phylogeny of rockfishes and their role in determining the genetic architecture of reproductive isolation (Burri et al., 2015; Nachman & Payseur, 2012).

Our findings suggested that incomplete postmating isolation acting at advanced generation hybrids (backcross, F_2_, and so forth) can develop within 10 million years in rockfishes. The pace could be comparable to that reported for F_2_ crosses of African cichlids (Stelkens et al., 2015). Contrastingly, we are currently unable to make any assumption regarding the evolution of postmating isolation acting at F_1_ hybrids, because the isolation was not formally examined here. We did not observe any signs of reduced survival rate (such as mass mortality) in the F_1_ progeny between *S. schlegelii* and *S. trivittatus* at least after the juvenile stage. This is also true for other combinations of F_1_ crosses within the northwest Pacific subclade of rockfishes, such as between *S. vulpes* and *S. schlegelii* (T. Kawasaki, personal observations). Instead, these F_1_ hybrids seem to be more resilient to harsh and fluctuating rearing conditions, reminiscent of heterosis. Nonetheless, they could in fact have suffered from an unnoticeable reduction in survival during their earlier life stages or fertilization failure. Postmating isolation, or more specifically, intrinsic postzygotic isolation, is generally stronger in advanced generation hybrids in which recessive BDMI is expressed (hybrid breakdown), whereas most previous studies on the evolution of postmating isolation focused on F_1_ hybrids (e.g., Coyne & Orr, 1989a; Stelkens et al., 2010).

### Possible mechanisms of isolation

Owing to the present cross design and the distinct (asymmetric) genetic architectures of the two families, we suggest that he most likely isolating mechanisms causing TRD at the prenatal stage are gametic isolation and hybrid inviability involving maternal factors. Under the present cross design, both F_1_ sperm and embryos of the two families have been stored in ovaries of reciprocal maternal species during sperm storage and gestation, with the embryos possessing maternally inherited genetic factors from their respective maternal species (Figure S1). BDMI involving maternally inherited factors such as mito-nuclear incompatibility is a predominant mechanism of asymmetric reproductive isolation in reciprocal hybrid crosses (Turelli & Moyle, 2007). Empirical evidence that mitochondrial genomes from reciprocal parental populations cause TRD in distinct sets of chromosomes in reciprocal F_2_ crosses has been reported for marine copepod (Lima et al., 2019). Besides, interactions between maternally expressed factors and sperm or zygotes within the mother’s body can result in asymmetric reproductive isolation (e.g., Devigili et al., 2018). Notably, such interactions are predicted to be driven by mother-offspring conflict in viviparous, matrotrophic animals, including rockfishes (viviparity-driven conflict hypothesis; Zeh & Zeh, 2000). In any case, the mechanisms of the TRD at the prenatal stage might be genetically complex, involving epistatic interactions among multiple nuclear loci, as well as the interactions with maternally inherited or expressed factors.

The TRD at the postnatal stage may also be explained by multiple factors, including intrinsic hybrid inviability due to BDMI involving maternally inherited factors and environmentally-dependent (extrinsic) inviability. The former can have a common mechanistic and genetic basis with that acting at the prenatal stage. For instance, mito-nuclear incompatibility may disrupt the metabolic function of mitochondria essential to all developmental stages (Burton & Barreto, 2012). In fact, five loci showing significant excess of homozygotes on CHRs 1, 4, 7, and 13 were shared between the two stages in BC_Ssc. These may represent loci that confer fitness effects across the developmental stages via intrinsic asymmetric hybrid inviability. The remaining distorted loci that were not shared with the prenatal stage are considered to have been caused by stage-specific intrinsic hybrid inviability or extrinsic inviability.

Extrinsic hybrid inviability occurs when ecologically divergent species produce hybrids with intermediate phenotypes, and the hybrid suffers reduced fitness in the parental environments (Nosil, 2012). This can be facilitated by mito-nuclear incompatibility (Wang et al., 2021). The present study was not specifically designed to examine this process. Nonetheless, ca. 20 Mb region of CHR 6 showing heterozygote excess in BC_Str and homozygote excess in BC_Ssc suggest additive selection against *S. trivittatus* alleles, implying extrinsic hybrid inviability under an environment where *S. trivittatus* is less fit than *S. schlegelii* (Rundle & Whitlock, 2001). The pattern also implicates immigrant inviability, a class of premating isolation whereby immigrants of parental species suffer reduced fitness in their non-native habitats (Nosil, 2012). Additionally, regions with homozygote excess in BC_Ssc subadults (e.g., CHR 10) or heterozygote excess in BC_Str subadults (e.g., CHR 2) with no concurrent TRD in the other family could also be associated with extrinsic selection against *S. trivittatus* alleles with dominance effects. A possible environmental factor underlying the extrinsic selection is water temperature. Species distribution models for *S. schlegelii* have suggested that water temperature is the most important environmental variable determining the natural distribution of the species, and it prefers water temperatures of 3–20 ºC (Chen et al., 2021). Although the thermal tolerance of *S. trivittatus* is unknown, it is reasonable to assume that this species is adapted to colder waters than *S. schlegelii*, given the more northern distributional of the former. On the other hand, the water temperatures of the tanks were 3.1–21.3 ºC in the present study (see Appendix S1). These temperatures are within the range tolerated by *S. schlegelii* but may have deviated from that by *S. trivittatus*, thus, selecting against *S. trivittatus* alleles at relevant loci.

The terminal locus (‘scaffold6_37022538’) of the distorted region of CHR 6 additionally showed TRD in larvae, although the distortion in BC_Str larvae was not statistically significant (Figure 3; Table S5). The directions of the distortion were the same between stages in each family (heterozygote excess in BC_Str; homozygote excess in BC_Ssc), suggesting that the extrinsic selection may have acted at the prenatal stage as well. Effects of ambient thermal conditions on prenatal viability of offspring via mother’s body temperature of viviparous snake have been previously documented (Lourdais et al., 2004).

### QTLs for morphological traits and sex determination

Among the morphological traits with significant QTLs detected, the following traits have been previously suggested to be related to convergence and/or speciation of rockfishes: orbit diameter, number of gill-rakers on the lower arch, head spination, intensities of pigmentations, stripe along the lateral line, and dark bands on the dorsum. The color clusters based on the patternize analysis are also considered among such traits since these are the quantitative representations of the color patterns. The lack of concordance between the two crosses should be treated with caution; although this may be explained by the dominance of the QTLs, the relatively small sample sizes or alleles segregating within the species may also be responsible. Nonetheless, the present study constitutes the first step toward understanding the genetic bases of the traits at the individual gene level, which will, in turn, provide insights into the traits’ evolutionary significance and histories within and across lineages (e.g., Kokita et al., 2021; Kratochwil et al., 2018). Below we discuss the results for the color-related traits whose candidate genes could be identified, focusing on the genes that have previously been associated with colorations in other teleosts. Subsequently, we briefly discuss the QTLs for other morphological characters and sex determination.

Horizontal stripes and vertical bands on the body are the most common color patterns in teleosts, and their genetic bases have been investigated in a handful of model systems. In African cichlids, regulatory changes of Agouti-related peptide 2 (*agrp2*), a member of the Agouti gene family, is associated with variation in horizontal stripes, facilitating convergence across the radiations (Kratochwil et al., 2018; Urban et al., 2021). We instead identified another member of the Agouti gene family, *asip1*, as a candidate gene for the horizontal stripe in the present system. Members of the Agouti gene family are part of the melanocortin system and act as antagonists of melanocortin receptors; both Asip1 and Agrp2 mainly antagonize one of five melanocortin receptors, Mc1r (Cal et al., 2017; Zhang et al., 2010). The convergence of the horizontal stripe in rockfishes and African cichlids may have different genetic bases (*asip1* vs. *agrp2*) but a common downstream molecular pathway (melanocortin system, via Mc1r). Activation of Mc1r induces cAMP production, which eventually leads to melanin synthesis (Cal et al., 2017). Mc1r is also involved in melanophore proliferation and melanosome dispersion (Singh & Nüsslein-Volhard, 2015). Any of these three processes can be involved in color patterning.

The QTL for PC 1 scores based on the 1st color cluster of the patternize analysis was located on LG 12. Since the scores primarily represented the horizontal stripe along the lateral line (Figure S3), the results suggest that the horizontal stripe in the present system is associated with multiple genomic regions. Interestingly, the QTL on LG 12 included a candidate gene *tyr*, which encodes a key enzyme of melanin synthesis activated by Mc1r via cAMP (Cal et al., 2017). The horizontal stripe in rockfishes thus might be controlled by changes in multiple genes in the melanin synthesis pathway modulated by the melanocortin system. In addition, this QTL for PC 1 scores of the patternize analysis partially overlapped with that of the intensity of yellow pigmentations, and the overlapping region encompassed *kcnj13*, which encodes a potassium channel conserved across vertebrates. Such possible associations of *kcnj13* with both the intensities of yellow pigmentations (supposedly representing xanthophores) and the color pattern is consistent with the known features of the gene. The potassium channel encoded by *kcnj13* is involved in the repulsion between melanophores and xanthophores during color pattern formation in zebrafish *Danio rerio*, and the divergence in *kcnj13* may have caused the color pattern diversification in the genus *Danio* (Podobnik et al., 2020).

For the vertical bands, we identified *mc1r* as a candidate gene, suggesting that the melanocortin system is associated with the vertical as well as horizontal color patterns in rockfishes. Alternatively, another candidate gene, *slc24a5*, may be a key component of the formation of the vertical bands. The gene *slc24a5* encodes a cation exchanger that is essential to melanin synthesis (Lamason et al., 2005), and it was included in one of several QTLs for vertical bars detected in an F_2_ cross between African cichlids *Pseudotropheus cyaneorhabdos* and *Chindongo demasoni* (Gerwin et al., 2021). Moreover, the upregulation of *slc24a5* (but not *mc1r*) was associated with the formation of vertical bars in an African cichlid *Haplochromis latifasciatus* (Liang et al., 2020).

Rockfish species occupying different habitat depths tend to have different eye size presumably due to divergent selection from contrasting light environments, and the eye-size differentiation tended to occur simultaneously with parapatric speciation by depth segregation throughout the phylogeny of the genus (Ingram 2011). Contrastingly, number of gill-rakers tends to be diverged within co-occurring assemblages of rockfishes, enabling resource partitioning, but the divergence did not tend to take place at speciation (Ingram & Shurin 2009; Ingram 2011). Interestingly, the significant QTL for the number of gill-rakers on LG 3, rather than that of orbit diameter, overlapped with one of the genomic islands of divergence between depth-segregated *S. miniatus* and “*S. crocotulus*” (Behrens et al., 2021; see the preceding section on the general pattern of TRD). This pattern implies that the maintenance of their species boundary is in part based on trophic divergence that has evolved after parapatric speciation. This hypothesis can be tested by further narrowing down the currently broad CIs of the QTL (Fig. 2) and examinations on the morphological and trophic niche differentiations between *S. miniatus* and “*S. crocotulus*”. Finally, it should also be noted that there may be unnoticed concordance between the genetic architecture of the morphological divergence and the genomic islands of divergence; we have detected only one significant QTL for each character, although those complex morphological characters should be polygenic.

The present result implicated a common sex-determination system for *S. schlegelii* and *S. trivittatus*, i.e., male-determination by *amhy*. The role of *amhy* has been originally demonstrated for *S. schlegelii* and additionally confirmed in *S. pachycephalus*, and *S. koreanus* (Song et al., 2021). Under the present cross design, *amhy* inherited from the male F_0_ *S. schlegelii* segregated in each backcross family. Besides, a highly significant QTL for sex was detected on LG 12 in each family, with CIs encompassing the presumed position of *amhy*. Taken together, *amhy* from F_0_ *S. schlegelii* successfully determined sex in both families despite the contrasting degrees of genetic contributions from *S. schlegelii* and *S. trivittatus*, implying that the sex of *S. trivittatus* is also controlled by this gene. The sex-determination role of *amhy* might be conserved across the members of the northwest Pacific subclade of the genus, to which the aforementioned four species belong (Figure 1).

### Concluding remarks

We provided a snapshot of postmating isolation in the speciation continuum of rockfishes at the point where reproductive isolation is complete in nature. It is most likely characterized by 1) the involvement of tens of loci across the genome, 2) asymmetry of the underlying loci concerning the direction of the crosses, and 3) multifarious mechanisms including asymmetric gametic isolation and hybrid inviability due to interaction with maternal factors, and extrinsic selection. Future studies employing direct observations of the isolating mechanisms at the phenotypic level, quantifications of the strength of the isolating mechanisms, and comparisons with other pairs of species at various points in the speciation continuum, will contribute to a more comprehensive understanding of the speciation in rockfishes. Besides, several limitations of the present study should also be noted. First, some of the observed TRD may reflect isolating mechanisms that have evolved after the completion of reproductive isolation. Second, the slight difference in rearing conditions for the two families could have affected the results. Third, some of the TRD that were unique to larvae of each family might be false positives. This is because markers showing TRD in larvae should also do so in subadults unless the TRD in larvae is counterbalanced by selective mortality at the postnatal stage. Finally, TRD may occur due to mechanisms other than reproductive isolation between parental species, such as inbreeding depression and BDMI involving alleles segregating within species (Corbett-Detig et al., 2013; Plough & Hedgecock, 2011). These issues can be partly addressed by careful control of rearing conditions and appropriate cross designs such as reciprocal F_2_ crosses.

## Acknowledgments

We thank K. Tsuboi (Tokai University) and the staff of Mariculture Fisheries Research Institute, Hokkaido Research Organization for their help in phenotyping and fish husbandry, S. Kondo (Ryukoku University) for help in genotyping. This study was carried out in compliance with the guidelines for the care and use of animals for scientific purposes at Tokai University.

## Author contributions

NM, TK, YoS, and HT conceived the study. TK and YoS performed the crosses and oversaw the fish husbandry. NM, TK, YuS, SI, YoS, and HT performed the phenotyping. NM, RK, and AJN performed the genotyping. NM and RK performed the analyses. NM wrote the manuscript. All authors read and approved the manuscript.

## Data availability

Sequence data have been deposited in DDBJ Sequence Read Archive under accession numbers DRR377585–377864. Genotype and phenotype data and linkage maps are available from Dryad.

