## Supplemental figures for "Genetic architectures of postmating isolation and morphology of two highly diverged rockfishes (genus *Sebastes*)"

**Supplementary figures**

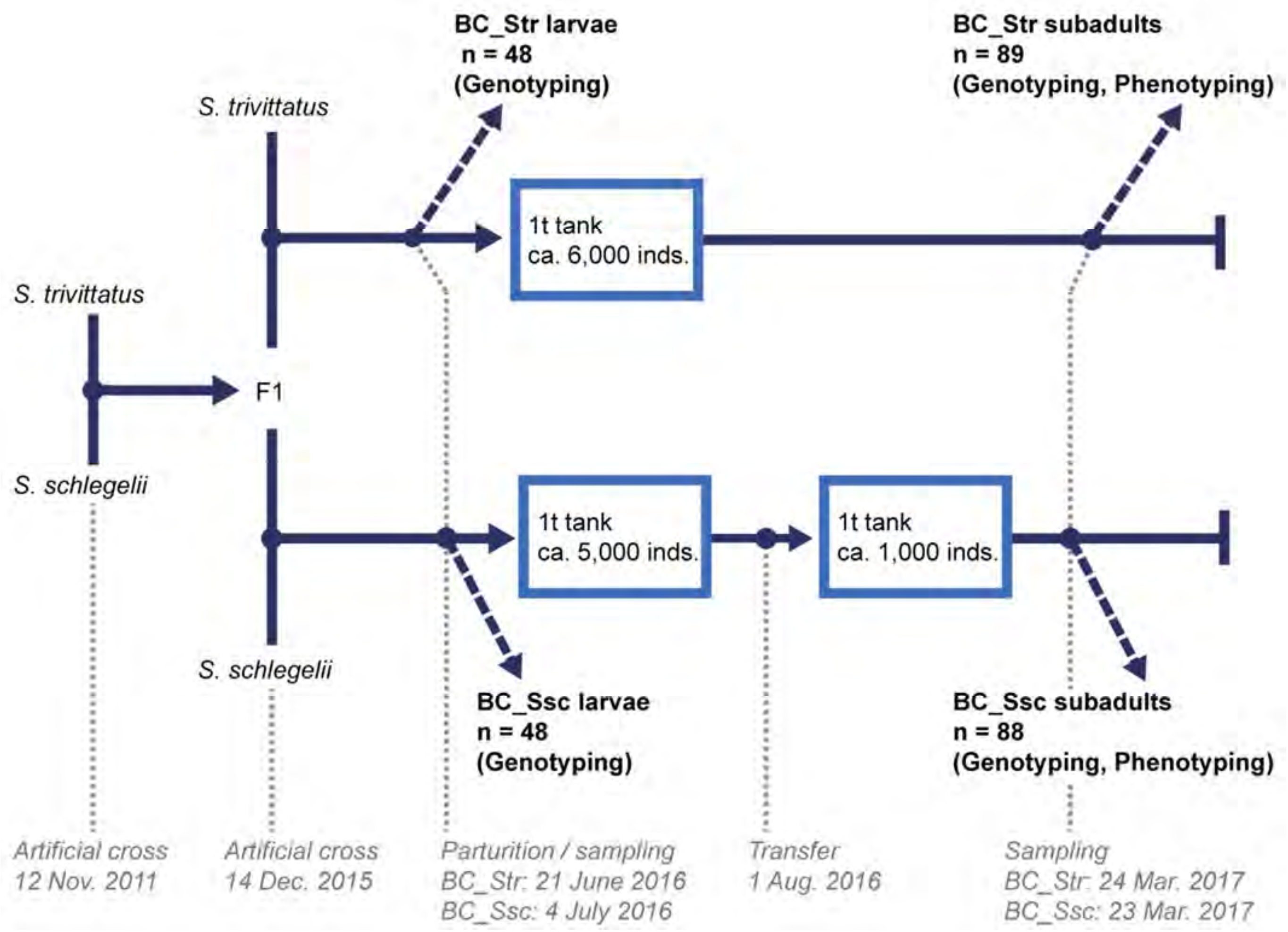


**FIGURE S1** Schematic diagram of the present experimental crosses. The numbers of individuals indicate those used for genotyping and phenotyping.

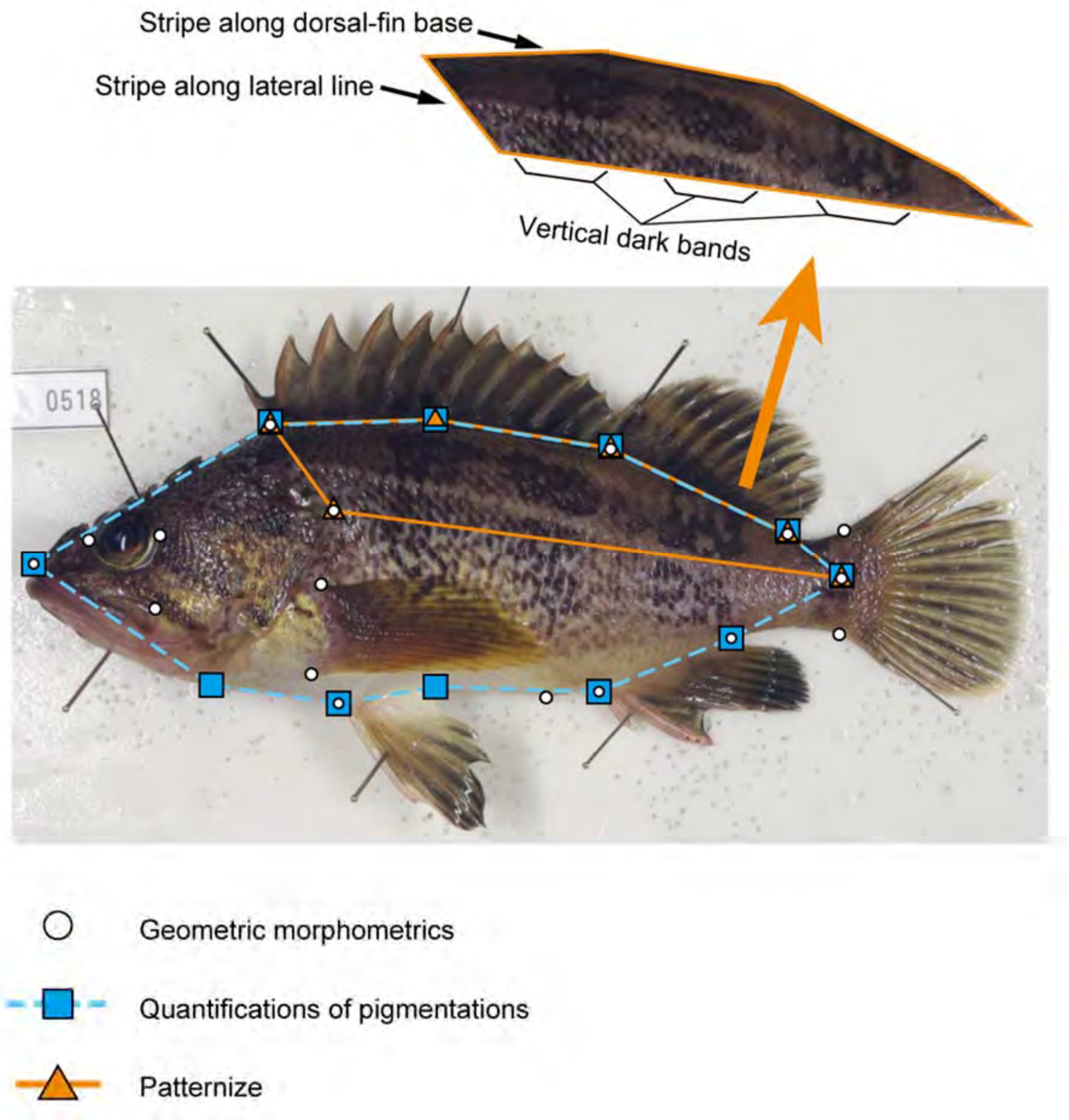


**FIGURE S2** Landmarks defining the polygons for cropping fish images for geometric morphometrics (circles), patternize analysis (triangles), and quantifications of intensities of pigmentations (squares). The color patterns involved in the convergence in rockfishes and phenotyped in the present study, are also shown.

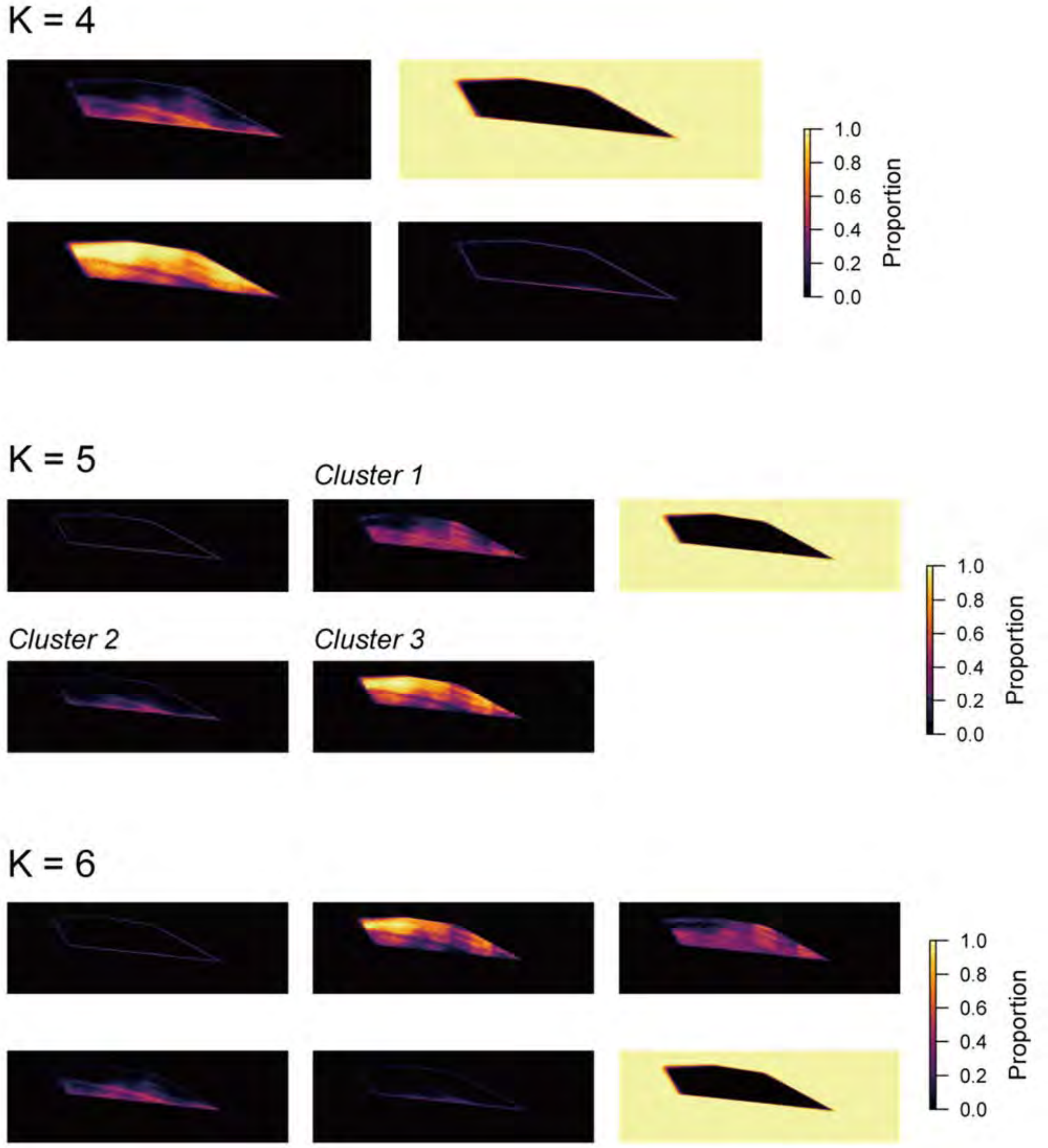


**FIGURE S3** Heatmaps for variation of color patterns in the present mapping families based on the patternize analysis, assuming the number of color clusters (K) = 4–6. The color on the heatmap represents the proportion of individuals having the color cluster at each pixel. Clusters 1–3 in K = 5 were used for QTL mapping.

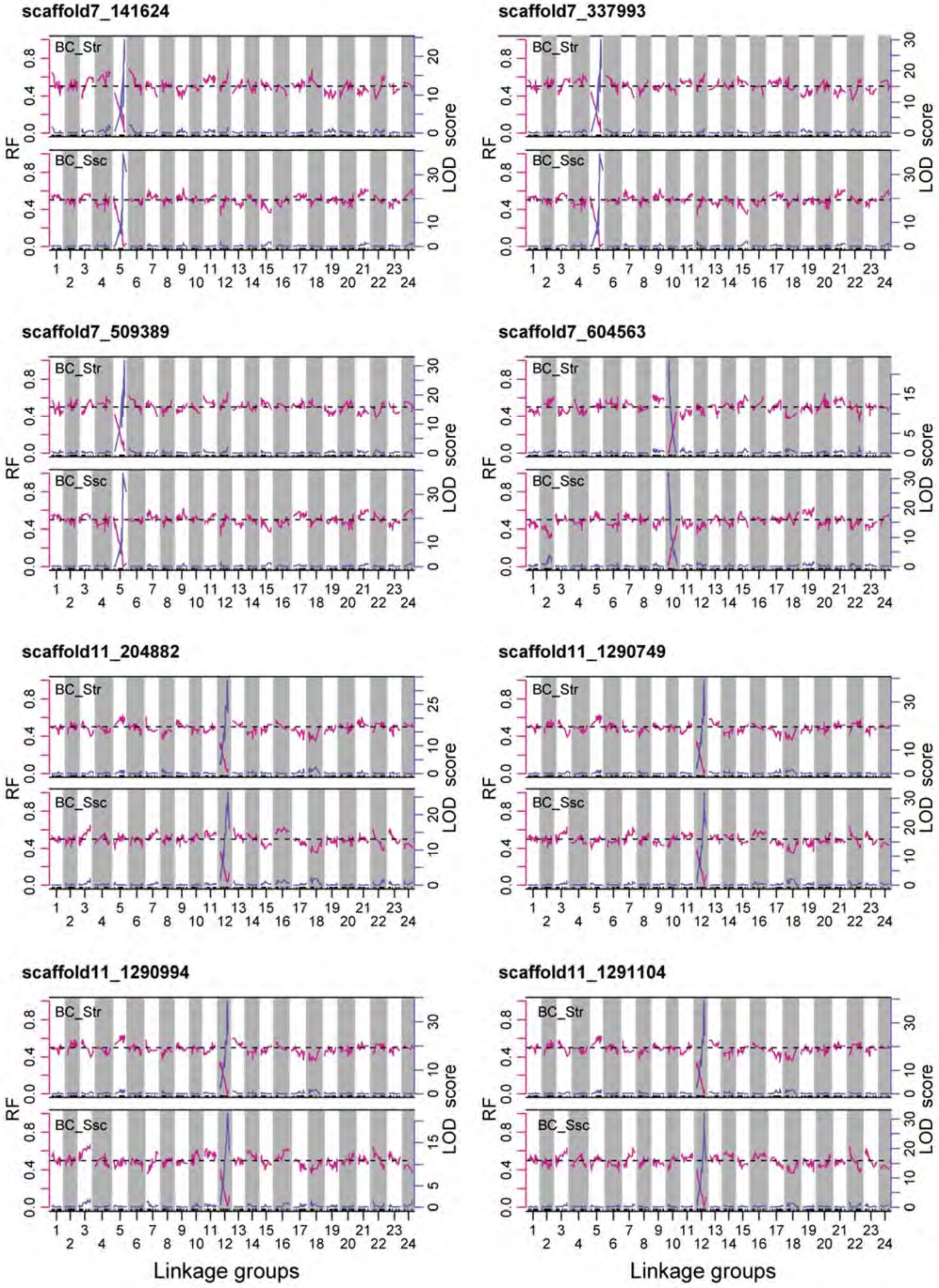
scaffold7_141624
scaffold7_337993
scaffold7_509389
scaffold7_604563
scaffold11_204882
scaffold11_1290749
scaffold11_1290994
scaffold11_1291104
BC_Str
BC_Ssc
RF
LOD score
Linkage groups

Figure S4

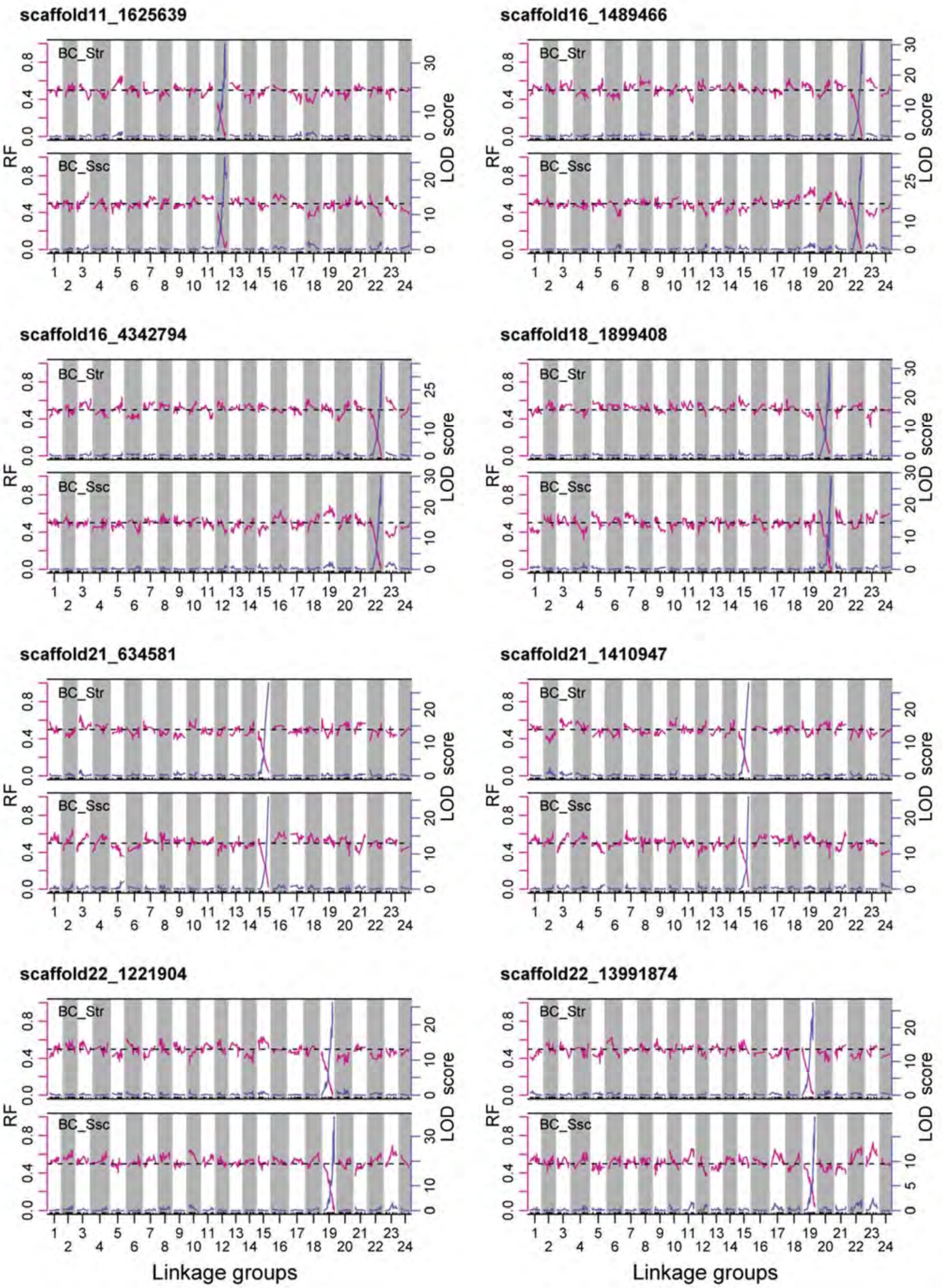


Figure S4 (continued)

**FIGURE S4** Recombination fractions (RF) and LOD scores for the markers eventually mapped to different (non-syntenic) chromosomes/LGs on the consensus map and on the reference genome assembly of *S. schlegelii* after the modifications (see also Table S3 and Appendix S1). Note that these markers did not show either reduced RF or increased LOD score with any LG other than the one to which they were assigned. The marker name above each panel consists of the scaffold number to which the marker is mapped and the physical position (bp) on the scaffold, connected by an underscore.

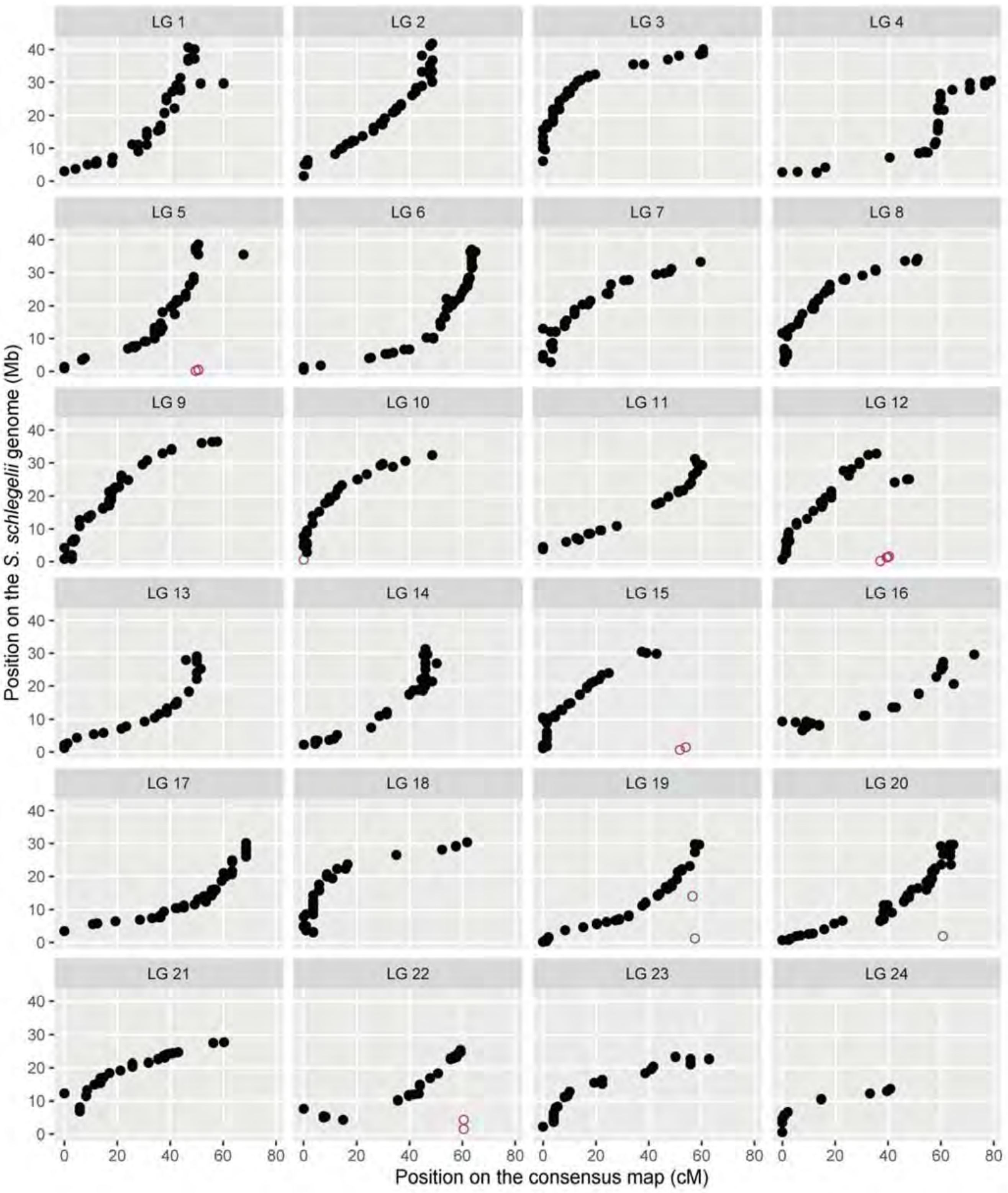


**FIGURE S5** Comparison of marker positions on the consensus map and the reference genome assembly of *S. schlegelii*. Closed and open circles indicate markers mapped to the same (syntenic) and different (non-syntenic) chromosomes/LGs on the consensus map and the reference genome, respectively.

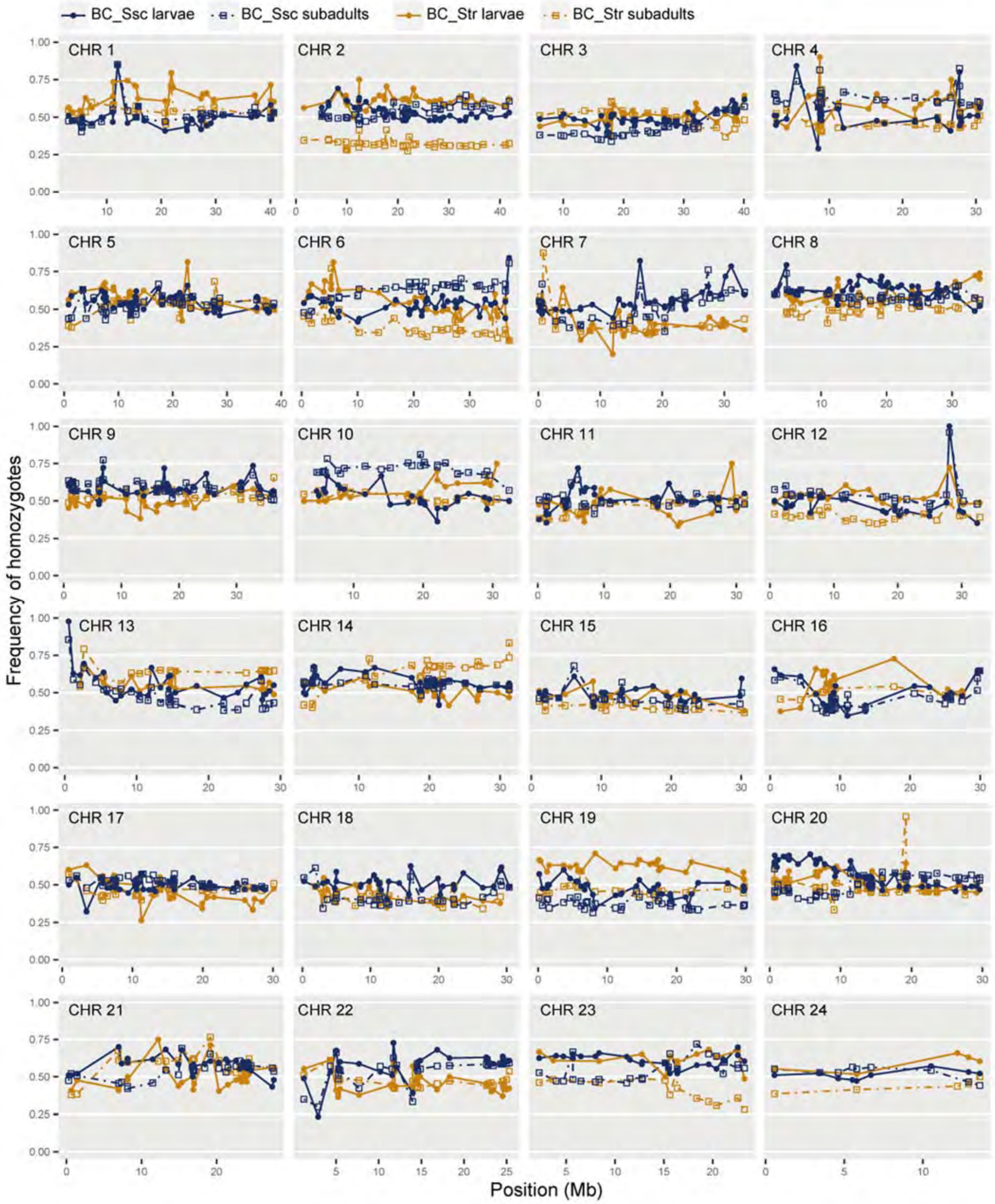


**FIGURE S6** Genotype frequencies of markers shown on the reference genome assembly of *S. schlegelii*. For each marker, frequencies of homozygotes are plotted.

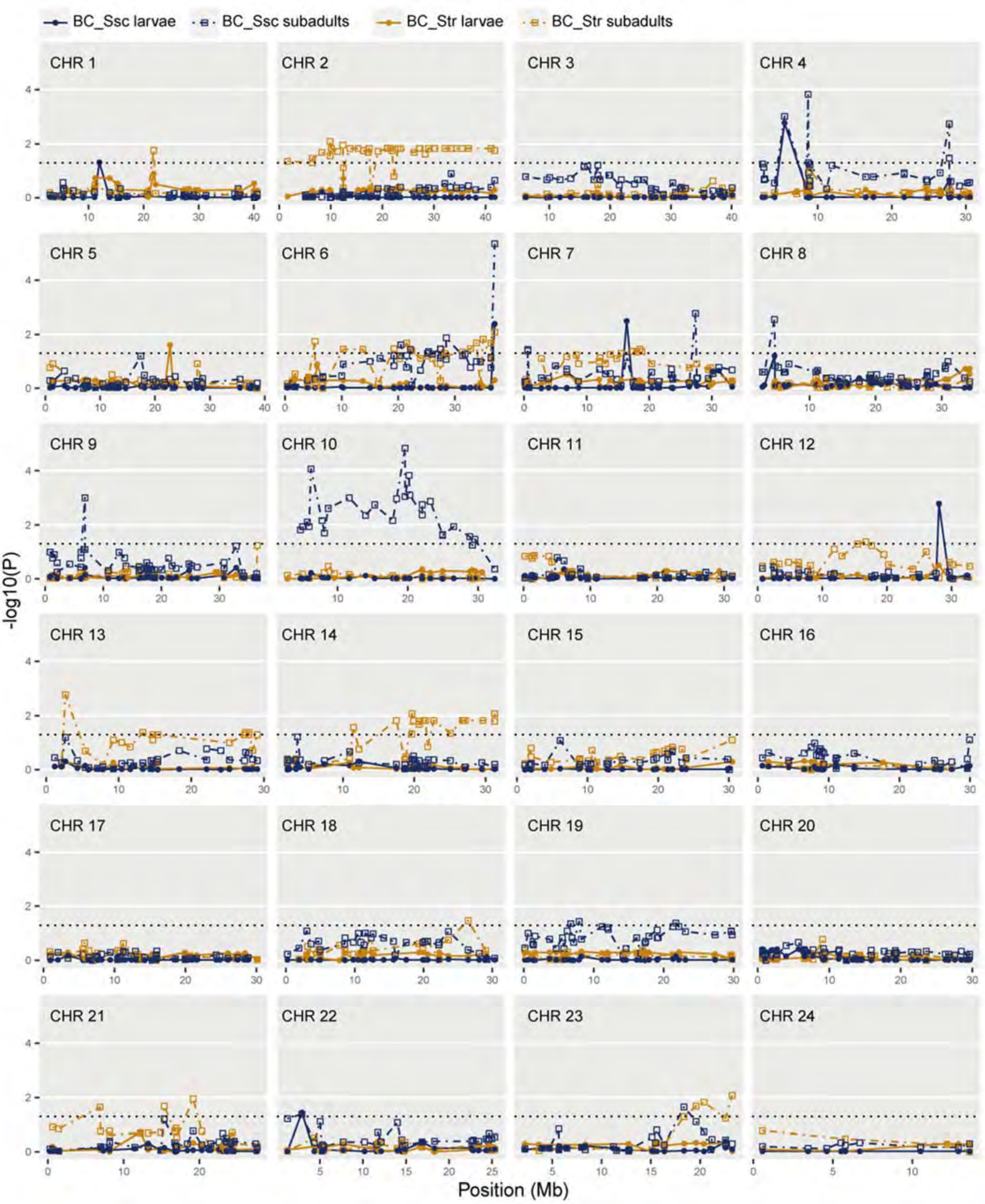


**FIGURE S7** Transmission ratio distortion of markers shown on the reference genome assembly of *S. schlegelii*. For each marker, *p* values from the chi-square tests for the deviations of genotype frequencies from Mendelian expectations corrected for false discovery rates (FDR) are given as $-\log_{10}p$. Dotted line indicates FDR-corrected $p = 0.05$.

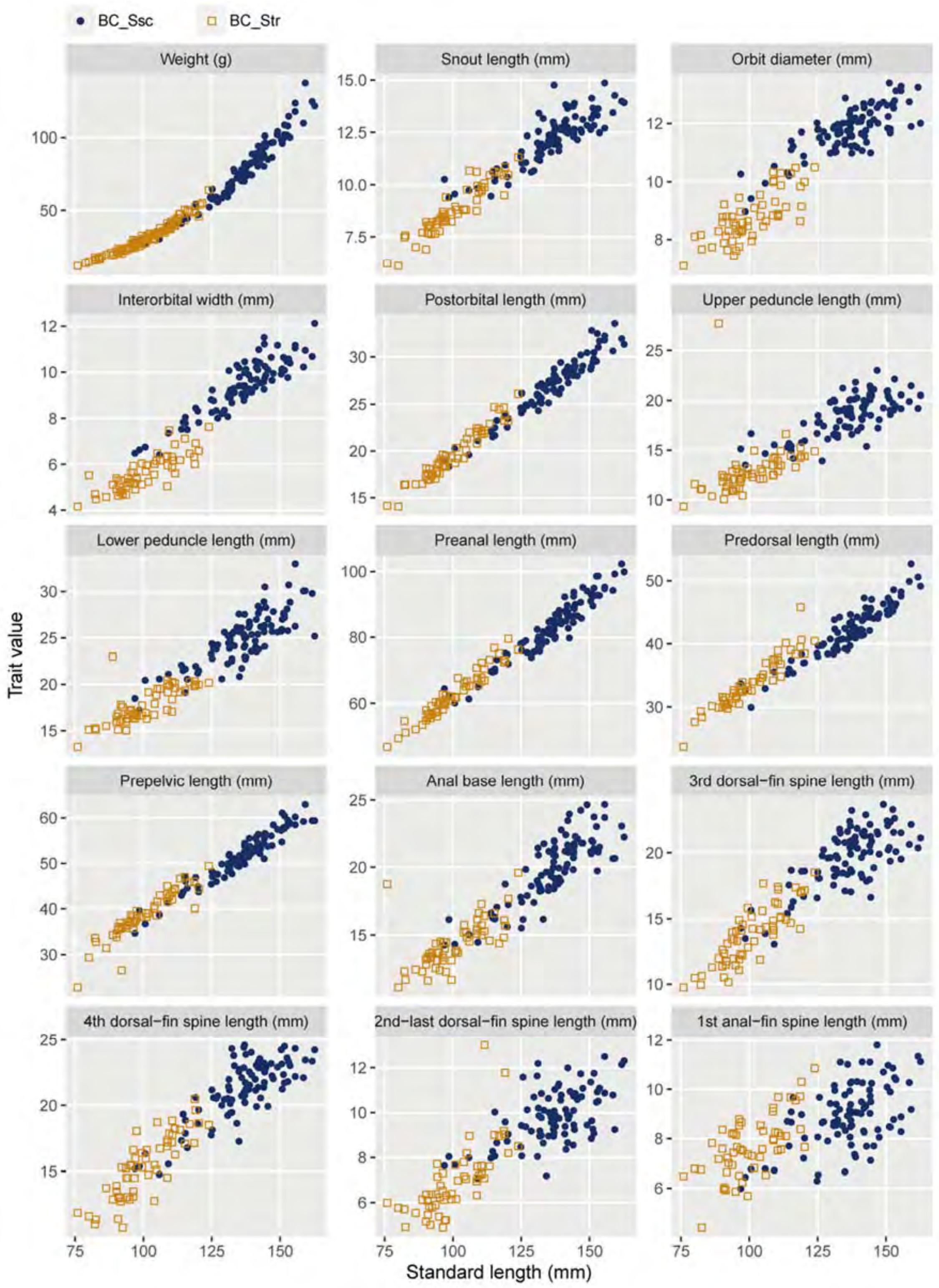
BC_Ssc
BC_Str
Weight (g)
Snout length (mm)
Orbit diameter (mm)
Interorbital width (mm)
Postorbital length (mm)
Upper peduncle length (mm)
Lower peduncle length (mm)
Preanal length (mm)
Predorsal length (mm)
Prepelvic length (mm)
Anal base length (mm)
3rd dorsal−fin spine length (mm)
4th dorsal−fin spine length (mm)
2nd−last dorsal−fin spine length (mm)
1st anal−fin spine length (mm)
Trait value
Standard length (mm)

FIGURE S8

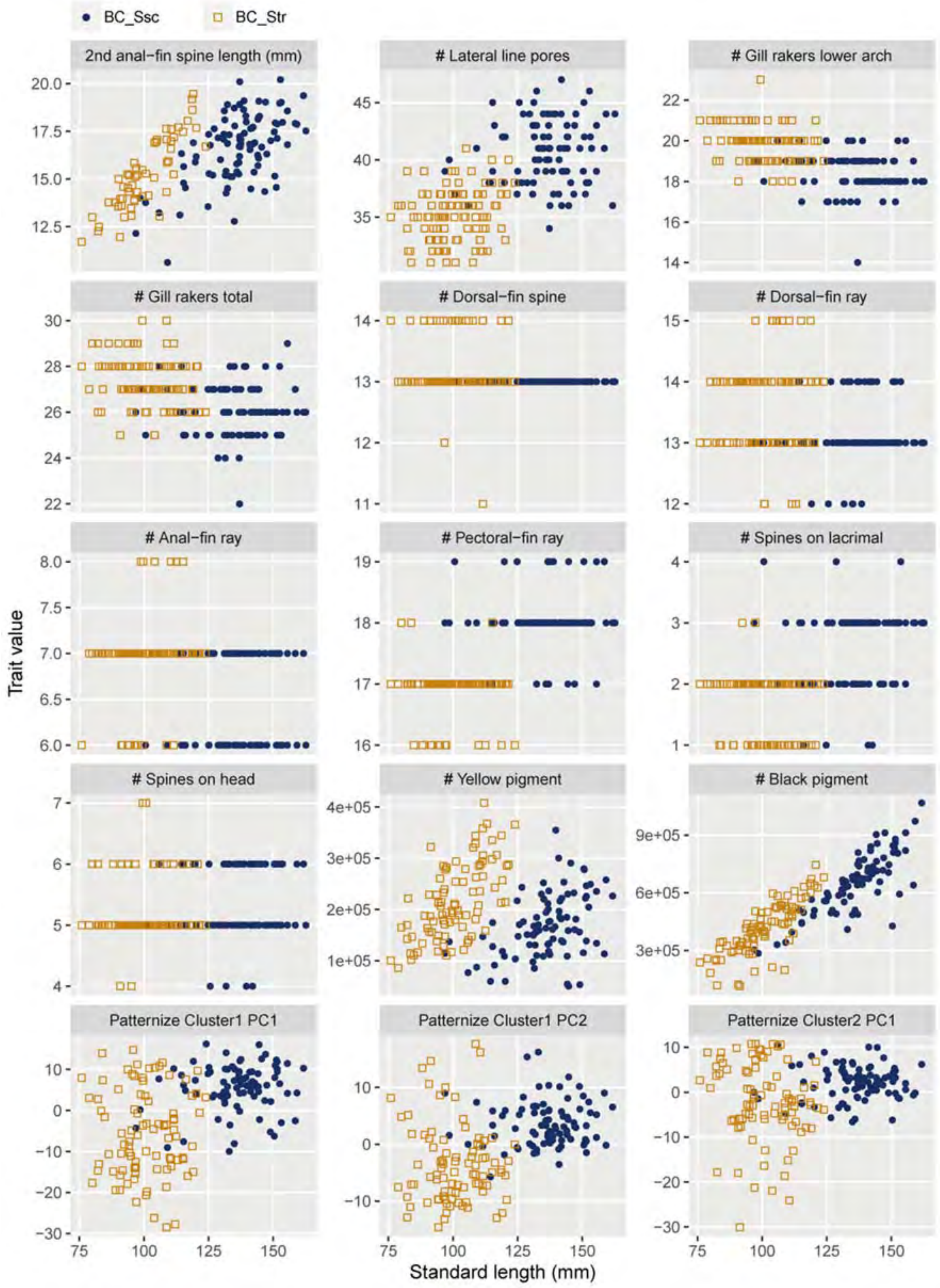


FIGURE S8 (continued)

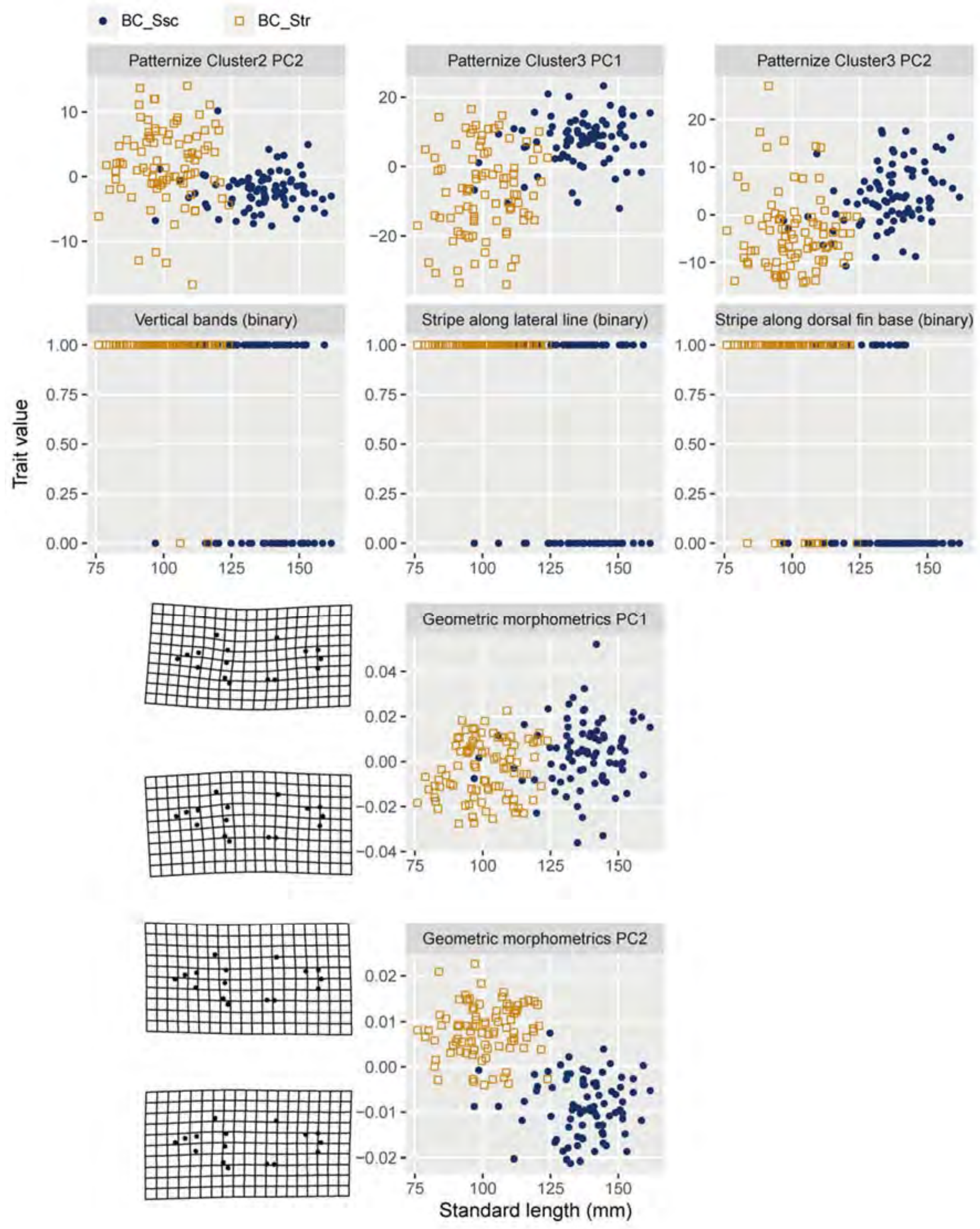


FIGURE S8 (continued)

**FIGURE S8** Morphological traits of the present mapping families. Only the traits used for QTL mapping are shown. Each trait is plotted against size (standard length: SL). For principal component (PC) scores 1 and 2 based on geometric morphometrics, representative shapes (positions of landmarks) for extreme PC scores are also shown.

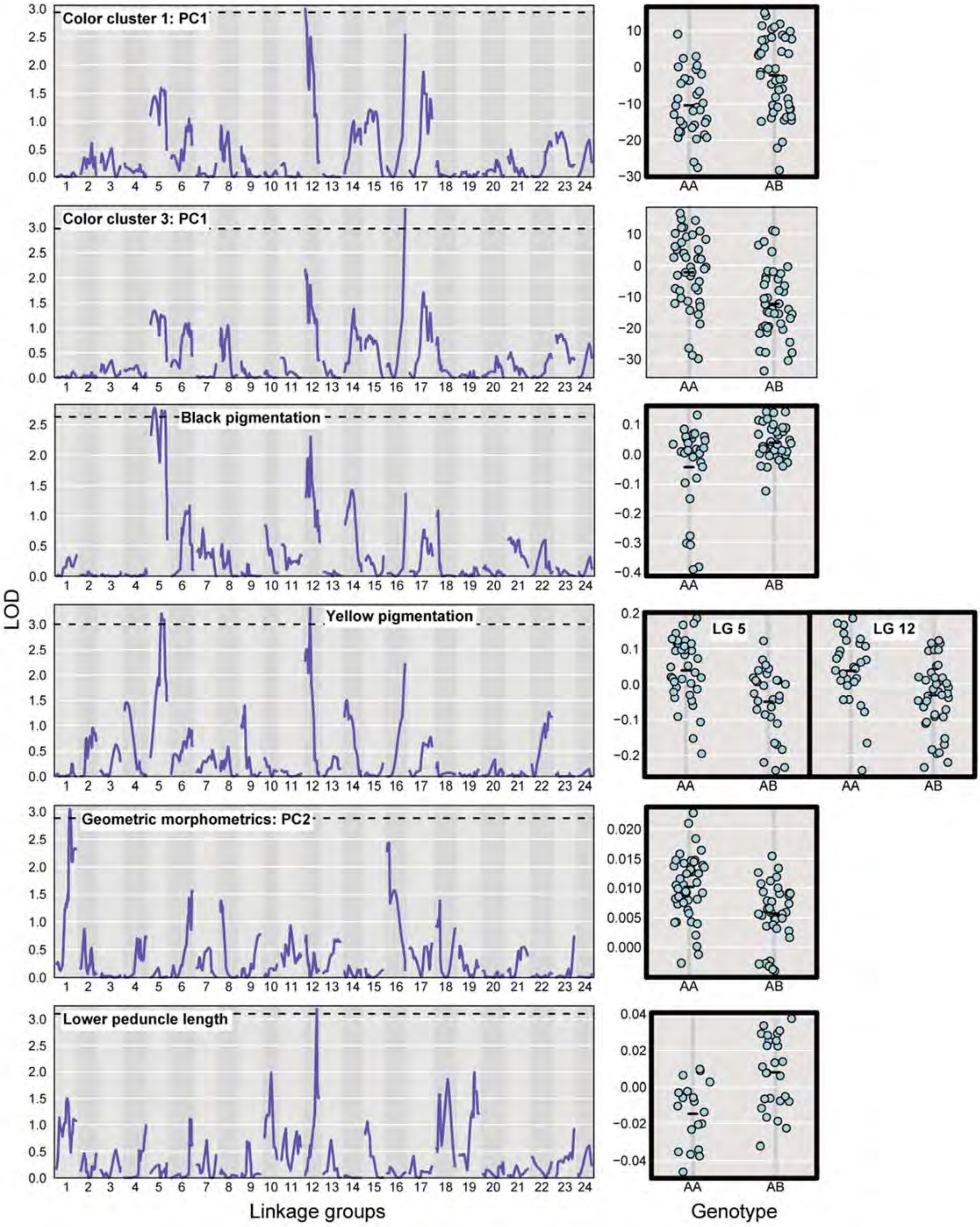
Color cluster 1: PC1
Color cluster 3: PC1
Black pigmentation
Yellow pigmentation
LG 5
LG 12
Geometric morphometrics: PC2
Lower peduncle length
LOD
Linkage groups
Genotype
AA
AB

53 FIGURE S9

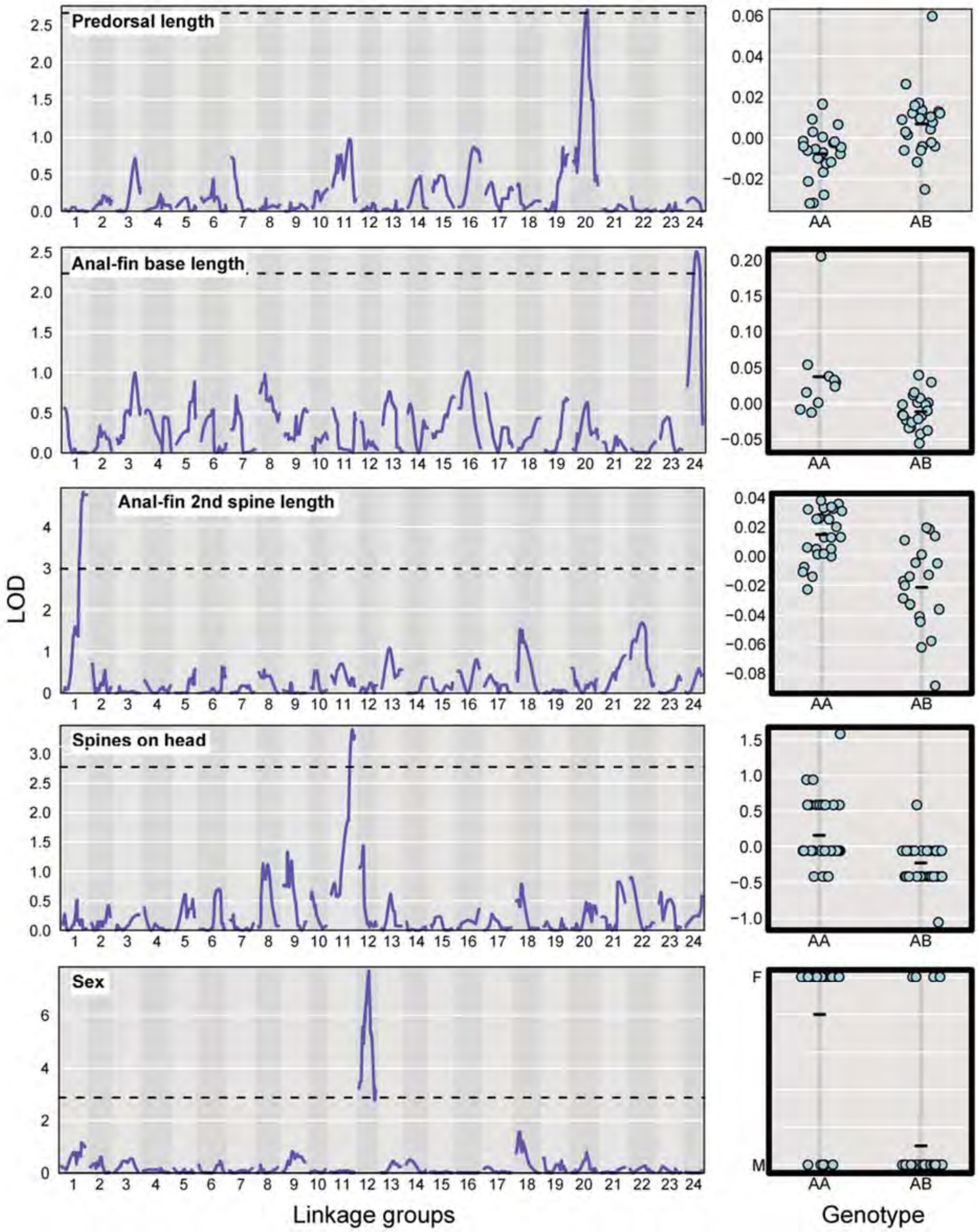


FIGURE S9 (continued)

**FIGURE S9**: LOD scores of QTL mapping across 24 LGs (left panel) and plot of phenotypic values against genotypes at the significant LOD peak (right panel) for each trait of BC_Str. Residuals of regression of each trait on size or sex are plotted if the trait was dependent on these variables, otherwise, raw phenotypic values are plotted. Genome-wide significance thresholds of LOD scores for each trait is indicated by a dotted line. The thick border of the right panel indicates allelic effects in the direction expected from the divergence between the parental species. A and B on the right panel indicate alleles from *S. trivittatus* and *S. schlegelii*, respectively. Only the traits with significant LOD peaks are shown.

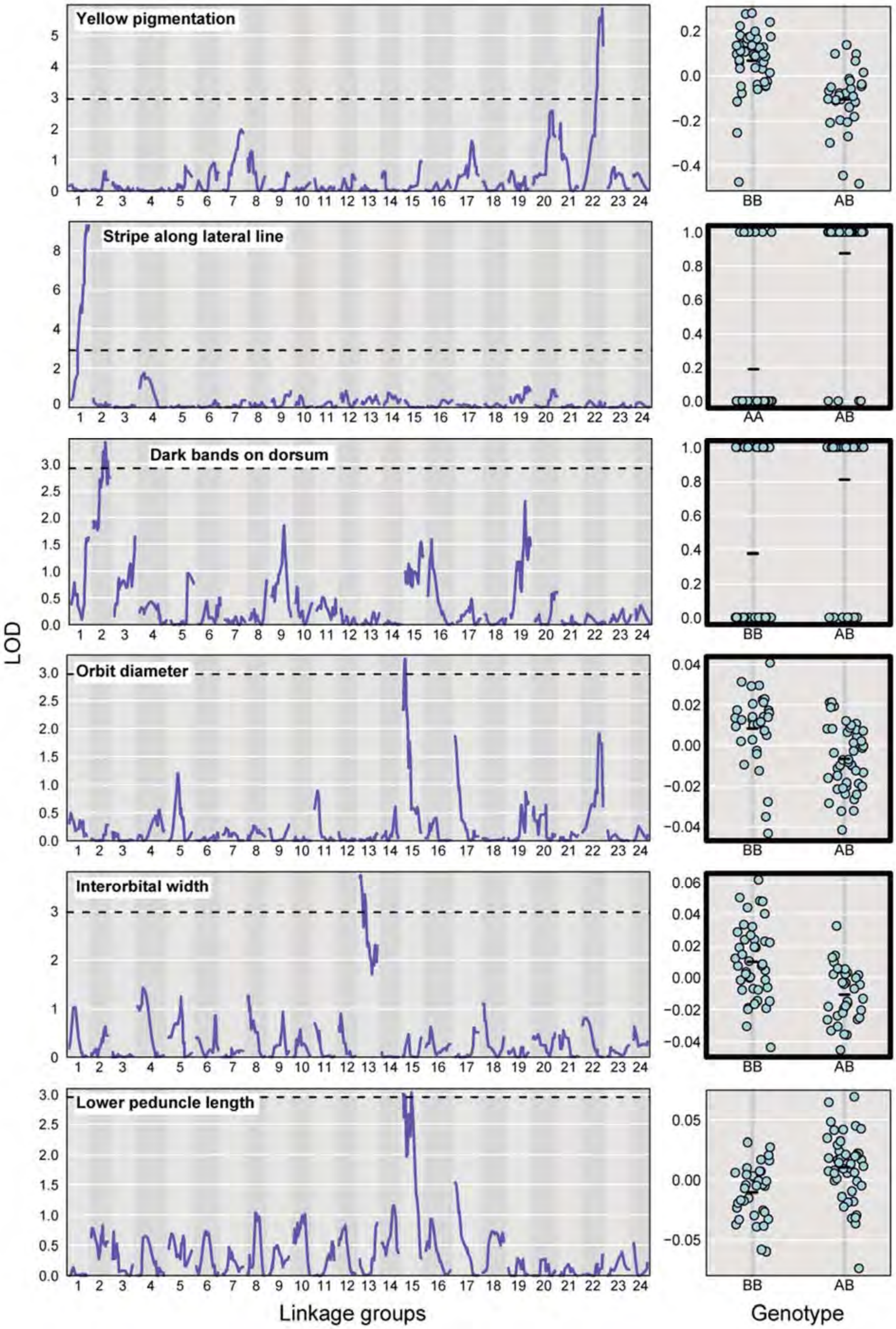


65 FIGURE S10

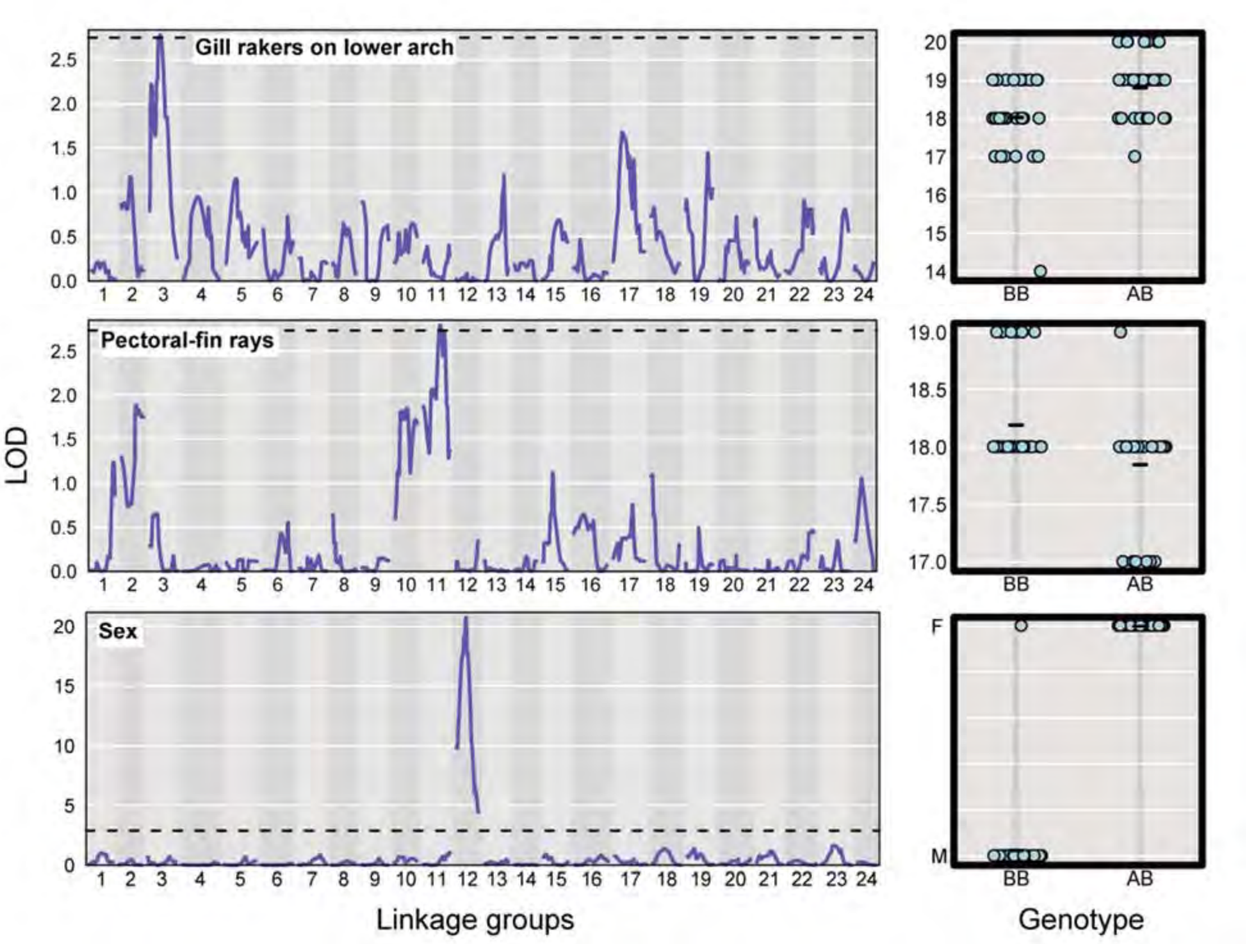


FIGURE S10 (continued)

**FIGURE S10**: LOD scores of QTL mapping across 24 LGs (left panel) and plot of phenotypic values against genotypes at the significant LOD peak (right panel) for each trait of BC_Ssc. Residuals of regression of each trait on size or sex are plotted if the trait was dependent on these variables, otherwise, raw phenotypic values are plotted. Genome-wide significance thresholds of LOD scores for each trait is indicated by a dotted line. The thick border of the right panel indicates allelic effects in the direction expected from the divergence between the parental species. A and B on the right panel indicate alleles from *S. trivittatus* and *S. schlegelii*, respectively. Only the traits with significant LOD peaks are shown.
