## Supplemental tables for "Genetic architectures of postmating isolation and morphology of two highly diverged rockfishes (genus *Sebastes*)"

### Supplementary tables

**TABLE S1** Differences in continuous morphological traits between families and sexes. Effects of size (standard length), family, sex, and their interactions, expressed as *p* values for the corresponding terms of linear regressions are presented. For size, family, and sex terms, *p* values based on the regression including an interaction term are presented if the interaction term was significant: otherwise, *p* values based on the regression excluding the interaction term are presented. If neither size term nor interaction term was significant, *p* values based on ANOVA (comparing between family or sex) are presented. *P* values < 0.01 are in bold type.

| | Both families | | BC_Str | | | BC_Ssc | | |
|---|---|---|---|---|---|---|---|---|
| | Family | Interaction: size (SL) x family | Size (SL) | Sex | Interaction: size (SL) x sex | Size (SL) | Sex | Interaction: size (SL) x sex |
| Standard length (SL) | **<2E-16**[b] | NA | NA | **0.775**[b] | NA | NA | **0.848**[b] | NA |
| Coloration | | | | | | | | |
| Color cluster 1: PC1 | **1.2E-05** | 0.36 | 0.73 | **0.061**[b] | 0.023 | 0.10 | **0.94**[b] | 0.73 |
| Color cluster 1: PC2 | **5.3E-07** | 0.35 | 0.44 | **0.84**[b] | 0.81 | 0.75 | **0.51**[b] | 0.44 |
| Color cluster 2: PC1 | **6.8E-04** | 0.15 | 0.38 | **8.1E-03**[b] | 0.95 | 0.39 | **0.53**[b] | 0.34 |
| Color cluster 2: PC2 | **4.2E-04** | 0.84 | 0.90 | **0.82**[b] | 0.42 | 0.72 | **0.50**[b] | 0.36 |
| Color cluster 3: PC1 | **1.1E-04** | 0.87 | 0.16 | **0.087**[b] | 0.10 | 0.029 | **0.81**[b] | 0.39 |
| Color cluster 3: PC2 | **5.1E-03** | 0.034 | 0.67 | **0.37**[b] | 0.50 | **4.1E-03** | 0.082 | 0.67 |
| Yellow pigmentation | **2.0E-16** | 0.27 | **1.2E-07** | **8.3E-03** | 0.31 | **1.5E-03** | **2.4E-03** | 0.89 |
| Black pigmentation | **6.3E-05** | 0.21 | **4.0E-13** | 0.030 | 0.89 | **2.0E-16** | 0.72 | 0.88 |
| Shape | | | | | | | | |
| Weight | **<2E-16**[b] | 0.30 | **<2e-16** | **0.797**[b] | 0.69 | **<2E-16** | **0.805**[b] | 0.43 |
| Head length | 0.11 | 0.049 | **<2e-16** | 0.34 | 0.96 | **<2E-16** | 0.59 | 0.06 |

| | | | | | | | | |
|---|---|---|---|---|---|---|---|---|
| Snout length | **1.6E-04**[a] | **2.2E-04** | **<2e-16** | 0.50 | 0.10 | **<2E-16** | 0.02 | 0.24 |
| Orbit diameter | **3.3E-13** | 0.23 | **2.9E-07** | 1.00 | 0.51 | **<2E-16** | 0.87 | 0.82 |
| Interorbital width | **<2E-16** | 0.35 | **1.4E-09** | 0.09 | 0.28 | **<2E-16** | 0.03 | 0.65 |
| Postorbital length | **5.4E-04** | 0.03 | **<2e-16** | 0.69 | 0.29 | **<2E-16** | 0.11 | 0.05 |
| Upper-jaw length | 0.70 | 0.39 | **<2e-16** | 0.28 | 0.63 | **<2E-16** | 0.06 | 0.74 |
| Body depth 1 | 0.45 | 0.84 | **<2e-16** | 0.12 | 0.73 | **<2E-16** | 0.17 | 0.91 |
| Body depth 2 | 0.16 | 0.42 | **<2e-16** | 0.63 | 0.33 | **<2E-16** | 0.79 | 0.65 |
| Body width | 0.13 | 0.41 | **8.5E-16** | 0.04 | 0.96 | **<2E-16** | 0.05 | 0.20 |
| Caudal peduncle depth | 0.31 | 0.60 | **<2e-16** | 0.12 | 0.37 | **<2E-16** | 0.59 | 0.86 |
| Upper peduncle length | **1.2E-05** | 0.53 | **1.5E-02** | 0.46 | 0.20 | **8.6E-16** | 0.27 | 0.21 |
| Lower peduncle length | **4.3E-05** | 0.15 | **4.6E-08** | 0.58 | 0.43 | **<2E-16** | 0.07 | 0.75 |
| Pectoral-fin length | 0.31 | 0.05 | **2.5E-14** | 0.96 | 0.58 | **<2E-16** | 0.37 | 0.49 |
| Pelvic fin length | 0.16 | 0.32 | **<2e-16** | 0.26 | 0.56 | **<2E-16** | 0.15 | 0.03 |
| Dorsal-fin base length | 0.73 | 0.48 | **<2e-16** | 0.60 | 0.76 | **<2E-16** | 0.61 | 0.34 |
| Spinous portion of dorsa-fin base | 0.37 | 0.79 | **<2e-16** | 0.30 | 0.37 | **<2E-16** | 0.20 | 0.06 |
| Soft-rayed portion of dorsal-fin base | 0.86 | 0.28 | **1.7E-13** | 0.55 | 0.11 | **<2E-16** | 0.91 | 0.49 |
| Preanal length | **2.2E-03** | 0.58 | **<2e-16** | 0.84 | 0.92 | **<2E-16** | 0.23 | 0.79 |
| Predorsal length | **9.2E-10** | 0.088 | **<2e-16** | 0.64 | 0.45 | **<2E-16** | 0.14 | 0.39 |
| Prepelvic length | **8.1E-03** | 0.086 | **4.5E-14** | 0.10 | 0.66 | **<2E-16** | 0.17 | 0.50 |
| Anal-fin base length | 0.013[a] | **9.9E-03** | **6.0E-13** | 0.94 | 0.21 | **<2E-16** | 0.40 | 0.29 |
| Pelvic-to-anal-fin length | 0.19 | 0.018 | **5.7E-09** | 0.78 | 0.59 | **<2E-16** | 0.20 | 0.70 |
| Dorsal-fin 1st spine length | 0.040 | 0.27 | **5.6E-09** | 0.98 | 0.14 | **<2E-16** | 0.11 | 0.96 |
| Dorsal-fin 2nd spine length | 0.31 | 0.18 | **1.1E-08** | 0.32 | 0.14 | **<2E-16** | 0.35 | 0.39 |

| | | | | | | | | |
|---|---|---|---|---|---|---|---|---|
| Dorsal-fin 3rd spine length | **3.7E-03**[a] | **4.7E-03** | **8.8E-13** | 0.53 | 0.92 | **3.7E-16** | 0.31 | 0.28 |
| Dorsal-fin 4th spine length | **4.6E-03**[a] | **4.9E-03** | **3.6E-13** | 0.38 | 0.86 | **<2E-16** | 0.39 | 0.80 |
| Dorsal-fin 5th spine length | 0.72 | 0.015 | **8.4E-14** | 0.82 | 0.95 | **<2E-16** | 0.80 | 0.59 |
| Dorsal-fin 3rd last spine length | 0.80 | 0.089 | **3.3E-07** | 0.97 | 0.62 | **5.4E-14** | 0.11 | 0.68 |
| Dorsal-fin 2nd last spine length | **4.3E-04**[a] | **5.3E-04** | **3.5E-09** | 0.81 | 0.66 | **5.7E-11** | **1.1E-03** | 0.31 |
| Dorsal-fin last spine length | 0.58 | 0.052 | **2.6E-11** | 0.97 | 0.90 | **9.5E-12** | 0.81 | 0.66 |
| Anal-fin 1st spine length | **2.9E-03** | 0.46 | **4.4E-07** | 0.49 | 0.02 | **0.13a** | **7.6E-04**[a] | **6.6E-04** |
| Anal-fin 2nd spine length | **7.3E-09** | 0.036 | **1.3E-12** | 0.63 | 0.75 | **9.1E-10** | 0.36 | 0.89 |
| Anal-fin 3rd spine length | 0.80 | 0.62 | **1.7E-11** | 0.34 | 0.94 | **<2E-16** | 0.07 | 0.62 |
| Pelvic-fin spine length | 0.015 | 0.44 | **<2e-16** | 0.03 | 0.44 | **1.5E-11** | 0.88 | 0.89 |
| Length of gill raker at angle | 0.96 | 0.94 | **1.5E-06** | 0.23 | 1.00 | **9.3E-08** | 0.09 | 0.69 |
| Geometric morphometrics: PC1 | **1.1E-3**[b] | 0.71 | 0.39 | **0.22**[b] | 0.23 | 0.44 | **0.11**[b] | 0.61 |
| Geometric morphometrics: PC2 | **<2E-16**[b] | 0.48 | 0.25 | **0.38**[b] | 0.36 | 0.50 | **0.078**[b] | 0.62 |

[a]Based on linear regressions including an interaction term (size by family or size by sex).

[b]Based on ANOVA.

**TABLE S2** Differences in discrete morphological traits between families and sexes. Countable and binary traits were tested using Wilcoxon's rank test and Fisher's exact test, respectively. *P* values < 0.01 are in bold type.

| | | Between sexes | |
|---|---|---|---|
| | Between families | BC_Str | BC_Ssc |
| Coloration | | | |
| Dark bands on dorsum | **1.1E-10** | 0.51 | 1.00 |
| Stripe along lateral line | **2.6E-13** | NA[a] | 0.50 |
| Stripe along dorsal-fin base | **2.2E-16** | 0.037 | 0.24 |
| Counts | | | |
| Lateral line pores | **<2E-16** | 0.88 | 0.36 |
| Gill rakers on upper arch | 0.014 | 0.94 | 0.95 |
| Gill rakers on lower arch | **<2E-16** | 0.67 | 0.59 |
| Gill rakers total | **6.2E-12** | 0.85 | 0.63 |
| Dorsal-fin spines | **2.4E-04** | 0.64 | NA[a] |
| Dorsa-fin soft-rays | **7.6E-08** | 0.83 | 0.019 |
| Anal-fin spines | 0.31 | NA[a] | 0.36 |
| Anal-fin soft-rays | **3.6E-07** | 0.59 | 0.95 |
| Pectoral-fin rays | **<2E-16** | 0.97 | 0.68 |
| Pelvic-fin spines | NA[a] | NA[a] | NA[a] |
| Pelvic-fin soft-rays | 0.015 | 0.54 | 0.81 |
| Spines on lacrimal | **<2E-16** | 0.47 | 0.060 |
| Spines on head | **6.5E-03** | **4.8E-04** | 0.56 |

[a]No variation observed.

**TABLE S3** Markers with inconsistent assignment patterns between the initial formation of linkage groups in the present study and the mapping to the reference genome assembly of *S. schlegelii*, and modifications made for such markers based on examinations of the individual LOD scores and recombination fractions. See Appendix S1 for details of the modifications. Results of the chi-square tests for transmission ratio distortion (*p* values after false discovery rate correction) are also shown. The marker name consists of a combination of the scaffold number to which the marker is mapped and the physical position (bp) on the scaffold, connected by an underscore.

| | | Initial map LG[b] | | Modified map LG[c] | | | TRD (FDR-corrected *p* value) | |
|---|---|---|---|---|---|---|---|---|
| Marker | CHR[a] | BC_Str | BC_Ssc | BC_Str | BC_Ssc | Modification | BC_Str | BC_Ssc |
| scaffold16_17681489 | 16 | **24** | NA | 16 | NA | Integrated to LG16 in BC_Str | 0.51 | NA |
| scaffold16_24969576 | 16 | **24** | 16 | 16 | 16 | Integrated to LG16 in BC_Str | 0.79 | 0.42 |
| scaffold16_25385024 | 16 | **24** | 16 | 16 | 16 | Integrated to LG16 in BC_Str | 0.89 | 0.75 |
| scaffold16_25805992 | 16 | **24** | 16 | 16 | 16 | Integrated to LG16 in BC_Str | 0.79 | 0.51 |
| scaffold16_25806172 | 16 | **24** | 16 | 16 | 16 | Integrated to LG16 in BC_Str | 0.89 | 0.83 |
| scaffold16_27402848 | 16 | **24** | 16 | 16 | 16 | Integrated to LG16 in BC_Str | 0.92 | 0.51 |
| scaffold24_565278 | 24 | **26** | 24 | 24 | 24 | Integrated with LG25 in BC_Str | 0.50 | 0.69 |
| scaffold24_5786991 | 24 | **26** | 24 | 24 | 24 | Integrated with LG25 in BC_Str | 0.48 | 0.74 |
| scaffold24_12198673 | 24 | **25** | NA | 24 | NA | Integrated with LG26 in BC_Str | 0.89 | NA |
| scaffold24_12907216 | 24 | **25** | 24 | 24 | 24 | Integrated with LG26 in BC_Str | 0.87 | 0.86 |
| scaffold24_13617396 | 24 | **25** | 24 | 24 | 24 | Integrated with LG26 in BC_Str | 0.60 | 0.69 |
| scaffold4_2539830 | 4 | NA | **25** | NA | 4 | Integrated to LG4 in BC_Ssc | NA | 0.11 |
| scaffold4_2732737 | 4 | NA | **25** | NA | 4 | Integrated to LG4 in BC_Ssc | NA | 0.19 |
| scaffold4_2765933 | 4 | 4 | **25** | 4 | 4 | Integrated to LG4 in BC_Ssc | 0.92 | 0.28 |
| scaffold4_2849964 | 4 | NA | **25** | NA | 4 | Integrated to LG4 in BC_Ssc | NA | 0.37 |
| scaffold4_4189686 | 4 | 4 | **25** | 4 | 4 | Integrated to LG4 in BC_Ssc | 0.92 | 0.40 |
| scaffold7_141624 | 7 | 5 | 5 | 5 | 5 | Left unmodified | 0.70 | 0.69 |

| | | | | | | | | |
|---|---|---|---|---|---|---|---|---|
| scaffold7_337993 | 7 | 5 | 5 | 5 | 5 | Left unmodified | 0.41 | 0.60 |
| scaffold7_509389 | 7 | 5 | 5 | 5 | 5 | Left unmodified | 0.50 | 0.60 |
| scaffold7_604563 | 7 | 10 | 10 | 10 | 10 | Left unmodified | 0.83 | 0.15 |
| scaffold11_204882 | 11 | 12 | 12 | 12 | 12 | Left unmodified | 0.26 | 0.60 |
| scaffold11_1290749 | 11 | 12 | 12 | 12 | 12 | Left unmodified | 0.30 | 0.72 |
| scaffold11_1290994 | 11 | 12 | 12 | 12 | 12 | Left unmodified | 0.30 | 0.27 |
| scaffold11_1291104 | 11 | 12 | 12 | 12 | 12 | Left unmodified | 0.21 | 0.83 |
| scaffold11_1625639 | 11 | 12 | 12 | 12 | 12 | Left unmodified | 0.33 | 0.79 |
| scaffold16_1489466 | 16 | 22 | 22 | 22 | 22 | Left unmodified | 0.39 | 0.11 |
| scaffold16_4342794 | 16 | 22 | 22 | 22 | 22 | Left unmodified | 0.41 | 0.22 |
| scaffold18_1899408 | 18 | 20 | 20 | 20 | 20 | Left unmodified | 0.89 | 0.52 |
| scaffold21_634581 | 21 | 15 | 15 | 15 | 15 | Left unmodified | 0.12 | 0.83 |
| scaffold21_1410947 | 21 | 15 | 15 | 15 | 15 | Left unmodified | 0.25 | 0.86 |
| scaffold22_1221904 | 22 | 19 | 19 | 19 | 19 | Left unmodified | 0.74 | 0.12 |
| scaffold22_13991874 | 22 | 19 | 19 | 19 | 19 | Left unmodified | 0.92 | 0.07 |
| scaffold5_5483780 | 5 | NA | 12 | NA | NA | Discarded | NA | 0.33 |
| scaffold7_604409 | 7 | NA | 10 | NA | NA | Discarded | NA | 0.07 |
| scaffold7_817258 | 7 | NA* | 5 | NA | NA | Discarded | 3.1.E-06 | 1.00 |
| scaffold11_5039720 | 11 | NA | 2 | NA | NA | Discarded | NA | 0.09 |
| scaffold16_627436 | 16 | NA | 22 | NA | NA | Discarded | NA | 0.13 |
| scaffold16_29600040 | 16 | NA | 22 | NA | NA | Discarded | NA | 0.45 |
| scaffold16_29938855 | 16 | NA | 14 | NA | NA | Discarded | NA | 0.03 |
| scaffold17_912416 | 17 | 21 | NA | NA | NA | Discarded | 0.29 | NA |
| scaffold17_960284 | 17 | NA | 21 | NA | NA | Discarded | NA | 0.80 |
| scaffold17_2041277 | 17 | NA | 21 | NA | NA | Discarded | NA | 0.43 |

| | | | | | | | | |
|---|---|---|---|---|---|---|---|---|
| scaffold18_213337 | 18 | NA | 20 | NA | NA | Discarded | NA | 0.60 |
| scaffold21_277480 | 21 | NA | 15 | NA | NA | Discarded | NA | 0.86 |
| scaffold22_13991674 | 22 | 19 | NA | NA | NA | Discarded | 0.70 | NA |

*Removed from the BC_Str dataset before forming linkage groups because of the severe transmission ratio distortion ($p$ < 0.001 after FDR correction)

[a]Position on the reference genome assembly of *S. schlegelii*.

[b]Position on the initial linkage map.

[c]Position on the modified (final) consensus linkage map.

Abbreviation: CHR, chromosome; FDR, false discovery rate; LG, linkage group; TRD, transmission ratio distortion.

**TABLE S4** Summary linkage map statistics for the consensus map. For each linkage group, the size (mega base) of the corresponding chromosome of the reference genome of *S. schlegelii* is also shown.

| | # markers | | | Size | | |
|---|---|---|---|---|---|---|
| LG | BC_Str | BC_Ssc | Shared | cM[a] | cM[b] | Mb |
| 1 | 27 | 33 | 21 | 60.2 | 60.2 | 37.6 |
| 2 | 38 | 44 | 35 | 48.7 | 48.7 | 40.2 |
| 3 | 32 | 39 | 31 | 60.5 | 60.5 | 34.0 |
| 4 | 29 | 30 | 22 | 78.9 | 78.9 | 28.0 |
| 5 | 39 | 48 | 35 | 67.7 | 67.7 | 37.8 |
| 6 | 32 | 37 | 23 | 65.0 | 65.0 | 36.6 |
| 7 | 22 | 30 | 17 | 59.6 | 59.6 | 30.5 |
| 8 | 35 | 45 | 29 | 51.1 | 51.1 | 31.4 |
| 9 | 34 | 42 | 27 | 57.8 | 57.8 | 35.7 |
| 10 | 21 | 26 | 16 | 48.6 | 48.6 | 29.5 |
| 11 | 22 | 27 | 21 | 60.2 | 60.2 | 27.4 |
| 12 | 28 | 29 | 20 | 47.9 | 47.9 | 32.1 |
| 13 | 18 | 29 | 17 | 51.6 | 51.6 | 27.9 |
| 14 | 26 | 33 | 22 | 50.2 | 50.3 | 29.1 |
| 15 | 25 | 27 | 19 | 54.0 | 42.9 | 29.3 |
| 16 | 16 | 24 | 13 | 72.6 | 72.6 | 23.1 |
| 17 | 31 | 40 | 26 | 68.7 | 68.7 | 26.7 |
| 18 | 23 | 30 | 18 | 61.8 | 61.8 | 27.3 |
| 19 | 28 | 38 | 26 | 59.1 | 59.1 | 29.6 |

| | | | | | | |
|---|---|---|---|---|---|---|
| 20 | 41 | 55 | 38 | 64.7 | 64.7 | 29.0 |
| 21 | 24 | 29 | 21 | 60.4 | 60.4 | 20.7 |
| 22 | 27 | 30 | 24 | 60.5 | 59.3 | 21.0 |
| 23 | 13 | 25 | 12 | 62.8 | 62.8 | 21.0 |
| 24 | 5 | 9 | 4 | 40.9 | 40.9 | 13.1 |

[a]Including the markers that were assigned to different (non-syntenic) LGs/chromosomes on the consensus map and on the reference genome assembly of *S. schlegelii*.

[b]Excluding the markers that were assigned to different (non-syntenic) LGs/chromosomes on the consensus map and on the reference genome assembly of *S. schlegelii.*

Abbreviation: LG, linkage group; Mb, mega base; #, number.

**TABLE S5** Markers showing significant single-locus transmission ratio distortion from the expected even ratio of homozygotes and heterozygotes. Both genotype frequency and the associated *p* value based on the chi-square test are shown in bold type if the *p* value after the false discovery rate correction is < 0.05. The marker name consists of a combination of the scaffold number to which the marker is mapped and the physical position (bp) on the scaffold, connected by an underscore.

| | | | | | Frequency of homozygotes | | | | Chi-square test *p* value after FDR correction | | | |
|---|---|---|---|---|---|---|---|---|---|---|---|---|
| | Position | | | | BC_Str | | BC_Ssc | | BC_Str | | BC_Ssc | |
| Marker | CHR[a] | Mb[a] | LG[b] | cM[b] | Larvae (n=48) | Subadults (n=87) | Larvae (n=48) | Subadults (n=87) | Larvae | Subadults | Larvae | Subadults |
| scaffold1_11977649* | 1 | 12.0 | NA | NA | NA | NA | **0.85** | **0.85** | NA | NA | **0.049** | **2.6.E-06** |
| scaffold1_21843854* | 1 | 21.8 | NA | NA | 0.79 | **0.7** | NA | NA | 0.13 | **0.017** | NA | NA |
| scaffold2_1581258 | 2 | 1.6 | 2 | 0.0 | 0.56 | **0.35** | NA | NA | 0.90 | **0.045** | NA | NA |
| scaffold2_6418196 | 2 | 6.4 | 2 | 1.5 | 0.58 | **0.34** | 0.6 | 0.49 | 0.69 | **0.035** | 0.87 | 0.93 |
| scaffold2_8260951 | 2 | 8.3 | 2 | 11.9 | 0.68 | **0.33** | 0.69 | 0.49 | 0.51 | **0.021** | 0.61 | 0.93 |
| scaffold2_9848810 | 2 | 9.8 | 2 | 13.9 | 0.6 | **0.33** | NA | NA | 0.59 | **0.028** | NA | NA |
| scaffold2_9848980 | 2 | 9.8 | 2 | 13.9 | 0.64 | **0.29** | 0.61 | 0.50 | 0.51 | **0.0083** | 0.81 | 1.00 |
| scaffold2_10090091 | 2 | 10.1 | 2 | 14.7 | 0.58 | **0.28** | 0.61 | 0.49 | 0.64 | **0.015** | 0.81 | 0.93 |
| scaffold2_11323203 | 2 | 11.3 | 2 | 16.2 | 0.5 | **0.33** | NA | NA | 1 | **0.019** | NA | NA |
| scaffold2_12342287 | 2 | 12.3 | 2 | 18.5 | 0.6 | **0.3** | 0.61 | 0.56 | 0.54 | **0.011** | 0.81 | 0.59 |
| scaffold2_13731483 | 2 | 13.7 | 2 | 22.3 | 0.54 | **0.32** | 0.6 | 0.47 | 0.84 | **0.015** | 0.81 | 0.79 |
| scaffold2_15281965 | 2 | 15.3 | 2 | 26.4 | 0.62 | **0.31** | 0.54 | 0.49 | 0.51 | **0.015** | 0.96 | 0.89 |
| scaffold2_16348796 | 2 | 16.3 | 2 | 26.4 | 0.64 | **0.32** | 0.56 | 0.52 | 0.51 | **0.018** | 0.96 | 0.79 |
| scaffold2_17251729 | 2 | 17.3 | 2 | 29.8 | 0.64 | **0.33** | 0.56 | 0.52 | 0.51 | **0.021** | 0.96 | 0.79 |
| scaffold2_17429664 | 2 | 17.4 | 2 | 29.1 | 0.62 | **0.31** | 0.5 | 0.54 | 0.51 | **0.015** | 1.0 | 0.68 |

| | | | | | | | | | | | | |
|---|---|---|---|---|---|---|---|---|---|---|---|---|
| scaffold2_19158518 | 2 | 19.2 | 2 | 30.6 | 0.62 | **0.33** | 0.5 | 0.6 | 0.51 | **0.019** | 1.0 | 0.46 |
| scaffold2_20983387 | 2 | 21.0 | 2 | 33.7 | 0.62 | **0.31** | 0.48 | 0.52 | 0.51 | **0.015** | 0.96 | 0.89 |
| scaffold2_20983545 | 2 | 21.0 | 2 | 34.1 | 0.62 | **0.3** | 0.53 | 0.56 | 0.51 | **0.014** | 0.96 | 0.46 |
| scaffold2_21570782 | 2 | 21.6 | 2 | 35.0 | 0.63 | **0.31** | 0.48 | 0.55 | 0.51 | **0.015** | 0.96 | 0.55 |
| scaffold2_21899912 | 2 | 21.9 | 2 | 35.0 | 0.62 | **0.27** | 0.5 | 0.56 | 0.51 | **0.018** | 1.0 | 0.51 |
| scaffold2_23033311 | 2 | 23.0 | 2 | 36.8 | 0.69 | **0.31** | 0.53 | 0.53 | 0.39 | **0.015** | 0.96 | 0.79 |
| scaffold2_23495478 | 2 | 23.5 | 2 | 36.8 | 0.64 | **0.32** | 0.52 | 0.59 | 0.51 | **0.015** | 0.96 | 0.41 |
| scaffold2_26166240 | 2 | 26.2 | 2 | 41.0 | 0.6 | **0.31** | 0.48 | 0.53 | 0.54 | **0.021** | 0.96 | 0.74 |
| scaffold2_27175102 | 2 | 27.2 | 2 | 42.4 | 0.61 | **0.3** | 0.48 | 0.58 | 0.54 | **0.015** | 0.96 | 0.43 |
| scaffold2_27277755 | 2 | 27.3 | 2 | 42.4 | 0.61 | **0.32** | 0.5 | 0.56 | 0.54 | **0.015** | 1.0 | 0.55 |
| scaffold2_28296210 | 2 | 28.3 | 2 | 42.5 | 0.62 | **0.33** | 0.49 | 0.55 | 0.51 | **0.024** | 0.96 | 0.55 |
| scaffold2_28933847 | 2 | 28.9 | 2 | 44.7 | 0.58 | **0.31** | 0.5 | 0.58 | 0.69 | **0.015** | 1.0 | 0.43 |
| scaffold2_30201033 | 2 | 30.2 | 2 | 48.6 | 0.6 | **0.31** | 0.51 | 0.57 | 0.59 | **0.015** | 0.96 | 0.46 |
| scaffold2_32110727 | 2 | 32.1 | 2 | 47.8 | 0.64 | **0.31** | 0.48 | 0.62 | 0.51 | **0.015** | 0.96 | 0.30 |
| scaffold2_33321169 | 2 | 33.3 | 2 | 48.6 | 0.59 | **0.31** | 0.5 | 0.65 | 0.64 | **0.015** | 1.0 | 0.13 |
| scaffold2_34391311 | 2 | 34.4 | 2 | 47.8 | 0.64 | **0.31** | 0.52 | 0.57 | 0.51 | **0.015** | 0.96 | 0.43 |
| scaffold2_35428472 | 2 | 35.4 | 2 | 47.8 | 0.59 | **0.3** | 0.5 | 0.58 | 0.75 | **0.015** | 1.0 | 0.43 |
| scaffold2_36727739 | 2 | 36.7 | 2 | 48.7 | 0.61 | **0.32** | 0.54 | 0.59 | 0.54 | **0.015** | 0.96 | 0.36 |
| scaffold2_41086016 | 2 | 41.1 | 2 | 47.8 | 0.57 | **0.31** | 0.51 | 0.57 | 0.69 | **0.015** | 0.96 | 0.41 |
| scaffold2_41791688 | 2 | 41.8 | 2 | 48.6 | 0.63 | **0.33** | 0.53 | 0.61 | 0.51 | **0.018** | 0.96 | 0.23 |
| scaffold4_5530907* | 4 | 5.5 | NA | NA | NA | NA | **0.84** | **0.74** | NA | NA | **0.0017** | **9.54E-04** |
| scaffold4_8712230 | 4 | 8.7 | 4 | 54.3 | NA | NA | 0.52 | **0.81** | NA | NA | 0.96 | **1.50E-04** |
| scaffold4_27762331* | 4 | 27.8 | NA | NA | NA | NA | 0.80 | **0.83** | NA | NA | 0.22 | **0.0018** |
| scaffold4_27763424 | 4 | 27.8 | 4 | 70.9 | NA | NA | 0.5 | **0.71** | NA | NA | 1.0 | **0.034** |
| scaffold5_22744824 | 5 | 22.7 | 5 | 45.9 | **0.81** | 0.55 | 0.66 | 0.58 | **0.025** | 0.64 | 0.71 | 0.46 |

| | | | | | | | | | | | | |
|---|---|---|---|---|---|---|---|---|---|---|---|---|
| scaffold6_5304596 | 6 | 5.3 | 6 | 30.9 | 0.67 | **0.77** | 0.54 | 0.57 | 0.47 | **0.018** | 0.96 | 0.43 |
| scaffold6_10274562 | 6 | 10.3 | 6 | 46.5 | 0.63 | **0.34** | NA | NA | 0.51 | **0.035** | NA | NA |
| scaffold6_13743546 | 6 | 13.7 | 6 | 51.8 | 0.61 | **0.34** | NA | NA | 0.54 | **0.035** | NA | NA |
| scaffold6_19228256 | 6 | 19.2 | 6 | 54.6 | 0.54 | **0.34** | 0.47 | 0.68 | 0.84 | **0.034** | 0.96 | 0.062 |
| scaffold6_20451351 | 6 | 20.5 | 6 | 55.5 | NA | NA | 0.55 | **0.68** | NA | NA | 0.96 | **0.024** |
| scaffold6_21438472 | 6 | 21.4 | 6 | 57.3 | 0.58 | **0.33** | 0.46 | 0.62 | 0.64 | **0.021** | 0.96 | 0.17 |
| scaffold6_22147786 | 6 | 22.1 | 6 | 54.0 | NA | NA | 0.55 | **0.68** | NA | NA | 0.96 | **0.036** |
| scaffold6_22390672 | 6 | 22.4 | 6 | 59.0 | 0.5 | **0.32** | 0.57 | 0.55 | 1 | **0.041** | 0.95 | 0.75 |
| scaffold6_25354904 | 6 | 25.4 | 6 | 60.8 | NA | NA | 0.52 | **0.66** | NA | NA | 0.96 | **0.043** |
| scaffold6_27417444 | 6 | 27.4 | 6 | 62.3 | NA | NA | 0.55 | **0.67** | NA | NA | 0.96 | **0.034** |
| scaffold6_27584088 | 6 | 27.6 | 6 | 62.0 | 0.44 | **0.33** | 0.51 | 0.65 | 0.84 | **0.036** | 0.96 | 0.061 |
| scaffold6_28465114 | 6 | 28.5 | 6 | 62.8 | 0.43 | 0.36 | 0.57 | **0.7** | 0.80 | 0.062 | 0.96 | **0.013** |
| scaffold6_31352569 | 6 | 31.4 | 6 | 63.5 | 0.52 | **0.33** | 0.52 | 0.65 | 0.93 | **0.045** | 0.96 | 0.061 |
| scaffold6_32751328 | 6 | 32.8 | 6 | 63.5 | 0.51 | **0.34** | 0.57 | 0.63 | 0.94 | **0.034** | 0.95 | 0.17 |
| scaffold6_33789300 | 6 | 33.8 | 6 | 63.5 | 0.44 | **0.33** | 0.53 | 0.64 | 0.84 | **0.021** | 0.96 | 0.10 |
| scaffold6_34983247 | 6 | 35.0 | 6 | 63.5 | 0.45 | **0.31** | NA | NA | 0.84 | **0.015** | NA | NA |
| scaffold6_36349294 | 6 | 36.3 | 6 | 65.0 | 0.45 | **0.33** | 0.55 | 0.62 | 0.88 | **0.021** | 0.96 | 0.17 |
| scaffold6_37022538 | 6 | 37.0 | 6 | 63.5 | 0.30 | **0.29** | **0.84** | **0.81** | 0.50 | **0.0083** | **0.0041** | **4.46E-06** |
| scaffold7_604409* | 7 | 0.6 | NA | NA | NA | NA | 0.53 | **0.67** | NA | NA | 0.96 | **0.036** |
| scaffold7_604563 | 7 | 0.6 | 10 | 0.0 | 0.52 | 0.42 | 0.47 | **0.67** | 0.93 | 0.64 | 0.96 | **0.041** |
| scaffold7_817258* | 7 | 0.8 | NA | NA | 0.54 | **0.88** | 0.49 | 0.51 | 0.84 | **7.83E-09** | 0.96 | 0.93 |
| scaffold7_16407349* | 7 | 16.4 | NA | NA | NA | NA | **0.82** | 0.65 | NA | NA | **0.0032** | 0.071 |
| scaffold7_17393580 | 7 | 17.4 | 7 | 12.0 | 0.33 | **0.35** | 0.54 | 0.45 | 0.47 | **0.042** | 0.96 | 0.60 |
| scaffold7_17988801 | 7 | 18.0 | 7 | 12.0 | 0.42 | **0.33** | 0.54 | 0.45 | 0.69 | **0.045** | 0.96 | 0.55 |
| scaffold7_18488819 | 7 | 18.5 | 7 | 12.0 | 0.35 | **0.34** | NA | NA | 0.50 | **0.035** | NA | NA |

| | | | | | | | | | | | | |
|---|---|---|---|---|---|---|---|---|---|---|---|---|
| scaffold7_18808783 | 7 | 18.8 | 7 | 12.0 | 0.4 | **0.35** | 0.55 | 0.51 | 0.59 | **0.042** | 0.96 | 0.93 |
| scaffold7_27405461* | 7 | 27.4 | NA | NA | NA | NA | 0.65 | **0.76** | NA | NA | 0.60 | **0.0017** |
| scaffold8_4697274 | 8 | 4.7 | 8 | 1.8 | 0.60 | 0.48 | 0.79 | **0.74** | 0.59 | 0.86 | 0.06 | **0.0028** |
| scaffold9_6853412 | 9 | 6.9 | 9 | 4.1 | 0.51 | 0.54 | 0.72 | **0.77** | 0.94 | 0.69 | 0.39 | **0.0010** |
| scaffold10_4730380 | 10 | 4.7 | 10 | 0.0 | NA | NA | 0.56 | **0.69** | NA | NA | 0.96 | **0.015** |
| scaffold10_5028859 | 10 | 5.0 | 10 | 1.2 | 0.5 | 0.57 | 0.53 | **0.69** | 1 | 0.66 | 0.96 | **0.011** |
| scaffold10_5706527 | 10 | 5.7 | 10 | 1.2 | NA | NA | 0.58 | **0.70** | NA | NA | 0.95 | **0.0078** |
| scaffold10_5964461 | 10 | 6.0 | 10 | 0.1 | 0.5 | 0.51 | 0.56 | **0.69** | 1 | 0.93 | 0.96 | **0.011** |
| scaffold10_6220789* | 10 | 6.2 | NA | NA | NA | NA | 0.67 | **0.78** | NA | NA | 0.60 | **8.63E-05** |
| scaffold10_7800913 | 10 | 7.8 | 10 | 0.0 | NA | NA | 0.61 | **0.70** | NA | NA | 0.89 | **0.0069** |
| scaffold10_8131604 | 10 | 8.1 | 10 | 0.5 | 0.52 | 0.56 | 0.58 | **0.68** | 0.93 | 0.59 | 0.89 | **0.020** |
| scaffold10_8672346 | 10 | 8.7 | 10 | 1.2 | 0.55 | 0.59 | 0.58 | **0.72** | 0.84 | 0.34 | 0.91 | **0.0025** |
| scaffold10_11665757 | 10 | 11.7 | 10 | 3.4 | 0.55 | 0.56 | 0.54 | **0.73** | 0.84 | 0.69 | 0.96 | **0.0010** |
| scaffold10_13989197 | 10 | 14.0 | 10 | 3.4 | NA | NA | 0.67 | **0.71** | NA | NA | 0.76 | **0.0044** |
| scaffold10_15281490 | 10 | 15.3 | 10 | 5.9 | NA | NA | 0.48 | **0.72** | NA | NA | 0.96 | **0.0018** |
| scaffold10_17842647 | 10 | 17.8 | 10 | 8.1 | 0.55 | 0.53 | 0.49 | **0.75** | 0.84 | 0.77 | 0.96 | **0.0069** |
| scaffold10_18394075 | 10 | 18.4 | 10 | 9.8 | 0.52 | 0.51 | 0.53 | **0.73** | 0.93 | 0.93 | 0.96 | **0.0011** |
| scaffold10_19603171 | 10 | 19.6 | 10 | 10.7 | 0.55 | 0.51 | 0.51 | **0.81** | 0.84 | 0.91 | 0.96 | **1.47E-05** |
| scaffold10_19603551 | 10 | 19.6 | 10 | 9.8 | 0.54 | 0.51 | 0.49 | **0.74** | 0.84 | 0.91 | 0.96 | **9.09E-04** |
| scaffold10_20160394 | 10 | 20.2 | 10 | 11.9 | NA | NA | 0.49 | **0.76** | NA | NA | 0.96 | **1.50E-04** |
| scaffold10_20265391 | 10 | 20.3 | 10 | 11.9 | NA | NA | 0.48 | **0.74** | NA | NA | 0.96 | **7.96E-04** |
| scaffold10_22044498 | 10 | 22.0 | 10 | 13.0 | 0.70 | 0.49 | 0.36 | **0.73** | 0.44 | 0.91 | 0.80 | **0.0042** |
| scaffold10_22044683 | 10 | 22.0 | 10 | 13.0 | 0.64 | 0.50 | 0.45 | **0.74** | 0.51 | 1.00 | 0.96 | **0.0018** |
| scaffold10_23209355 | 10 | 23.2 | 10 | 14.4 | 0.60 | 0.49 | 0.45 | **0.75** | 0.59 | 0.91 | 0.96 | **0.0014** |
| scaffold10_24937527 | 10 | 24.9 | 10 | 20.5 | 0.62 | 0.52 | 0.5 | **0.68** | 0.51 | 0.86 | 1.0 | **0.022** |

| | | | | | | | | | | | | |
|---|---|---|---|---|---|---|---|---|---|---|---|---|
| scaffold10_24937619 | 10 | 24.9 | 10 | 20.3 | NA | NA | 0.52 | **0.68** | NA | NA | 0.96 | **0.024** |
| scaffold10_26523142 | 10 | 26.5 | 10 | 23.9 | NA | NA | 0.55 | **0.69** | NA | NA | 0.96 | **0.011** |
| scaffold10_28818570 | 10 | 28.8 | 10 | 33.7 | 0.62 | 0.51 | 0.52 | **0.67** | 0.54 | 0.91 | 0.96 | **0.027** |
| scaffold10_29538671 | 10 | 29.5 | 10 | 29.9 | 0.62 | 0.49 | 0.51 | **0.67** | 0.51 | 0.93 | 0.96 | **0.034** |
| scaffold12_16728946 | 12 | 16.7 | 12 | 15.0 | 0.51 | **0.35** | NA | NA | 0.94 | **0.042** | NA | NA |
| scaffold12_28068021 | 12 | 28.1 | 12 | 26.2 | 0.72 | 0.49 | **1.00** | **0.96** | 0.32 | 0.93 | **0.0017** | **5.30E-08** |
| scaffold13_529410* | 13 | 0.5 | NA | NA | NA | NA | **0.98** | **0.86** | NA | NA | **1.20E-07** | **5.30E-08** |
| scaffold13_2656770 | 13 | 2.7 | 13 | 1.1 | 0.68 | **0.79** | 0.69 | 0.67 | 0.50 | **0.0017** | 0.49 | 0.063 |
| scaffold13_13317110 | 13 | 13.3 | 13 | 38.8 | 0.54 | **0.65** | 0.53 | 0.43 | 0.84 | **0.042** | 0.96 | 0.55 |
| scaffold13_27481855 | 13 | 27.5 | 13 | 50.0 | 0.56 | **0.65** | 0.45 | 0.39 | 0.84 | **0.042** | 0.96 | 0.23 |
| scaffold13_27938903 | 13 | 27.9 | 13 | 50.0 | 0.52 | **0.65** | 0.49 | 0.4 | 0.93 | **0.042** | 0.96 | 0.24 |
| scaffold14_11493049 | 14 | 11.5 | 14 | 31.4 | 0.54 | **0.73** | NA | NA | 0.84 | **0.027** | NA | NA |
| scaffold14_17490768 | 14 | 17.5 | 14 | 40.0 | 0.51 | **0.68** | 0.6 | 0.54 | 0.94 | **0.015** | 0.87 | 0.63 |
| scaffold14_19668358 | 14 | 19.7 | 14 | 45.3 | 0.51 | **0.72** | 0.57 | 0.55 | 0.94 | **0.0083** | 0.91 | 0.59 |
| scaffold14_19668557 | 14 | 19.7 | 14 | 45.3 | 0.57 | **0.65** | 0.54 | 0.56 | 0.77 | **0.048** | 0.96 | 0.51 |
| scaffold14_19896971 | 14 | 19.9 | 14 | 45.9 | 0.53 | **0.68** | 0.57 | 0.59 | 0.90 | **0.015** | 0.91 | 0.31 |
| scaffold14_20678275 | 14 | 20.7 | 14 | 45.9 | 0.5 | **0.68** | 0.57 | 0.57 | 1 | **0.016** | 0.95 | 0.46 |
| scaffold14_20681350 | 14 | 20.7 | 14 | 45.9 | 0.48 | **0.67** | 0.59 | 0.53 | 0.93 | **0.021** | 0.91 | 0.84 |
| scaffold14_21341700 | 14 | 21.3 | 14 | 48.6 | 0.59 | **0.68** | 0.42 | 0.53 | 0.64 | **0.015** | 0.91 | 0.74 |
| scaffold14_21600321 | 14 | 21.6 | 14 | 45.9 | 0.55 | **0.68** | 0.56 | 0.56 | 0.84 | **0.015** | 0.95 | 0.55 |
| scaffold14_22755109 | 14 | 22.8 | 14 | 45.9 | 0.42 | **0.68** | 0.58 | 0.58 | 0.71 | **0.015** | 0.93 | 0.43 |
| scaffold14_25067613 | 14 | 25.1 | 14 | 46.0 | 0.54 | **0.67** | 0.57 | 0.56 | 0.84 | **0.045** | 0.95 | 0.46 |
| scaffold14_26786413 | 14 | 26.8 | 14 | 46.0 | 0.44 | **0.68** | NA | NA | 0.84 | **0.015** | NA | NA |
| scaffold14_26894274 | 14 | 26.9 | 14 | 50.3 | 0.48 | **0.71** | NA | NA | 0.94 | **0.015** | NA | NA |
| scaffold14_27139719 | 14 | 27.1 | 14 | 46.0 | 0.51 | **0.68** | 0.53 | 0.53 | 0.94 | **0.015** | 0.96 | 0.79 |

| | | | | | | | | | | | | |
|---|---|---|---|---|---|---|---|---|---|---|---|---|
| scaffold14_29638125 | 14 | 29.6 | 14 | 46.8 | 0.5 | **0.69** | NA | NA | 1 | **0.015** | NA | NA |
| scaffold14_31284624 | 14 | 31.3 | 14 | 46.0 | 0.47 | **0.73** | 0.54 | 0.52 | 0.90 | **0.016** | 0.96 | 0.84 |
| scaffold14_31284794 | 14 | 31.3 | 14 | 46.0 | 0.51 | **0.83** | 0.56 | 0.54 | 0.94 | **0.0083** | 0.95 | 0.63 |
| scaffold18_26504961 | 18 | 26.5 | 18 | 35.0 | 0.39 | **0.34** | 0.48 | 0.42 | 0.69 | **0.034** | 0.96 | 0.41 |
| scaffold19_6709982 | 19 | 6.7 | 19 | 27.4 | NA | NA | 0.47 | **0.34** | NA | NA | 0.96 | **0.043** |
| scaffold19_7911118 | 19 | 7.9 | 19 | 32.4 | NA | NA | 0.43 | **0.31** | NA | NA | 0.96 | **0.036** |
| scaffold19_21759817 | 19 | 21.8 | 19 | 52.5 | NA | NA | 0.36 | **0.33** | NA | NA | 0.71 | **0.041** |
| scaffold20_19207887 | 20 | 19.2 | 20 | 56.6 | 0.64 | **0.96** | 0.47 | 0.54 | 0.54 | **3.16E-07** | 0.96 | 0.68 |
| scaffold21_6890635 | 21 | 6.9 | 21 | 5.8 | 0.42 | **0.68** | NA | NA | 0.84 | **0.023** | NA | NA |
| scaffold21_15366255 | 21 | 15.4 | 21 | 13.7 | 0.46 | **0.69** | 0.68 | 0.69 | 0.94 | **0.021** | 0.71 | 0.061 |
| scaffold21_19168602 | 21 | 19.2 | 21 | 21.2 | 0.71 | **0.76** | 0.6 | 0.65 | 0.47 | **0.011** | 0.87 | 0.17 |
| scaffold22_2897784* | 22 | 2.9 | NA | NA | NA | NA | **0.23** | **0.3** | NA | NA | **0.036** | **0.041** |
| scaffold23_18365872 | 23 | 18.4 | 23 | 38.7 | 0.65 | 0.36 | 0.58 | **0.72** | 0.50 | 0.051 | 0.91 | **0.022** |
| scaffold23_19589860 | 23 | 19.6 | 23 | 41.0 | 0.68 | **0.33** | 0.58 | 0.65 | 0.46 | **0.021** | 0.89 | 0.078 |
| scaffold23_20391696 | 23 | 20.4 | 23 | 41.7 | 0.63 | **0.31** | 0.55 | 0.64 | 0.51 | **0.015** | 0.96 | 0.18 |
| scaffold23_23252174 | 23 | 23.3 | 23 | 50.1 | 0.49 | **0.28** | 0.60 | 0.56 | 0.94 | **0.0083** | 0.81 | 0.51 |

*Removed from dataset during the linkage map construction.

[a]Position on the reference genome assembly of *S. schlegelii*

[b]Position on the consensus map.

Abbreviation: CHR, chromosome; cM, centimorgan; FDR, false discovery rate; LG, linkage group; Mb, mega base.

**TABLE S6** Possible epistatic interactions between pairs of nuclear loci as inferred from non-random associations of genotypes between pairs of marker blocks on different LGs (chi-square test; nominal $p < 0.01$). Marker positions for a given pair of marker blocks are the ranges of positions of the markers constituting the marker block on each LG. Genotype frequencies for a given pair of marker blocks are the average over the individual pairs of markers involved: A and B represent alleles from *S. trivittatus* and *S. schlegelii*, respectively, with XX_YY representing homozygous for allele X on the First LG and homozygous for allele Y on the 2nd LG. Associations involving the markers assigned to different (non-syntenic) LGs/chromosomes on the consensus linkage map and the reference genome assembly of *S. schlegelii* are indicated by 'Y' in the 'Inconsistent marker' column.

| | | | | Marker positions[a] | | | | | | | | | | | | | |
|---|---|---|---|---|---|---|---|---|---|---|---|---|---|---|---|---|---|
| | | LG | | 1st LG (cM) | | 2nd LG (cM) | | 1st LG (kb) | | 2nd LG (kb) | | Genotype frequency[b] | | | | | |
| Family | Stage | 1st | 2nd | from | to | from | to | from | to | from | to | AA_AA / BB_BB | AA_AB / BB_AB | AB_AA / AB_BB | AB_AB | # marker pairs | Inconsistent marker[c] |
| BC_Str | Larva | 3 | 4 | 14.4 | 19.7 | 64.3 | 70.9 | 30,758 | 32,395 | 27,806 | 27,979 | 0.16 | 0.39 | 0.37 | 0.08 | 6 | N |
| BC_Str | Larva | 3 | 6 | 59.3 | 60.5 | 59.8 | 63.5 | 38,468 | 38,799 | 23,915 | 32,751 | 0.12 | 0.38 | 0.37 | 0.13 | 8 | N |
| BC_Str | Larva | 8 | 12 | 46.4 | 51.1 | 25.2 | 42.5 | 33,562 | 34,232 | 24,057 | 32,818 | 0.46 | 0.26 | 0.02 | 0.25 | 9 | Y |
| BC_Str | Larva | 9 | 23 | 24.1 | 31.3 | 38.7 | 41.0 | 24,769 | 30,783 | 18,366 | 19,590 | 0.49 | 0.08 | 0.16 | 0.27 | 4 | N |
| BC_Str | Larva | 11 | 20 | 0.0 | 8.8 | 2.3 | 11.5 | 3,797 | 6,069 | 708 | 2,716 | 0.33 | 0.09 | 0.16 | 0.41 | 7 | N |
| BC_Str | Larva | 12 | 18 | 0.0 | 1.5 | 0.0 | 3.6 | 723 | 6,352 | 3,023 | 4,742 | 0.33 | 0.17 | 0.08 | 0.42 | 3 | N |
| BC_Str | Subadult | 1 | 18 | 25.7 | 60.2 | 0.0 | 12.7 | 11,117 | 40,069 | 3,023 | 22,286 | 0.31 | 0.26 | 0.08 | 0.34 | 16 | N |
| BC_Str | Subadult | 1 | 21 | 40.8 | 43.9 | 25.7 | 40.6 | 27,293 | 27,607 | 20,321 | 24,212 | 0.22 | 0.33 | 0.33 | 0.11 | 8 | N |
| BC_Str | Subadult | 3 | 16 | 19.7 | 34.2 | 11.2 | 60.2 | 32,395 | 35,450 | 8,017 | 24,970 | 0.31 | 0.12 | 0.20 | 0.37 | 5 | N |
| BC_Str | Subadult | 7 | 9 | 4.8 | 8.9 | 14.6 | 17.0 | 12,155 | 15,513 | 16,157 | 21,250 | 0.12 | 0.25 | 0.42 | 0.21 | 3 | N |
| BC_Str | Subadult | 10 | 18 | 1.2 | 3.4 | 12.7 | 16.6 | 8,672 | 11,666 | 22,286 | 23,643 | 0.33 | 0.26 | 0.07 | 0.35 | 3 | N |
| BC_Str | Subadult | 10 | 21 | 8.1 | 14.4 | 11.4 | 13.7 | 17,843 | 23,209 | 14,910 | 16,751 | 0.22 | 0.29 | 0.37 | 0.11 | 6 | N |
| BC_Str | Subadult | 11 | 20 | 0.0 | 12.6 | 54.5 | 62.0 | 3,797 | 7,249 | 18,289 | 27,555 | 0.10 | 0.30 | 0.37 | 0.23 | 8 | N |

| | | | | | | | | | | | | | | | | | |
|---|---|---|---|---|---|---|---|---|---|---|---|---|---|---|---|---|---|
| BC_Str | Subadult | 12 | 18 | 25.2 | 35.6 | 8.8 | 35.0 | 26,096 | 32,818 | 19,356 | 26,505 | 0.24 | 0.16 | 0.14 | 0.46 | 6 | N |
| BC_Str | Subadult | 19 | 24 | 0.0 | 1.3 | 39.6 | 40.9 | 161 | 338 | 12,907 | 13,617 | 0.12 | 0.32 | 0.35 | 0.21 | 3 | N |
| BC_Str | Subadult | 20 | 22 | 6.9 | 10.0 | 55.6 | 60.5 | 2,095 | 2,546 | 22,629 | 24,653 | 0.30 | 0.19 | 0.14 | 0.37 | 5 | Y |
| BC_Str | Subadult | 22 | 23 | 43.9 | 47.8 | 0.0 | 5.5 | 14,507 | 16,861 | 2,232 | 8,359 | 0.12 | 0.33 | 0.36 | 0.20 | 5 | N |
| BC_Ssc | Larva | 1 | 23 | 0.0 | 4.2 | 8.2 | 10.1 | 3,015 | 3,735 | 11,195 | 12,787 | 0.41 | 0.06 | 0.20 | 0.32 | 4 | N |
| BC_Ssc | Larva | 2 | 20 | 29.1 | 36.8 | 5.3 | 11.5 | 17,430 | 23,033 | 1,921 | 2,716 | 0.47 | 0.06 | 0.22 | 0.26 | 12 | N |
| BC_Ssc | Larva | 3 | 6 | 51.5 | 59.3 | 51.6 | 56.3 | 38,093 | 38,468 | 14,998 | 20,247 | 0.17 | 0.40 | 0.34 | 0.08 | 5 | N |
| BC_Ssc | Larva | 3 | 12 | 59.3 | 60.5 | 2.7 | 5.4 | 38,468 | 40,095 | 6,330 | 11,283 | 0.19 | 0.41 | 0.31 | 0.09 | 3 | N |
| BC_Ssc | Larva | 3 | 17 | 0.0 | 8.2 | 36.3 | 63.3 | 13,544 | 25,643 | 8,189 | 20,625 | 0.11 | 0.37 | 0.37 | 0.15 | 23 | N |
| BC_Ssc | Larva | 3 | 17 | 13.1 | 19.7 | 36.3 | 63.3 | 30,320 | 32,395 | 8,189 | 20,625 | 0.11 | 0.38 | 0.36 | 0.15 | 25 | N |
| BC_Ssc | Larva | 3 | 21 | 6.2 | 11.6 | 56.3 | 60.4 | 24,498 | 28,771 | 27,430 | 27,639 | 0.09 | 0.40 | 0.36 | 0.16 | 6 | N |
| BC_Ssc | Larva | 4 | 13 | 59.9 | 61.1 | 23.4 | 30.3 | 21,685 | 24,818 | 7,819 | 9,267 | 0.13 | 0.35 | 0.38 | 0.15 | 3 | N |
| BC_Ssc | Larva | 4 | 14 | 76.7 | 78.9 | 40.0 | 44.5 | 29,131 | 30,522 | 17,491 | 21,994 | 0.42 | 0.11 | 0.14 | 0.33 | 8 | N |
| BC_Ssc | Larva | 5 | 24 | 47.4 | 67.7 | 39.6 | 40.9 | 26,273 | 38,636 | 12,907 | 13,617 | 0.14 | 0.36 | 0.38 | 0.11 | 5 | Y |
| BC_Ssc | Larva | 6 | 12 | 38.0 | 63.5 | 0.0 | 18.5 | 1,301 | 31,353 | 723 | 24,018 | 0.37 | 0.14 | 0.12 | 0.38 | 32 | N |
| BC_Ssc | Larva | 7 | 21 | 25.7 | 59.6 | 11.4 | 17.0 | 26,418 | 33,263 | 14,910 | 18,415 | 0.47 | 0.14 | 0.10 | 0.29 | 9 | N |
| BC_Ssc | Larva | 10 | 23 | 11.9 | 13.0 | 19.4 | 38.7 | 20,160 | 22,044 | 15,470 | 18,366 | 0.37 | 0.07 | 0.19 | 0.38 | 4 | N |
| BC_Ssc | Larva | 12 | 15 | 23.2 | 29.2 | 10.4 | 22.3 | 27,619 | 29,650 | 14,933 | 23,362 | 0.35 | 0.10 | 0.15 | 0.40 | 6 | N |
| BC_Ssc | Larva | 12 | 24 | 37.1 | 47.2 | 39.6 | 40.9 | 24,958 | 24,958 | 12,907 | 12,907 | 0.33 | 0.07 | 0.21 | 0.39 | 4 | N |
| BC_Ssc | Larva | 13 | 23 | 14.8 | 21.6 | 10.1 | 19.4 | 5,840 | 7,087 | 12,787 | 15,470 | 0.17 | 0.32 | 0.41 | 0.09 | 3 | N |
| BC_Ssc | Larva | 15 | 22 | 51.8 | 54.0 | 47.8 | 50.8 | NA | NA | NA | NA | 0.45 | 0.08 | 0.20 | 0.28 | 4 | Y |
| BC_Ssc | Larva | 19 | 23 | 0.8 | 15.2 | 10.1 | 22.4 | 612 | 4,624 | 12,787 | 16,371 | 0.15 | 0.36 | 0.39 | 0.10 | 8 | N |
| BC_Ssc | Subadult | 1 | 3 | 4.2 | 8.7 | 14.4 | 19.7 | 3,735 | 5,053 | 30,758 | 32,395 | 0.32 | 0.16 | 0.14 | 0.39 | 4 | N |
| BC_Ssc | Subadult | 1 | 6 | 4.2 | 12.2 | 6.4 | 24.8 | 3,735 | 5,516 | 1,778 | 3,977 | 0.13 | 0.33 | 0.38 | 0.15 | 5 | N |
| BC_Ssc | Subadult | 1 | 20 | 31.2 | 35.3 | 54.5 | 56.2 | 13,827 | 15,385 | 18,289 | 19,259 | 0.38 | 0.14 | 0.19 | 0.29 | 3 | N |

| | | | | | | | | | | | | | | | | | |
|---|---|---|---|---|---|---|---|---|---|---|---|---|---|---|---|---|---|
| BC_Ssc | Subadult | 2 | 9 | 22.3 | 36.8 | 0.0 | 3.6 | 13,731 | 23,033 | 835 | 6,125 | 0.22 | 0.27 | 0.40 | 0.10 | 9 | N |
| BC_Ssc | Subadult | 2 | 10 | 17.5 | 48.6 | 0.1 | 20.5 | 11,503 | 33,174 | 5,029 | 24,938 | 0.48 | 0.06 | 0.25 | 0.22 | 93 | N |
| BC_Ssc | Subadult | 3 | 9 | 3.8 | 17.2 | 40.5 | 57.8 | 19,988 | 31,532 | 33,909 | 36,501 | 0.32 | 0.12 | 0.21 | 0.36 | 14 | N |
| BC_Ssc | Subadult | 5 | 13 | 24.1 | 43.0 | 41.8 | 47.1 | 6,925 | 21,309 | 14,458 | 18,350 | 0.31 | 0.22 | 0.12 | 0.35 | 10 | N |
| BC_Ssc | Subadult | 5 | 21 | 49.6 | 50.6 | 38.8 | 56.3 | 26,273 | 38,636 | 23,538 | 27,430 | 0.22 | 0.32 | 0.35 | 0.11 | 7 | Y |
| BC_Ssc | Subadult | 6 | 20 | 24.8 | 51.6 | 46.0 | 64.7 | 3,977 | 14,998 | 12,366 | 29,631 | 0.23 | 0.35 | 0.32 | 0.11 | 42 | N |
| BC_Ssc | Subadult | 6 | 22 | 49.0 | 60.8 | 35.6 | 58.6 | 10,275 | 25,355 | 10,189 | 24,308 | 0.45 | 0.20 | 0.12 | 0.23 | 19 | N |
| BC_Ssc | Subadult | 8 | 18 | 13.7 | 16.7 | 57.6 | 61.8 | 21,766 | 24,587 | 29,233 | 30,369 | 0.36 | 0.21 | 0.13 | 0.30 | 3 | N |
| BC_Ssc | Subadult | 8 | 23 | 6.4 | 50.7 | 0.0 | 10.1 | 15,528 | 33,462 | 2,232 | 12,787 | 0.18 | 0.39 | 0.30 | 0.14 | 32 | N |
| BC_Ssc | Subadult | 9 | 17 | 40.5 | 57.8 | 19.3 | 45.2 | 33,909 | 36,501 | 6,410 | 10,594 | 0.38 | 0.16 | 0.15 | 0.31 | 11 | N |
| BC_Ssc | Subadult | 9 | 21 | 0.0 | 1.3 | 56.3 | 60.4 | 835 | 1,578 | 27,430 | 27,639 | 0.43 | 0.20 | 0.11 | 0.26 | 4 | N |
| BC_Ssc | Subadult | 10 | 12 | 11.9 | 48.6 | 0.0 | 5.4 | 20,160 | 32,356 | 723 | 11,283 | 0.47 | 0.23 | 0.09 | 0.21 | 17 | N |
| BC_Ssc | Subadult | 11 | 22 | 12.6 | 27.9 | 7.5 | 14.9 | 6,693 | 10,907 | 4,338 | 5,278 | 0.37 | 0.12 | 0.19 | 0.33 | 10 | N |
| BC_Ssc | Subadult | 13 | 17 | 35.8 | 41.8 | 10.7 | 28.5 | 11,582 | 14,458 | 5,550 | 6,883 | 0.32 | 0.12 | 0.21 | 0.35 | 6 | N |
| BC_Ssc | Subadult | 16 | 19 | 7.6 | 9.1 | 51.0 | 52.5 | 6,550 | 7,543 | 21,329 | 22,165 | 0.06 | 0.34 | 0.28 | 0.32 | 3 | N |
| BC_Ssc | Subadult | 21 | 22 | 5.8 | 14.8 | 35.6 | 57.1 | 8,151 | 17,017 | 10,189 | 23,435 | 0.21 | 0.30 | 0.37 | 0.12 | 12 | N |
| BC_Ssc | Subadult | 21 | 22 | 37.5 | 38.8 | 14.9 | 35.6 | 23,538 | 24,018 | 4,338 | 10,189 | 0.25 | 0.33 | 0.32 | 0.10 | 2 | N |

[a]Ranges of the positions of the markers constituting the marker blocks on the consensus map (cM) and on the reference genome assembly of S. schlegelii (kb).

[b]Average two-marker genotype frequency of markers involved in the interaction. A and B represents alleles from S. trivittatus and S. schlegelii, respectively, with XX_YY representing homozygous for allele X on the First LG and homozygous for allele Y on the 2nd LG.

[c]Whether the markers mapped to non-syntenic (inconsistent) LGs/chromosomes on the consensus map and on the reference genome of S. schlegelii were involved.

**TABLE S7** Pigmentation-related genes included in the 95% CIs of the QTLs of color-related traits. The genes discussed in the main text are shown in bold type.

| **Stripe along lateral line (BC_Ssc; CHR 1; 27292667–40631577 bp)** | | | | | | | | |
|---|---|---|---|---|---|---|---|---|
| *S. schlegelii* annotation | Pannzer 2 | | | blastp | | | | |
| ID | PPV | GO_ID | GO_term | RefSeq_ID | Gene_symbol | Entrez Gene_ID | Description | Organism |
| Ssc_10010037 | 0.60 | 48753 | pigment granule organization | XP_044075813.1 | bloc1s1 | 122887052 | biogenesis of lysosomal organelles complex-1, subunit 1 | *Siniperca chuatsi* |
| Ssc_10012719 | 0.47 | 48066 | developmental pigmentation | XP_037636662.1 | pax7a | 119494668 | paired box 7a | *Sebastes umbrosus* |
| Ssc_10012729 | 0.35 | 46148 | pigment biosynthetic process | XP_037640238.1 | tamm41 | 119496760 | TAM41 mitochondrial translocator assembly and maintenance homolog | *Sebastes umbrosus* |
| Ssc_10006988 | 0.36 | 46148 | pigment biosynthetic process | XP_037641106.1 | cers5 | 119497225 | ceramide synthase 5 | *Sebastes umbrosus* |
| **Ssc_10004626** | **0.39** | **43473** | **pigmentation** | **XP_037621902.1** | **asip1** | **119486107** | **agouti signaling protein 1** | ***Sebastes umbrosus*** |

**TABLE S7 (continued)**

| Dark bands on dorsum (BC_Ssc; LG 2; 11503180–41791688 bp) | | | | | | | | |
|---|---|---|---|---|---|---|---|---|
| *S. schlegelii* | | | | | | | | |
| annotation | Pannzer 2 | | | blastp | | | | |
| ID | PPV | GO_ID | GO_term | RefSeq_ID | Symbol | Entrez Gene_ID | Description | Organism |
| Ssc_10020201 | 0.34 | 46148 | pigment biosynthetic process | XP_037611203.1 | lpcat2 | 119479534 | lysophosphatidylcholine acyltransferase 2 | *Sebastes umbrosus* |
| Ssc_10020241 | 0.52 | 30318 | melanocyte differentiation | XP_037613508.1 | trpm7 | 119480919 | transient receptor potential cation channel, subfamily M, member 7 | *Sebastes umbrosus* |
| — | 0.50 | 6582 | melanin metabolic process | — | — | — | — | — |
| **Ssc_10005231** | **0.35** | **42470** | **melanosome** | **XP_037647613.1** | **slc24a5** | **119501369** | **solute carrier family 24 member 5** | ***Sebastes umbrosus*** |
| — | **0.65** | **97324** | **melanocyte migration** | — | — | — | — | — |
| — | **0.63** | **30318** | **melanocyte differentiation** | — | — | — | — | — |
| Ssc_10007153 | 0.39 | 42470 | melanosome | XP_037648247.1 | rab27a | 119501738 | RAB27A, member RAS oncogene family | *Sebastes umbrosus* |
| **Ssc_10018924** | **0.85** | **4980** | **melanocyte-stimulating hormone receptor activity** | **XP_037646789.1** | **mc1r** | **119500870** | **melanocortin 1 receptor** | ***Sebastes umbrosus*** |
| Ssc_10018991 | 0.39 | 32402 | melanosome transport | XP_031702313.1 | arl6 | 116382967 | ADP-ribosylation factor-like 6 | *Anarrhichthys ocellatus* |
| Ssc_10019041 | 0.47 | 43473 | pigmentation | XP_037613313.1 | tjp1a | 119480764 | tight junction protein 1a | *Sebastes umbrosus* |
| Ssc_10019081 | 0.56 | 43473 | pigmentation | XP_037604284.1 | ctr9 | 119475543 | CTR9 homolog, Paf1/RNA polymerase II complex component | *Sebastes umbrosus* |
| Ssc_10020460 | 0.63 | 8057 | eye pigment granule organization | XP_003456933.1 | NA | 100705898 | V-type proton ATPase subunit d 1 | *Oreochromis niloticus* |
| — | 0.62 | 3406 | retinal pigment epithelium development | — | — | — | — | — |
| — | 0.59 | 32438 | melanosome organization | — | — | — | — | — |
| Ssc_10020513 | 0.60 | 51877 | pigment granule aggregation in cell center | XP_037614338.1 | bbs4 | 119481468 | Bardet-Biedl syndrome 4 | *Sebastes umbrosus* |
| — | 0.58 | 32402 | melanosome transport | — | — | — | — | — |
| Ssc_10005813 | 0.85 | 1903232 | melanosome assembly | XP_005815677.1 | ap1s2 | 102234402 | adaptor related protein complex 1 subunit sigma 2 | *Xiphophorus maculatus* |
| Ssc_10005829 | 0.64 | 30318 | melanocyte differentiation | XP_037610623.1 | leo1 | 119479286 | LEO1 homolog, Paf1/RNA polymerase II complex component | *Sebastes umbrosus* |

**TABLE S7 (continued)**

| **Yellow pigmentation (BC_Str; LG 12; 722823–20247139 bp)** | | | | | | | | |
|---|---|---|---|---|---|---|---|---|
| *S. schlegelii* | | | | | | | | |
| annotation | Pannzer 2 | | | blastp | | | | |
| ID | PPV | GO_ID | GO_term | RefSeq_ID | Gene_symbol | Entrez Gene_ID | Description | Organism |
| Ssc_10012479 | 0.86 | 3406 | retinal pigment epithelium development | XP_037631136.1 | yap1 | 119491314 | Yes1 associated transcriptional regulator | *Sebastes umbrosus* |
| Ssc_10002085 | 0.34 | 46148 | pigment biosynthetic process | XP_037623425.1 | fa2h | 119486939 | fatty acid 2-hydroxylase | *Sebastes umbrosus* |
| **Ssc_10016373** | **0.66** | **51876** | **pigment granule dispersal** | **XP_037630364.1** | **kcnj13** | **119490898** | **potassium inwardly rectifying channel subfamily J member 13** | ***Sebastes umbrosus*** |
| **—** | **0.64** | **51877** | **pigment granule aggregation in cell center** | **—** | **—** | **—** | **—** | **—** |
| **—** | **0.63** | **97324** | **melanocyte migration** | **—** | **—** | **—** | **—** | **—** |
| **—** | **0.61** | **30318** | **melanocyte differentiation** | **—** | **—** | **—** | **—** | **—** |

**TABLE S7 (continued)**

| **Color cluster 1: PC1 (BC_Str; LG 12; 722823–28068021 bp)** | | | | | | | | |
|---|---|---|---|---|---|---|---|---|
| *S. schlegelii* | | | | | | | | |
| annotation | Pannzer 2 | | | blastp | | | | |
| ID | PPV | GO_ID | GO_term | RefSeq_ID | Gene_symbol | Entrez Gene_ID | Description | Organism |
| Ssc_10012479 | 0.86 | 3406 | retinal pigment epithelium development | XP_037631136.1 | yap1 | 119491314 | Yes1 associated transcriptional regulator | *Sebastes umbrosus* |
| Ssc_10002085 | 0.34 | 46148 | pigment biosynthetic process | XP_037623425.1 | fa2h | 119486939 | fatty acid 2-hydroxylase | *Sebastes umbrosus* |
| Ssc_10016518 | 0.40 | 32438 | melanosome organization | XP_037630394.1 | rab38a | 119490922 | RAB38a, member RAS oncogene family | *Sebastes umbrosus* |
| **Ssc_10016373** | **0.66** | **51876** | **pigment granule dispersal** | **XP_037630364.1** | **kcnj13** | **119490898** | **potassium inwardly rectifying channel subfamily J member 13** | ***Sebastes umbrosus*** |
| **—** | **0.64** | **51877** | **pigment granule aggregation in cell center** | **—** | **—** | **—** | **—** | **—** |
| **—** | **0.63** | **97324** | **melanocyte migration** | **—** | **—** | **—** | **—** | **—** |
| **—** | **0.61** | **30318** | **melanocyte differentiation** | **—** | **—** | **—** | **—** | **—** |
| **Ssc_10016521** | **0.83** | **33162** | **melanosome membrane** | **XP_037630485.1** | **tyr** | **119491001** | **tyrosinase** | ***Sebastes umbrosus*** |
| **—** | **0.83** | **42438** | **melanin biosynthetic process** | **—** | **—** | **—** | **—** | **—** |
| **—** | **0.63** | **30318** | **melanocyte differentiation** | **—** | **—** | **—** | **—** | **—** |
| **—** | **0.62** | **32438** | **melanosome organization** | **—** | **—** | **—** | **—** | **—** |
| Ssc_10000968 | 0.44 | 48066 | developmental pigmentation | XP_037630444.1 | dharma | 119490963 | dharma | *Sebastes umbrosus* |
| Ssc_10008107 | 0.61 | 97324 | melanocyte migration | XP_037632096.1 | bace2 | 119491904 | beta-secretase 2 | *Sebastes umbrosus* |
| Ssc_10000812 | 0.44 | 48066 | developmental pigmentation | XP_037630444.1 | dharma | 119490963 | dharma | *Sebastes umbrosus* |

*S. schlegelii* annotation: gene IDs as reported in the gene annotation of the reference genome of *S. schlegelii*.

Pannzer 2: results from the Gene Ontology (GO) prediction using Pannzer 2.

blastp: best-hit of the blastp search against protein sequences of teleost genes deposited in the NCBI RefSeq protein database.

PPV: Positive Predictive Value, an estimate for the reliability of the predicted GO, reported by Pannzer 2. Takes values between 0 and 1.

GO: Gene Ontology.

—: same as above
