## Supplemental text for "Genetic architectures of postmating isolation and morphology of two highly diverged rockfishes (genus *Sebastes*)"

### Appendix S1 Supplementary methods

#### Rearing conditions of mapping families

All fishes, including the parents and the grandparents of the mapping families, were accommodated in Mariculture Fisheries Research Institute, Hokkaido Research Organization, Muroran, Hokkaido, Japan. Approximately 6,000 individuals of BC_Str progeny were housed in a 1-ton (t) tank from parturition (21 June 2016) till the termination of the experiment. Approximately 5,000 individuals of BC_Ssc progeny were housed in a 1-t tank upon parturition (4 July 2016), of which ca. 1,000 were transferred to another 1-t tank at 37 days after parturition (d) to be kept until the termination. The difference in the rearing scheme between families was intended to alleviate relative overcrowding of BC_Ssc, which grows faster than BC_Str (Table S1; Figure S8). The tanks were supplied with ambient seawater. The water was artificially cooled from the parturition until 18 August 2016 to mitigate the high water temperature in the summer: the actual water temperatures during the experiment were 3.1–21.3 ºC. Both families were fed with rotifers, *Artemia* nauplii, or pellets (in this order) daily according to their age and the body size, following the standard aquaculture practices for *S .schlegelii* in Japan (Fisheries Research Agency, 2010).

#### Patternize analysis

Extraction of color patterns from fish images using the *patRegK* function with predefined K = 4–6 resulted in 2, 3, and 3 informative clusters, respectively: other uninformative clusters apparently represented background colors (Figure S3). We regarded that K = 5 provided finer resolution of color patterns than K = 4, whereas K > 5 does not lead to further improvement. Therefore, we used the three color clusters extracted with K = 5 in the QTL mapping. Principal components 1 and 2 based on these color clusters explained 4.9% and 2.2% for the 1st color cluster, 6.0% and 3.0% for the 2nd color cluster, and 7.5% and 3.1% for the 3rd color cluster, respectively.

#### Geometric morphometrics

Principal components 1 and 2 based on geometric morphometrics explained 26.5% and 14.4% of the total variation in overall body shape, respectively, being subjected to QTL mapping. Each succeeding PCs explained less than 10% of the total variation.

**ddRAD library preparation, sequencing, and filtering**

Total genomic DNA was extracted from tissues using the Favorgen Tissue Genomic DNA Extraction Mini Kit (Favorgen). The DNA concentration was determined using a Quantus Fluorometer (Promega). Each sample was digested with high-fidelity EcoRI (New England Biolabs) and BglII (Takara Bio) and ligated with sequencing adaptors using T4 DNA Ligase (Enzymatics) in NEB buffer 2.1. After purification with AMPure XP (Beckman Coulter), each ligated sample was amplified using KAPA HiFi HS ReadyMix (Kapa Biosystems) with indexed primers. The PCR cycle was: 95 ºC for 3 min, followed by 20 cycles of 98 ºC for 20 s, 65 ºC for 10 s, and 72 ºC for 30 s. The obtained PCR products were pooled in equimolar and size-selected for a target range of 320–450 bp after purification, resulting in the final library size of ca. 260–400 bp.

A total of 211,402,675 and 749,545,358 reads were obtained by sequencing the two libraries, one for BC_Str subadults and another for the remaining samples, with the 50 bp single-end mode of HiSeq 2500 and 150 bp paired-end mode of HiSeq X systems, respectively. The number of reads per sample (mean ± SD) was 2,202,111 ± 748,215 and 5,260,618 ± 1,662,858, respectively. The initial genotype calling using the 'gstacks' program in Stacks v2.5.4 resulted in 769,998 RAD loci. After performing the filtering step separately for the two families as described in the main text, 642 and 823 SNPs (markers) with average depths of 84.8 and 107.4 were retained for BC_Str and BC_Ssc, respectively.

**Linkage map construction**

The initial formation of linkage groups (LGs) with a minimum LOD score of 6 and a maximum recombination fraction (RF) of 0.35 resulted in 26 LGs in BC_Str with 29 markers being assigned to inconsistent (non-syntenic) LGs/chromosomes in the linkage map and the reference genome assembly of *S. schlegelii*, and 25 LGs in BC_Ssc with 32 such 'inconsistent' markers. We manually reassigned those 'inconsistent' markers to alternative LGs to resolve the inconsistency as long as doing so was supported by the data supported. In BC_Str, markers that mapped to chromosome (CHR) 16 of the reference genome formed two separate LGs in the initial linkage map. Nonetheless, the markers assigned separately to the two LGs showed moderate linkage, forming a single LG when a lower LOD score threshold of 3 was applied. Furthermore, these markers formed a single LG in the initial linkage map of the other family, BC_Ssc. Therefore, we manually reassigned these markers to a single LG in the map of BC_Str (Table S3: 'Integrated to LG 16 in BC_Str'). Similarly, markers that mapped to CHRs 24 and 4 of the reference genome were each split into two LGs in the initial maps of BC_Str and BC_Ssc, respectively. These markers were manually merged into a single LG in each family since the markers formed a single LG with lower LOD score thresholds of 5 and 4 in BC_Str and BC_Ssc, respectively. The remaining 'inconsistent' markers were a few markers on each CHR that were initially assigned to a different LG from the one to which the other majority of markers on the same CHR were assigned. We did not manually reassign these markers to other LGs since none of them showed signs of linkage with markers on other LGs (Figure S4). Instead, we either left such markers on the LGs they were initially assigned to (Table S3: 'Left unmodified') or discarded them ('Discarded'), depending on whether the initial assignments to LGs were concordant between the two families. For example, three markers that mapped to CHR 7 of the reference genome were initially assigned to a LG corresponding to CHR 5 in both families and thus left on that LG. The resulting numbers of LGs were 24 in both families, with some LGs consisting of two groups of markers that mapped to different CHRs of the reference genome (Figure S5).
